# Longitudinal single-cell transcriptional profiling identifies dynamic neurodegenerative mechanisms in Christianson Syndrome rat

**DOI:** 10.64898/2026.10.14.756427

**Authors:** Hasib A. Riaz, Smijin Karthully Soman, Jessica F. Devlin, Gajendra Kumar, YouJin Lee, Eric M. Morrow

## Abstract

Endolysosomal dysfunction is a central feature of many neurodegenerative disorders. Loss-of-function mutations in the endosomal Na+/H+ exchanger NHE6 cause lysosomal dysfunction and the early-onset neurodegenerative disorder Christianson syndrome (CS). CS shares pathological features with more common neurodegenerative diseases; however, the extent of mechanistic overlap remains unclear. Using single-nucleus RNA sequencing across the lifespan in a CS rat model, we define the temporal progression of disease at single-cell resolution in the hippocampus. Neurons exhibit early transcriptional signatures of mitochondrial dysfunction. We investigated mitochondrial function in vitro and ex vivo and discover progressive defects in mitochondrial quality control and bioenergetics in NHE6-null tissue. Furthermore, both transcriptomic and proteomic analyses reveal robust activation of innate immunity, including cGAS-STING signaling which localizes to microglia. Therefore, in CS rat brain, early neuronal endosomal and mitochondrial impairments are followed by pronounced microglial and astrocytic activation, as well as age-dependent decline in oligodendrocyte myelinating capacity. Cross species comparison reveals substantial overlap between single-cell analyses in CS rat and in human postmortem Alzheimer’s disease hippocampus. Together, our findings define the cellular trajectories of neurodegeneration in CS, and identify convergent endolysosomal, mitochondrial, and inflammatory mechanisms that support a broader role for endolysosomal dysfunction in Alzheimer’s disease pathogenesis.

## Introduction

The endolysosomal system is an intricate network crucial for maintaining cellular homeostasis. Impaired endolysosomal function is one of the earliest neuropathological hallmarks in some neurodegenerative diseases, including Alzheimer’s disease (AD) and Parkinson’s disease (PD)^1,2^. Mitochondrial dysfunction also plays a central role in neurodegeneration, contributing to bioenergetic failure, oxidative stress, and the accumulation of damaged organelles in aging and neurodegenerative diseases^3,4^. The endolysosomal system and mitochondria are functionally interconnected through membrane contact sites, lysosome-dependent mitophagy, and pH-dependent ionic signaling, suggesting that impaired endolysosome function disrupts this crosstalk and plays a critical role in mitochondrial dysfunction and neurodegeneration^5,6^.

Christianson Syndrome (CS) is an endolysosomal disorder characterized by both neurodevelopmental and neurodegenerative features and caused by loss-of-function (LoF) mutations in the endosomal Na+/H+ exchanger 6 (NHE6) encoded by the X-linked SLC9A6 gene^7–9^. CS is clinically characterized by postnatal microcephaly, intellectual disability, cerebellar atrophy, ataxia and epilepsy^10,11^. Diffuse neuronal loss and tau deposition have also been reported in postmortem studies of CS patients^12^. Overlapping mechanisms, such as endolysosome dysfunction, a strong glial response in disease, and tau deposition, suggest that studies of CS may be informative to mechanisms in more common neurodegenerative disease such as AD^7,13^.

NHE6 localizes to early, recycling and late endosomes, and LoF mutations in NHE6 lead to endosome lumen overacidification that is associated with decreased neurotrophin endosomal signaling, reduced neuronal arborization and circuit defects in hippocampus and neocortex^8^. We previously reported that NHE6 loss in hippocampal neurons impairs lysosomal function by disrupting endosome-lysosome fusion^14^. Significant increases in microglial and astrocyte activity have been reported in the aging NHE6^-/y^ mouse brain^15^. Cortical neurons derived from human induced pluripotent stem cells of NHE6^-/y^ patients show augmented levels of pathogenically phosphorylated and sarkosyl-insoluble tau^16^.

Rats are more genetically and physiologically similar to humans than mice, and under certain pathological conditions^17–19^, they exhibit endogenous deposition of tau and amyloid-β^20,21^. However, endogenous tau deposition is generally not observed in mouse models of neurodegenerative disorders, including AD. Our laboratory identified endogenous deposition of amyloid-β and pathogenically-phosphorylated tau in the CS rat model, a particularly significant finding as few non-transgenic animal models endogenously demonstrate amyloid-β and tau deposition, making the CS rat model potentially relevant to AD pathology^13^. Additionally, we also found profound microglial and astrocyte activity, accompanied by autophagic and lysosomal defects in the CS rat. However, the underlying mechanisms of CS mediated neurodegeneration remain undefined.

To investigate the dynamics and molecular mechanisms of endolysosomal dysfunction in the CS rat at the single cell level, we performed longitudinal single nucleus RNA sequencing (snRNAseq) analysis from hippocampal tissue at 1, 8.5 and 17 months. We sub-categorized nuclei into specific cell types and identified salient transcriptional signatures from neuronal and nonneuronal cell types. Importantly, in both excitatory and inhibitory neurons, we identified transcriptional signatures of early impairment of mitochondrial function and oxidative phosphorylation. We also validated the mitochondrial dysfunction through complementary in vitro and ex vivo analysis, demonstrating progressive bioenergetic failure, electron transport chain (ETC) impairments, and disrupted mitochondrial quality control. We also identified and defined “CS-enriched” microglial and astrocyte clusters with transcriptomic expression similar to the disease-associated microglia (DAM) and disease-associated astrocyte (DAA) states observed in AD. Moreover, mass spectrometry (MS) proteomic analyses from hippocampal tissue identified early activation of the innate immune cGAS-STING pathway, a pathway activated by endolysosomal and mitochondrial dysfunction. This snRNAseq atlas across the lifespan in CS rats defines the sequence and dynamics of impaired endolysosomal and mitochondrial function in a highly penetrant and robust genetic model with strong overlap with AD, thereby potentially informing endolysosomal mechanisms in more common neurodegenerative diseases.

## Results

### Single-nuclei transcriptome atlas of hippocampus from Nhe6^-/y^ rat brain across the lifespan

To investigate the impact of loss of NHE6 on specific cellular states in vivo and across the lifespan, we performed snRNAseq analysis on hippocampus tissue on control and NHE6^-/y^ male rats at 1, 8.5 and 17 months of age. After quality checks (see Methods, **Extended Data Fig.1-3**) we obtained a combined total of 151,299 nuclei (**Fig. 1a**). Specifically, we identified: 48,887 nuclei (26,030 control, 22,857 NHE6^-/y^) at 1 month; 43,173 nuclei (24,505 control, 18,668 NHE6^-/y^) at 8.5 months; and 59,239 nuclei (33,058 control, 26,181 NHE6^-/y^) at 17 months. The overall number of nuclei were consistent across replicates (**Extended Data Fig. 4a)**. Using the integrated dataset from all timepoints, we identified 32 distinct clusters and by utilizing the expression of canonical marker genes. We annotated these clusters into 6 major cell types: excitatory neurons (*Camk2a, Slc17a7*), inhibitory neurons (*Gad1, Gad2*), microglia (*C1qa, Arhgap15*), astrocytes (*Aqp4*, *Gja1*), oligodendrocytes (*Mbp, Cnp*) and oligodendrocytes precursor cells (*OPC; C1ql1, Pdgfra*). Additionally, we identified less abundant cell types: vascular and leptomeningeal cells (VLMC, *Slc6a13*) (**Fig. 1b**). We were unable to identify the cell type of one cluster, which we labeled as an ambiguous cluster (353 total nuclei), and excluded this cluster from downstream analyses. The compositions of these cell types were consistent across the replicates (**Extended Data Fig. 4b**). To assess the distribution of cell types across time points, we calculated normalized cell type proportions (**Fig. 1c**). We further confirmed the identity of cell types using additional established markers (**Fig. 1d**; **Extended Data Fig. 4c**) and confirmed cell types by using the Allen Cell Type Database (MapMyCells) nomenclature. For downstream analysis, we focused on six major cell types. Our results reveal that NHE6 was expressed in both neuronal and glial cells, with highest expression in neurons, particularly inhibitory neurons, and notably low expression in microglia (**Extended Data Fig. 4d**). We confirmed these findings through immunostaining of NHE6 with Gad1 (inhibitory neurons), Camk2a (excitatory neurons), Gfap1 (astrocytes) and IBA1 (microglia) (**Extended Data Fig. 5a**). To understand global transcriptional changes, we investigated gene expression changes between control and NHE6^-/y^ rats across the timepoints on all nuclei using Model based Analysis of Single cell Transcriptomics (MAST) analysis with a random effect for sample of origin. Our results identified 1,599 downregulated and 1,117 upregulated genes at 1 month, 1,234 downregulated and 1,415 upregulated genes at 8.5 months, and 3,072 downregulated and 1,931 upregulated genes at 17 months (adjusted pvalue < 0.05 and log2fc > 0.1) (**Extended Data Fig. 5b; Extended Data Table.1**).

**Fig. 1.**
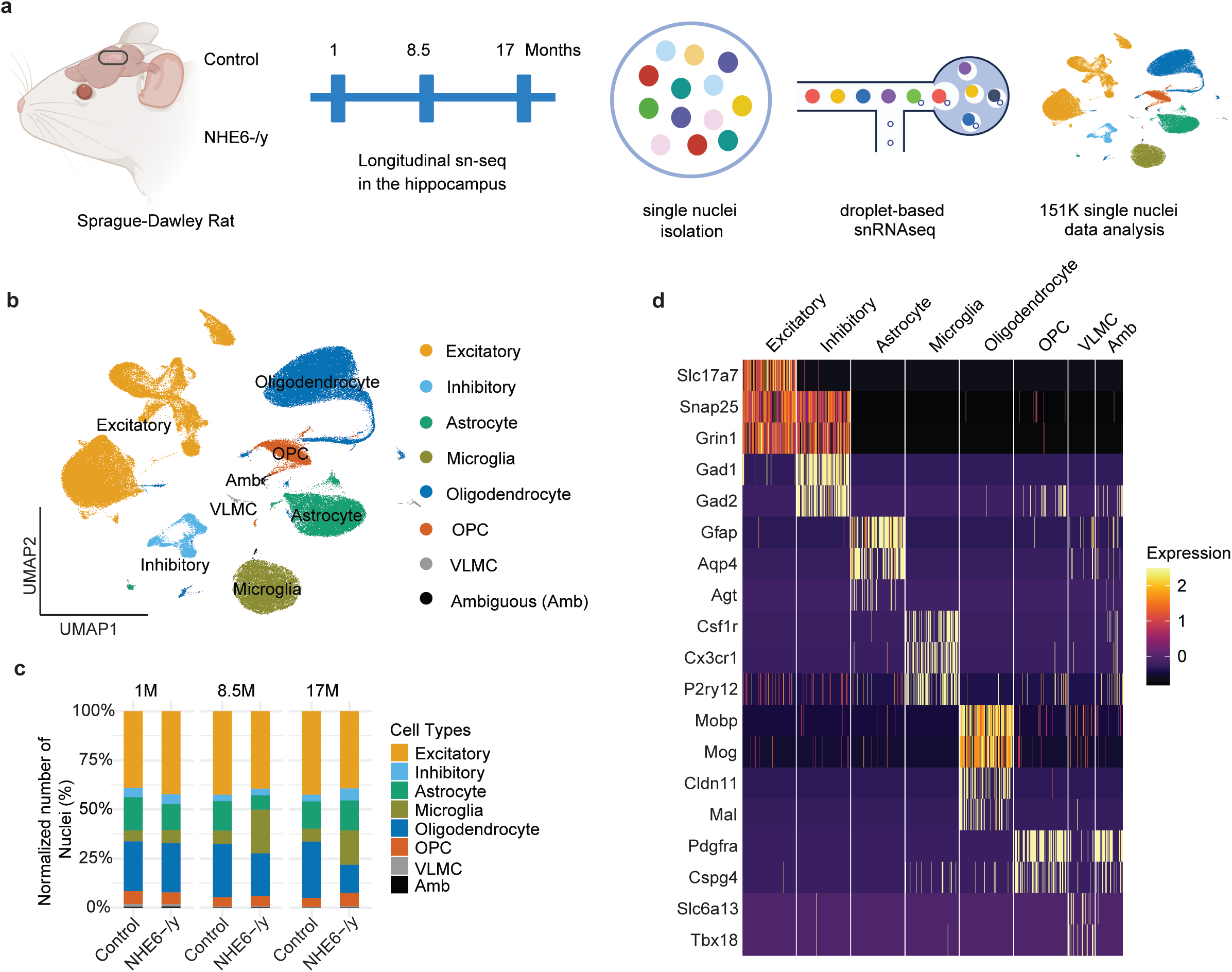
Single-nuclei transcriptome atlas of hippocampus from Nhe6-/y rat brain across the lifespan. **a,** Experimental workflow of snRNAseq study from dissection through analysis (created with BioRender). **b,** Uniform Manifold Approximation and Projection (UMAP) plot of snRNAseq of all 151,530 nuclei used for analysis. Clustering analysis revealed 7 distinct cell types. Oligodendrocyte precursor cells (OPC); Vascular leptomeningeal cells (VLMC). **c,** Bar chart indicating normalized percentage of cell types at each time point of the analysis. Number of nuclei were normalized with the total number of nuclei from each animal. **d,** Heatmap depicts the expression of cell type markers, a maximum of 100 nuclei per cluster are displayed.

### NHE6^-/y^ excitatory and inhibitory neurons demonstrate transcriptional evidence of early impairments in mitochondrial function

Hippocampal neurons play crucial roles in learning and memory and have been extensively studied in the context of neurodegenerative diseases for their involvement in cognitive function and impairment^22^. To address the role of hippocampal neurons in CS, we sub-categorized and analyzed neuronal nuclei into excitatory (62,466 total nuclei) and inhibitory neurons (6,626 total nuclei). The normalized nuclei count indicates that there was no significant change in the number of overall excitatory and inhibitory nuclei at all timepoints (**Extended Data Fig. 6a,b**). We performed differential expression analysis on the overall population of excitatory and inhibitory neurons. For excitatory neurons, we identified: 1,242 upregulated and 1,111 downregulated genes at 1 month; 225 upregulated and 835 downregulated genes at 8.5 months; and 653 upregulated and 968 downregulated genes at 17 months (with an adjusted p-value < 0.05 and log2FC > 0.1) **(Extended Data Fig. 6c, Extended Data Table.2)**. For inhibitory neurons, we identified: 174 upregulated and 258 downregulated genes at 1 month; 2 upregulated and 147 downregulated genes at 8.5 months; and 19 upregulated and 54 downregulated genes at 17 months (with an adjusted p-value < 0.05 and log2FC > 0.1) **(Extended Data Fig. 6d, Extended Data Table.3)**. Interestingly, the overall Gene Set Enrichment Analysis (GSEA) for both excitatory and inhibitory neurons with hallmark^23^ and KEGG^24^ datasets was similar, and indicated a strong effect on mitochondrial assembly complex and oxidative phosphorylation at 1 and 8.5 months (**Fig. 2a**).

**Fig. 2.**
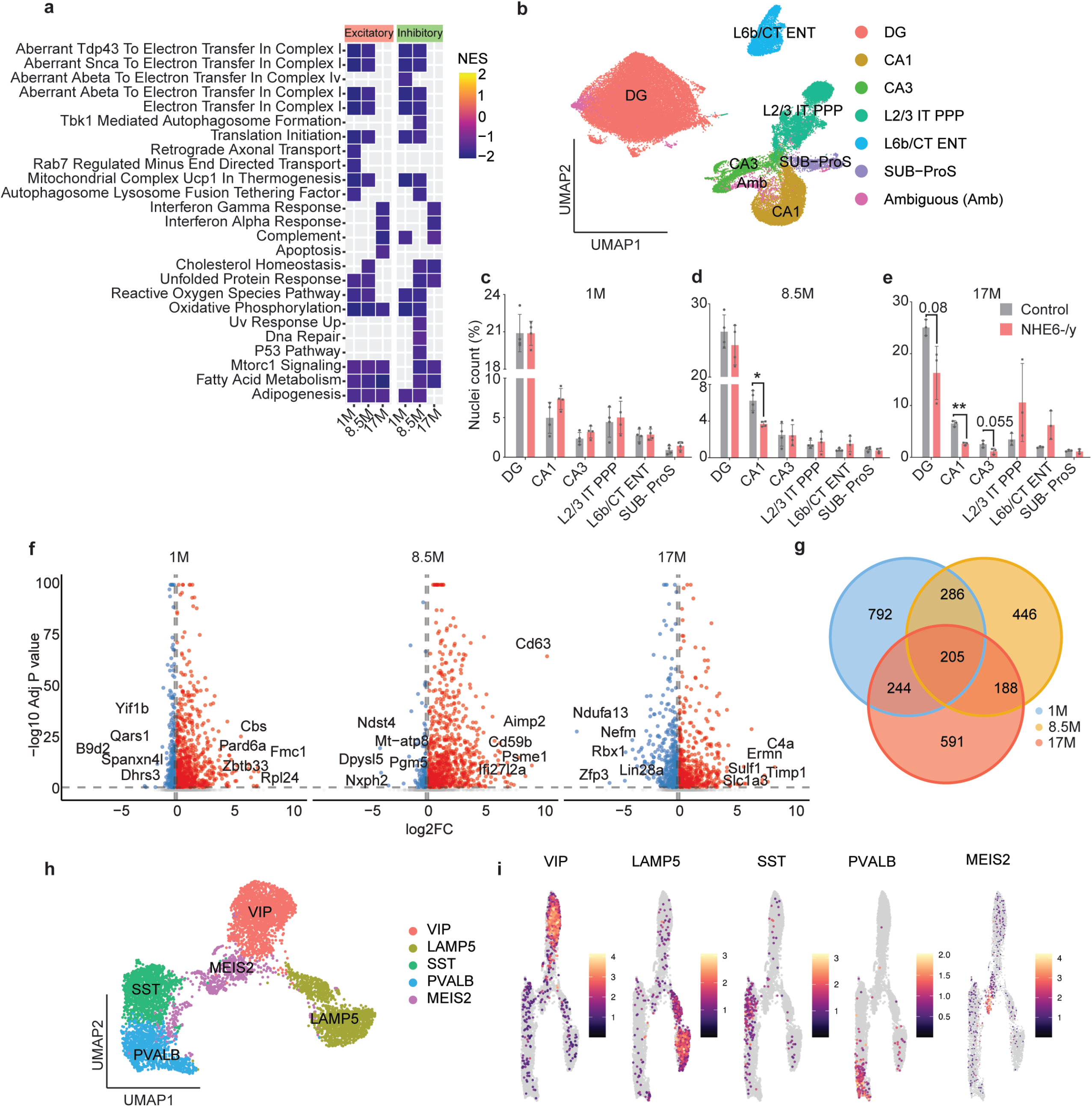
NHE6-/y excitatory and inhibitory neurons demonstrate transcriptional evidence of early impairments in mitochondrial function. **a,** Heatmap shows GSEA from hallmark and KEGG enriched pathways of both excitatory and inhibitory neurons across timepoints. Color shows normalized enrichment score (NES). **b,** UMAP plot shows subtypes of excitatory neuronal clusters from the rat hippocampal neurons labelled as DG, CA1, CA3, L2/3 IT PPP, L6b/CT ENT and SUB-ProS. **c-e,** Barplot indicates frequency of excitatory neurons cell type in subclusters at 1 **(c)** and 8.5 (**d)** and 17 **(e)** months in NHE6-/y rats. Number of nuclei in subtypes were normalized with the total number of nuclei form each animal. Data are presented as mean ± SEM. Each point in the graphs represents an individual brain. Two-tailed unpaired t-test with Welch’s correction was performed. *p-value, 0.014, **p-value, 0.002. **f,** Volcano plots exhibit the differentially expressed genes (DEGs) between control and NHE6-/y nuclei from dentate gyrus (DG) at all time points. Significant genes are labelled in color (Padj < 0.05, Log2FC > 0.1), while nonsignificant genes are shown in gray. Top 5 upregulated and downregulated genes are labelled. **g,** Venn diagrams show the shared DEGs genes in NHE6-/y rats from DG nuclei across time points. **h,** UMAP shows subtypes of inhibitory neuronal clusters in the rat hippocampus. **i,** Feature plot depicts expression of canonical marker genes of inhibitory subtype clusters; VIP, LAMP5, SST, PVALB and MEIS2.

To investigate excitatory neuron subtypes, we sub-clustered excitatory neurons and identified 15 distinct clusters as illustrated with the UMAP (**Extended Data Fig. 7a**). To confirm the identity of subclusters of excitatory neurons, we mapped subclusters to the Allen Cell Type Database (MapMyCells) and labelled clusters with their respective cell type identities. Our results show that excitatory neurons consist of 6 cell types Dentate Gyrus (DG) (34,511 total nuclei), Cornu Ammonis area 1 (CA1) (8216 total nuclei), Cornu Ammonis area 3 (CA3) (3,724 total nuclei), Layer 2/3 Intratelencephalic neurons, Parietal Posterior Parietal cortex (L2/3 IT PPP) (7,330 total nuclei), Layer 6b/ Corticothalamic neurons, Entorhinal cortex (L6b/CT ENT) (4,997 total nuclei) and Subiculum-Prosubiculum (SUB-ProS) (1,748 total nuclei) (**Fig. 2b**). It is important to note that we were unable to the identify cell type identity of one cluster and labeled it as an “Ambiguous” cluster (1,940 total nuclei) and subsequently excluded it from downstream analysis. To further confirm the identities of these subtypes, we used canonical marker gene expression, as shown in **Extended Data Fig. 7b**. Interestingly, the normalized nuclei count for these subtypes revealed no significant changes in nuclei numbers at 1 month (**Fig. 2c**). However, CA1 neurons were significantly reduced at both 8.5 and 17 months (**Fig. 2d**,**e**). DG (p-value 0.08) and CA3 (p-value 0.056) neurons were also reduced at 17 months. We did not observe any significant change in L2/3 IT PPP, L6b/CT ENT and SUB-ProS cell types at any timepoint. To validate the snRNAseq findings of progressive loss of CA1 and DG neurons, we stained control and NHE6^-/y^ rats with the NeuN antibody (neuronal marker) at all time points. As expected, the number of NeuN positive cells was similar in CA1 and DG area at 1 month (**Extended Data Fig. 7c**). Similar to snRNAseq data, the CA1 cells were significantly reduced at 8.5 and 17 months (**Extended Data Fig. 7d,e**). The differential gene expression analysis from DG neurons at all timepoints identified changes in mitochondrial genes (*Uqcrh, Ndufc1, Ndufb4, Ndufa13, Cox6c*) (**Fig. 2f, Extended Data Table.4**). The Gene Ontology (GO) analysis also indicated enriched pathways associated with dysfunctional mitochondria, axonal transport and amyloid beta binding (**Extended Data Fig. 8a**). To examine shared significant genes across all timepoints in DG neurons, we identified 205 genes that were significantly differentially expressed at each timepoint (**Fig. 2g, Extended Data Table.4)**. These genes were associated with mitochondrial function (*Ndufa13, Ndufb3, Uqcrh*) and amyloid-beta binding (*Gria1, Grin2A, Nlgn1, Pfdn2, Prnp*). The differential gene expression analysis from CA1 and CA3 neurons at all timepoints also identified DEGs between control and NHE6^-/y^ neurons at all timepoints (**Extended Data Fig. 8b,c, Extended Data Table.5**). It is important to note that the number of differentially expressed genes (DEGs) in these subtypes were higher at 1 month as compared 8.5 and 17 months (**Extended Data Table.5)**.

In addition to excitatory neurons, we also analyzed subclasses of inhibitory neurons, comprising a total of 6,626 nuclei. We sub-clustered inhibitory nuclei and identified 5 distinct clusters (**Extended Data Fig. 9a**). The Allen Cell Type Database (MapMyCells) identified 5 subtypes of inhibitory neurons as indicated by UMAP plot (**Fig. 2h**). The cell type identity of subtypes, namely Vasoactive Intestinal Polypeptide (VIP) (1,860 total nuclei), Lysosome-Associated Membrane Protein 5 (LAMP5) (1,501 total nuclei), Somatostatin (SST) (1,399 total nuclei), Parvalbumin (PVALB) (1,162 total nuclei) and Meis Homeobox 2 (Meis2) (704 total nuclei), were also confirmed through the expression of specific marker genes through FeaturePlot and heatmaps (**Fig. 2i, Extended Data Fig. 9b**). The normalized nuclei count for these inhibitory neuron subtypes revealed no significant changes in nuclei numbers at 1, 8.5 and 17 months (**Extended Data Fig. 9c-e**). We have further analyzed DEGs between control and NHE6^-/y^ inhibitory neuron subtypes but only few genes were significantly differentially expressed (**Extended Data Fig.9 f-j**). One possible explanation for the reduced number of DEGs is due to the low number of inhibitory neurons as compared with other cell types. Together these data suggest that NHE6^-/y^ rats exhibit transcriptomic evidence of mitochondrial and metabolic changes in both excitatory and inhibitory neurons. Notably, there are defects in retrograde axonal transport specifically in excitatory neurons early, at the 1 month timepoint only. This result is consistent with our prior data of the function of NHE6 in axonal transport^25^, and early failures in retrograde transport may secondarily lead to impaired mitophagy and mitochondrial health.

### Mitochondrial dysfunction in NHE6^-/y^ hippocampal neurons in vitro and ex vivo

Since the snRNAseq analyses reveal early mitochondrial dysfunction in neurons that may worsen with age, we further investigated mitochondrial impairments using both in vitro and ex vivo experiments. We hypothesized that mitochondrial membrane potential (Δψm) would be reduced in NHE6^-/y^ neurons and examined this using live cell TMRM fluorescence imaging in control and NHE6^-/y^ primary hippocampal neurons at 15 days in vitro (DIV). Under baseline conditions, NHE6^-/y^ neurons exhibited significantly lower steady state Δψm compared to controls **(Fig. 3a)**. To assess mitochondrial response to maximal uncoupling, FCCP was applied to induce maximal depolarization **(Extended Data Fig. 10a).** Both conditions underwent rapid depolarization; however, NHE6^-/y^ neurons displayed a reduced absolute loss of TMRM fluorescence relative to control, consistent with their lower initial polarization. Taken together, these findings demonstrate a global reduction in mitochondrial membrane potential in NHE6^-/y^ neurons. Δψm is a real time readout of mitochondrial health and essential for ATP generation, mitochondrial morphology, dynamics, Ca^2+^ sequestration and quality control^26^. We, therefore, assessed mitochondrial morphology in primary hippocampal neurons and observed no significant differences between control and NHE6^-/y^ neurons in somatic mitochondrial volume, number, and dendritic mitochondrial content at DIV-15 **(Extended Data Fig. 10b,c)**. Since adequate Δψm is vital for ATP synthesis, we hypothesized that there may be a functional bioenergetics deficit and performed Seahorse XF mitochondrial stress assays in DIV-15 primary hippocampal neurons. Oxygen consumption rate (OCR) was measured under basal conditions and following sequential injection of oligomycin, FCCP, and antimycin A/rotenone. NHE6^-/y^ neurons showed no significant differences from controls in basal respiration, ATP-linked respiration, proton leak, maximal respiratory capacity, or spare respiratory capacity at DIV-15 **(Extended Data Fig. 10d)**, indicating that steady-state mitochondrial respiration is broadly preserved in NHE6^-/y^ neurons at this age of cultures.

**Fig 3.**
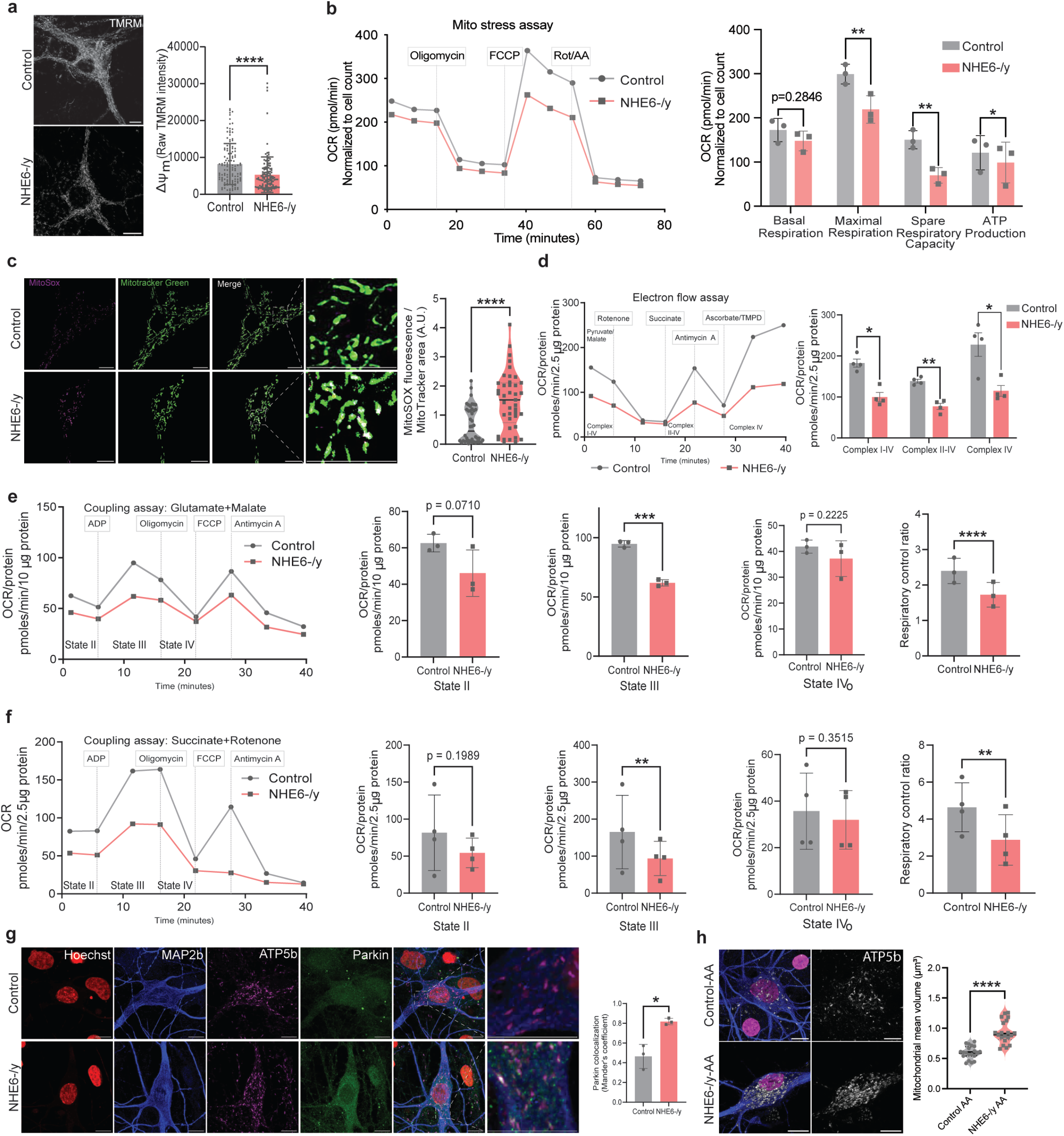
Mitochondrial dysfunction in NHE6-/y hippocampal neurons in vitro and ex vivo. **a,** Representative TMRM images of DIV-15 control and NHE6-/y primary hippocampal neurons show reduced mitochondrial membrane potential (ΔΨm) in NHE6-/y neurons. Data are presented as mean ± SEM; Each data point represents an individual neuron; n = 131 (Control) and 132 (NHE6-/y) neurons from 3 animals per genotype. Two-tailed unpaired t-test with Welch’s correction were performed. ****p-value 0.0001. Scale bar = 10 µm. **b,** Representative Seahorse XF in vitro Mito Stress Test traces showing oxygen consumption rate (OCR) in DIV-21 control and NHE6-/y hippocampal neurons. OCR was measured under basal conditions and following sequential injection of oligomycin, FCCP, and rotenone/antimycin A (Rot/AA). Quantification of basal respiration, maximal respiration, spare respiratory capacity and ATP production revealed significantly reduced OCR in NHE6-/y neurons compared to controls. OCR values were normalized to cell count. Data are presented as mean ± SEM. Each data point represents an individual brain (n = 3 per genotype). Two-tailed unpaired t-test with Welch’s correction was performed. *p-value 0.03, **p-value 0.006 (Maximal Respiration), **p-value 0.008 (Spare Respiratory Capacity). **c,** Representative images of MitoSOX (magenta) and MitoTracker Green (green) in DIV-21 control and NHE6-/y hippocampal neurons show elevated mitochondrial superoxide in NHE6-/y neurons. Quantification confirmed a significant increase in mitochondrial ROS production. Data are presented as mean ± SEM. Each data point represents an individual neuron; n = 41 (Control) and 41 (NHE6-/y) neurons from 3 animals per genotype. Two-tailed unpaired t-test with Welch’s correction were performed. ****p-value < 0.0001. Scale bar = 10 µm. **d,** Representative Seahorse XF ex vivo electron flow assay traces showing OCR in control and NHE6-/y P30 rat hippocampus mitochondria following sequential injection of rotenone, succinate, antimycin A, and ascorbate/TMPD to assess activities of mitochondrial electron transport chain complexes. Quantification revealed significant reductions in Complex I-IV, Complex II-IV, and Complex IV driven respiration in NHE6-/y neurons compared to controls. OCR values were normalized to protein concentration (2.5 µg protein). Data are presented as mean ± SEM. Each data point represents an individual brain (n = 3 per genotype). Two-tailed paired t-test was performed. *p-value 0.021 (Complex I-IV), **p-value 0.0074, *p-value 0.027 (Complex IV). **e, f,** Representative Seahorse XF ex vivo coupling assay traces showing OCR in control and NHE6-/y P30 rat hippocampus mitochondria supplemented with glutamate+malate **(e)** or succinate+rotenone **(f)** substrates. OCR was measured following sequential injection of ADP, oligomycin, FCCP, and antimycin A to assess mitochondrial respiratory states. Quantification demonstrated significantly reduced State III respiration in NHE6-/y neurons under both substrate conditions, while State IV₀ respiration was not significantly altered. Respiratory control ratio (RCR) was significantly decreased in NHE6-/y neurons, indicating impaired mitochondrial coupling efficiency. OCR values were normalized to protein concentration (10 µg protein in f; 2.5 µg protein in g). Data are presented as mean ± SEM. Each data point represents an individual brain (n = 3 per genotype). Two-tailed paired t-test was performed. ***p-value 0.001(Coupling assay: Glutamate+Malate: StateIII), ****p-value 0.0001 (Coupling assay: Glutamate+Malate: Respiratory control ratio), **p-value 0.005 (Coupling assay: Succinate+Rotenone: StateIII), **p-value 0.002 (Coupling assay: Succinate+Rotenone: Respiratory control ratio). **g,** Representative confocal images of DIV-21 control and NHE6-/y primary hippocampal neurons immunostained for Parkin (green), the mitochondrial marker ATP5b (magenta), MAP2b (blue), and DAPI (red). Merged and magnified images demonstrate increased colocalization of Parkin with mitochondria in NHE6-/y neurons compared to controls. Quantification of Manders coefficient revealed significantly increased Parkin-mitochondria colocalization in NHE6-/y neurons. Data are presented as mean ± SEM (n = 3 biological replicates). Each data point represents an individual experiment. Two-tailed unpaired t-test with Welch’s correction was performed. *p-value 0.03. Scale bar, 10 µm. **h,** Representative confocal images show mitochondrial morphology in DIV-21 control and NHE6-/y primary hippocampal neurons following AA treatment. Mitochondria are shown in grey and neuronal processes in blue. Quantification of mean mitochondrial volume revealed a significant increase in mitochondrial volume following AA treatment in NHE6-/y neurons compared to untreated NHE6-/y neurons and controls. Data are presented as mean ± SEM (n = 3 biological replicates). Each data point represents an individual experiment. Two-tailed unpaired t-test with Welch’s correction was performed. ****p-value 0.0001. Scale bar, 10 µm.

In our prior studies, we discovered that NHE6^-/y^ hippocampal neurons demonstrated worsening endolysosome dysfunction with age in culture relative to control^14^, and thereby, we reasoned that broader mitochondrial dysfunction may manifest in older cultures and animals. We re-assessed mitochondrial bioenergetic vulnerability at DIV-21 again using Seahorse XF mitochondrial stress assays in primary hippocampal neurons. Basal respiration was not significantly different between control and NHE6^-/y^ neurons, indicating that steady-state mitochondrial activity remains largely intact. However, by contrast to DIV-15, FCCP-induced maximal respiration showed a significant reduction in NHE6^-/y^ neurons compared to controls at DIV-21 (**Fig. 3b)**.

Spare respiratory capacity was also significantly reduced in NHE6^-/y^ neurons at DIV-21. This reduction reflects a diminished ability of mitochondria to respond to increased energetic demand despite relatively preserved basal OCR values. ATP-linked respiration, calculated as the oligomycin-sensitive fraction of basal OCR, was also significantly reduced in NHE6-/y neurons compared to control. To further assess mitochondrial health, we next assessed mitochondrial oxidative stress at DIV-21 by staining with MitoSOX, a mitochondria-targeted fluorescent indicator of superoxide production. Under basal conditions, NHE6^-/y^ neurons exhibited a significant elevation in MitoSOX integrated fluorescence normalized to MitoTracker Green intensity compared to control (**Fig. 3c),** indicating increased mitochondrial superoxide production independent of differences in mitochondrial mass.

To determine whether mitochondrial dysfunction observed in primary hippocampal neurons reflects intrinsic defects in oxidative phosphorylation in vivo, we performed ex vivo Seahorse-based analyses on fresh mitochondria isolated directly from 1 month old rat hippocampus. We first assessed ETC activity using electron flow assays. Under NADH-linked conditions (pyruvate/malate; Complex I-IV-driven), NHE6^-/y^ mitochondria exhibited a significant reduction in OCR compared to control, indicating impaired Complex I-dependent respiration (**Fig. 3d)**. This deficit persisted following rotenone-mediated inhibition of Complex I and stimulation of Complex II-IV respiration with succinate, where OCR remained significantly reduced in NHE6^-/y^ mitochondria. Furthermore, Complex IV-driven respiration (ascorbate/TMPD) was also significantly diminished in NHE6^-/y^ mitochondria, demonstrating compromised terminal electron transfer. Collectively, these findings indicate a broad impairment in ETC function in NHE6^-/y^ mitochondria freshly harvested from brain. These findings combined with the snRNAseq data support the presence of mitochondrial dysfunction in NHE6^-/y^ brain in vivo.

To assess the efficiency of oxidative phosphorylation further, we performed mitochondrial coupling assays under both NADH-linked (glutamate/malate) and FADH2-linked (succinate/rotenone) conditions on fresh mitochondria isolated directly from 1 month old rat hippocampus (**Fig. 3e,f)**. Under glutamate/malate-driven respiration, NHE6^-/y^ mitochondria exhibited significantly reduced State II respiration and a marked decrease in ADP-stimulated State III respiration, indicating impaired substrate utilization and ATP-generating capacity. In contrast, oligomycin-induced State IVₒ respiration was not significantly altered, suggesting that proton leak is not substantially affected (**Fig. 3e)**. Consequently, the respiratory control ratio (RCR) was significantly reduced in NHE6^-/y^ mitochondria, indicative of impaired coupling efficiency. A similar pattern was observed under succinate/rotenone-driven conditions, where NHE6^-/y^ mitochondria showed reduced State II and State III respiration, with no significant change in State IVₒ respiration (**Fig. 3f)**. Consistent with these findings, RCR was again significantly reduced in NHE6^-/y^ mitochondria, confirming a substrate-independent defect in coupling efficiency. Taken together, these results demonstrate that NHE6 deficiency impairs electron transport chain activity and the coupling of ETC to ATP synthesis in ex vivo hippocampal mitochondria.

Since NHE6 is an endosomal protein, and NHE6^-/y^ neurons in vitro and in vivo demonstrate evidence of worsening lysosome function with age^13,14^. We hypothesized that mitochondrial defects may be secondary to abnormal endolysosomal function. We therefore examined mitochondrial quality control by assessing Parkin localization, a key mediator of PINK1-dependent mitophagy, in DIV-21 primary hippocampal neurons. In control neurons, Parkin exhibited a predominantly diffuse cytosolic distribution with minimal co-localization with the mitochondrial marker ATP5B (**Fig. 3g)**. In contrast, NHE6^-/y^ neurons markedly increased Parkin recruitment to the outer mitochondrial membrane, with Manders’ coefficient analysis confirming significantly greater Parkin mitochondria overlap compared to controls. Given evidence of mitochondrial dysfunction and impaired mitophagy signaling in NHE6^-/y^ neurons, we next asked whether mitochondrial structural adaptations to stress are altered at later stages of neuronal maturation. To this end, we quantified mitochondrial volume in primary hippocampal neurons at DIV-21 under basal conditions and following mitochondrial stress induced by Antimycin A treatment (**Fig. 3h)**. Under basal conditions, mitochondrial volume was comparable between control and NHE6^-/y^ neurons **(Extended Data Fig. 10e)**, consistent with earlier observations at DIV-15 indicating preserved mitochondrial morphology in the absence of overt stress. Upon 100 nmolar Antimycin A treatment, control neurons exhibited a modest increase in mitochondrial volume, consistent with a stress-induced adaptive response. In contrast, NHE6^-/y^ neurons displayed a significantly greater increase in mitochondrial volume following stress, indicating an exaggerated or dysregulated mitochondrial remodeling response. These findings suggest that while mitochondrial structure appears largely intact under resting conditions, NHE6 deficiency alters mitochondrial plasticity under stress. Together, these findings corroborate the snRNAseq signature, confirming that NHE6 deficiency drives progressive bioenergetic impairment, oxidative stress, and dysregulated mitochondrial quality control with increasing neuronal maturity.

### NHE6^-/y^ hippocampus demonstrates strong elevations of disease-associated microglia observed in common neurodegenerative disorders

Microgliosis is a hallmark for neurodegeneration disorders, and distinct states of microglia have been identified in different neurodegenerative diseases^27^. While human postmortem brain analysis from CS patients and the NHE6^-/y^ rodent models show a prominent glial response, the role of microglia in CS disease pathogenesis remains unknown. Therefore, we subset microglial nuclei (15,357 total nuclei) from the snRNAseq dataset for detailed analysis (**Extended Data Fig. 11a**). The normalized nuclei count of microglia revealed increased microglial nuclei in NHE6^-/y^ rat at 8.5 (elevated by 228%) and 17 months (elevated by 204%) (**Fig. 4a**). To validate increase in microglia, we stained control and NHE6^-/y^ rats with IBA1 and counted the number of IBA1-positive cells. Consistent with the snRNAseq analysis, our results show a significant increase in microglial numbers at 8.5 and 17 months, while immunostaining also revealed a modest increase early at 1 month (**Extended Data Fig. 11b**). We further investigated gene expression changes in microglia between control and NHE6^-/y^ rats across the timepoints using MAST analysis. Our results identified 221 upregulated and 153 downregulated genes at 1 month, 256 upregulated and 153 downregulated genes at 8.5 months, and 458 upregulated and 383 downregulated genes at 17 months (adjusted p-value < 0.05 and log2fc > 0.1) (**Extended Data Fig. 11c; Extended Data Table.6**). GO pathway analysis showed upregulation of pathways associated with mitochondrial dysfunction, microglial phagocytic activity and activation of immune pathways at 8.5 and 17 months (**Extended Data Fig. 11d**).

**Fig 4.**
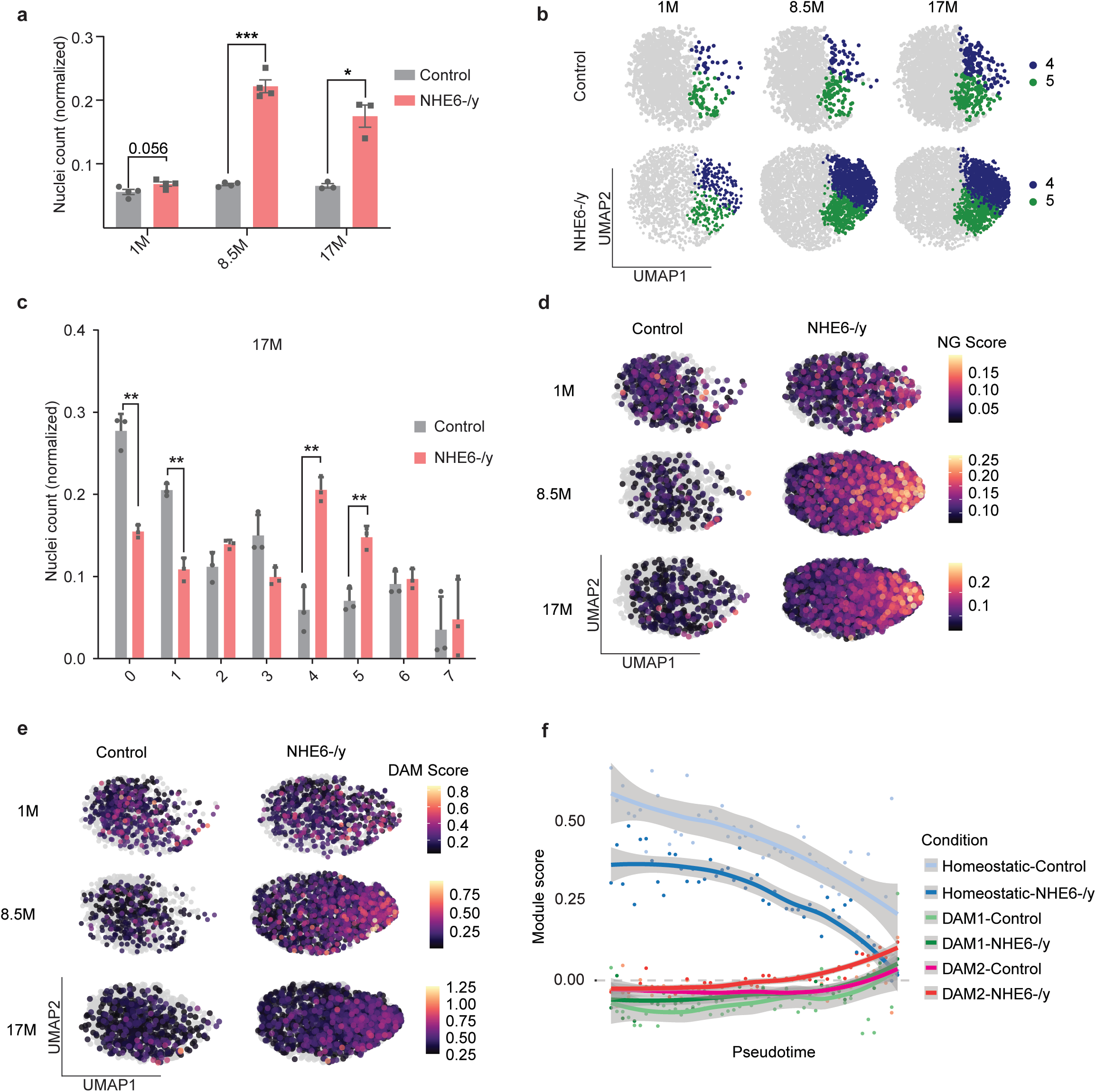
NHE6-/y hippocampus demonstrates strong elevations of disease-associated microglia observed in common neurodegenerative disorders. **a,** Barplot shows normalized count of microglial nuclei across timepoints and indicate NHE6-/y rats have increased number of microglial nuclei at 8.5 and 17 months as compared with controls. Data are presented as mean ± SEM. Each point in the graphs represents an individual brain. Two-tailed unpaired t-test with Welch’s correction was performed. ***p-value, 0.0005, *p-value, 0.02. **b,** Microglial sub-clustering UMAP at 1, 8.5 and 17 months shows distinct subclustering of microglial nuclei in cluster 4 and 5 in NHE6-/y rats. **c,** Barplot indicates normalized frequency of microglial nuclei in subclusters at 17 months in NHE6-/y rats as compared with control rats. Data are presented as mean ± SEM. Each point in the graphs represents an individual brain (Control = 3, NHE6-/y = 3). Two-tailed unpaired t-test with Welch’s correction was performed. **p-value, 0.0045 (cluster 0), **p-value, 0.001 (cluster 1), **p-value, 0.003 (cluster 4), **p-value, 0.002 (cluster 5). **d,** UMAP plot from microglial nuclei shows neurodegenerative (NG) gene expression score across timepoints split by genotypes. **e,** UMAP plot shows DAM gene expression score across timepoints split by genotypes. Color scale bars denote the gene module score. **f,** Plot depicts the module expression score of microglia states (homeostatic, DAM1 and DAM2) split by genotypes.

We next performed sub-clustering analysis of microglia nuclei, identifying eight distinct sub-clusters. Among these, clusters 4 and 5 were found to be predominantly enriched in NHE6^-/y^ rats at 1, 8.5, and 17 months as compared with control rats (**Fig. 4b, Extended Data Fig. 11e**). We termed these clusters as CS-enriched clusters. Statistical analysis revealed significant increases in microglial nuclei count in clusters 4 and 5 in NHE6^-/y^ rats across all time points, while nuclei count was decreased in clusters 0 and 1. However, the number of microglial nuclei remained consistent in clusters 2, 3, 6 and 7 in both control and NHE6^-/y^ rats (**Fig. 4c; Extended Data Fig. 11f,g**). Interestingly, CS-enriched microglial clusters correspond to previously identified genes associated with neurodegeneration and disease-associated microglia (DAM) states^27–29^. The overall neurodegenerative module score calculated by a list of neurodegenerations associated genes^30^ (NG) was progressively elevated in NHE6^-/y^ rats at 8.5 and 17 months, particularly in the CS-enriched microglial clusters (**Fig. 4d**). The DAM module score, calculated through DAM signature genes^30^, was upregulated in CS-enriched clusters at 8.5 and 17 months in NHE6^-/y^ rats, as compared to control rats (**Fig. 4e**). To confirm progressive loss of microglial homoeostatic transcriptional genes and the increase in DAM signature genes, we performed pseudo-time analysis from microglial nuclei (**Extended Data Fig. 12a**). Our results indicate that microglial nuclei from NHE6^-/y^ rats display reduced expression of homeostatic genes, with this expression declining further with age. Conversely, the expression of DAM2 genes shows an elevated expression in NHE6^-/y^ rats as compared with control (**Fig. 4f**). The expression of homeostatic genes (P2ry12, Cx3cr1, Csf1r) decreased along the pseudotime trajectory. While the expression of DAM1 genes (Apoe, Tyrobp, Ctsb) and DAM2 genes (Trem2, Axl, Cd9) increased along the pseudotime trajectory, indicating a progressive elevation of DAM signatures in NHE6^-/y^ rats with aging (**Extended Data Fig. 12b-d**). The transcriptional expression (**Extended Data Fig. 12e**) and GO analysis in CS enriched clusters as compared with other clusters indicated pathways associated with oxidative stress, phagocytosis and activation of immune pathways were upregulated at 8.5 and 17 months (**Extended Data Fig. 12f**). Collectively, these results suggest that NHE6^-/y^ rats exhibit distinct CS-enriched microglial states characterized by transcriptional profiles associated with traditional DAM and neurodegenerative diseases.

Microglia are the principal immune cells in the brain and function as phagocytic cells to clear debris of neurons after programmed cell death. Excessive microglial phagocytosis, however, can lead to neuroinflammation and neurodegeneration^31,32^. To study the impact of CS on microglial phagocytosis, we compared the expression of phagocytic marker genes in the control and NHE6^-/y^ rats and it revealed significant elevation of the phagocytic marker gene Myosin1E (*Myo1e*) (**Fig. 5a**) as well as glycoprotein non-metastatic melanoma protein B (*Gpnmb*) in the CS-enriched clusters at 8.5 and 17 months (**Fig. 5b**). To validate these findings, we performed immunohistochemical staining to assess the phagocytic characteristics in control and NHE6^-/y^ rats, focusing on the morphological characteristics of microglia using IBA1 staining and the phagocytic activity marker cluster of differentiation 68 (CD68). As anticipated from the snRNAseq analysis, microglia at 1 month showed similar morphological properties and CD68 expression in both control and NHE6^-/y^ rats (**Fig. 5c-e**). However, by 8.5 months, microglia in NHE6^-/y^ rats exhibited a phagocytic morphology, characterized by increased soma size, reduced or shorter branches, and elevated CD68 expression as compared to control rats (**Fig. 5f-h**). Consistent with 8.5 months analysis, microglia at 17 months also exhibited increased phagocytic characteristic (**Fig. 5i-k**). Taken together these results indicate that NHE6^-/y^ rats exhibit elevated phagocytic microglial activity at 8.5 and 17 months of age.

**Fig 5.**
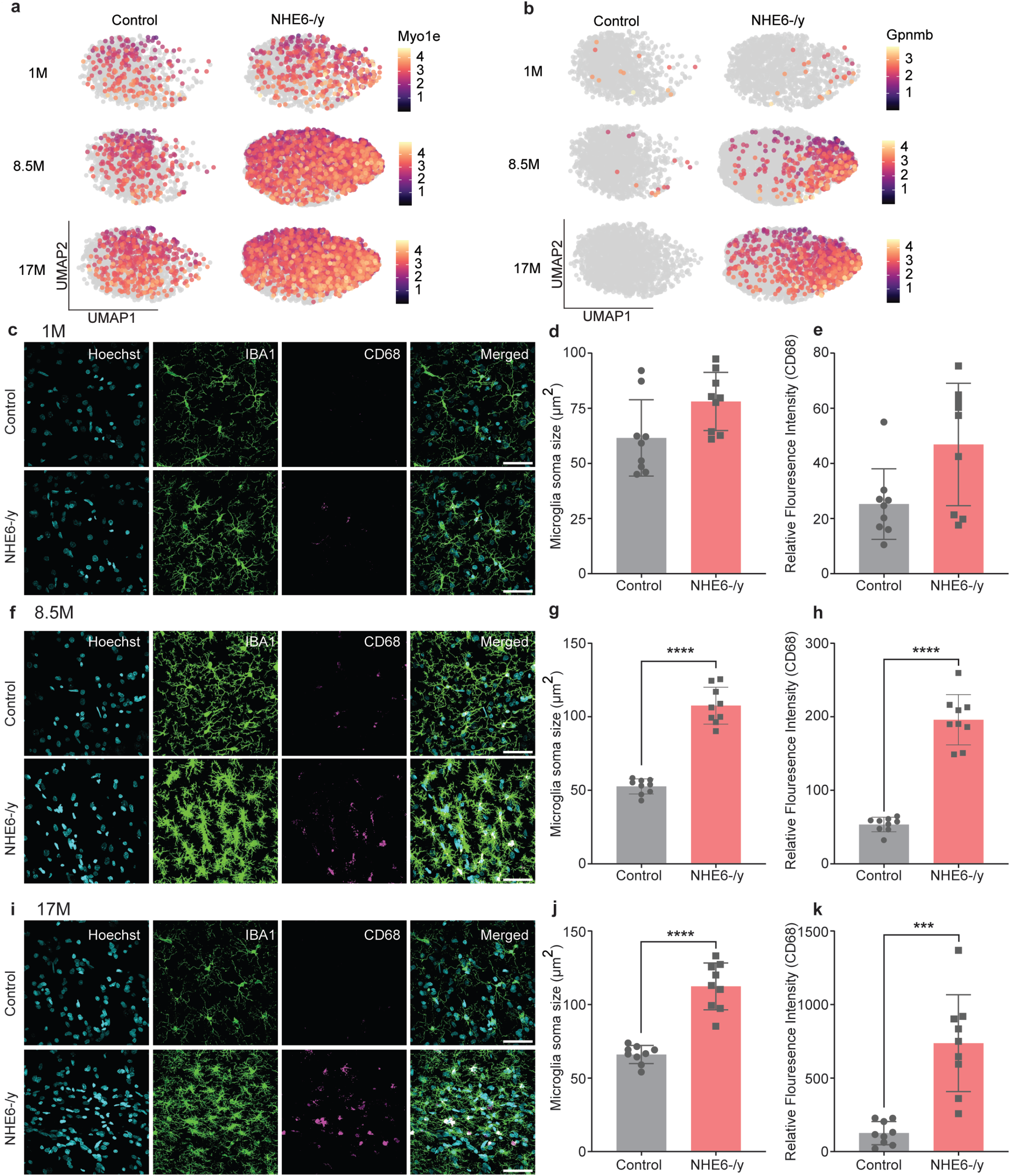
Microglia exhibit elevated phagocytic microglial activity. **a,** Feature plot shows elevated expression of phagocytic gene Myoe1 in NHE6-/y rats as compared with controls. **b,** Feature plot indicates elevated expression of phagocytic gene Gpnmb split by genotype. Color scale bar denotes gene expression. **c-e,** IBA1(green) and CD68 (magenta) staining indicate no change in morphology (d) and phagocytic microglial activity (CD68 expression) (e) at 1 month. Each point in the graphs represents an individual brain section (three sections per rat; Control = 3, NHE6-/y = 3). Data are presented as mean ± SEM. Scale bar 50 µm. **f-h,** IBA1(green) and Cd68 (magenta) staining show amoeboid morphology (increased soma size) and elevated phagocytic microglial activity (CD68 expression) in NHE6-/y rats at 8.5 months. Each point in the graphs represents an individual brain section (three sections per rat; Control = 3, NHE6-/y = 3). Data are presented as mean ± SEM. Two-tailed unpaired t-test with Welch’s correction was performed. ****p-value < 0.0001. Scale bar 50 µm. **i-k,** IBA1(green) and Cd68 (magenta) staining indicate amoeboid morphology (increased soma size) and elevated phagocytic microglial activity (CD68 expression) in NHE6-/y rats at 17 months of age respectively. Each point in the graphs represents an individual brain section (three sections per rat; Control = 3, NHE6-/y = 3). Data are presented as mean ± SEM. Two-tailed unpaired t-test with Welch’s correction was performed. ****p-value < 0.0001 and ***p-value, 0.0004. Scale bar 50 µm.

### Astrocytes exhibit neuroinflammatory signatures in NHE6^-/y^ rats

Similar to microglia, the astrocyte response is also pathology-dependent, and specific disease associated states of astrocytes (DAA) have been linked with progression of neurodegenerative diseases^33^. Increased astrocyte response has been reported in the mouse and rat brain of CS rodent models, but there is very limited knowledge of the role of astrocytes in CS^34^. To study the role of astrocytes in CS, we investigated astrocyte nuclei (20,216 total nuclei) from our snRNAseq dataset **(Extended Data Fig. 13a)**. We observed reduced numbers of astrocyte nuclei in NHE6^-/y^ rats at 8.5 months but no significant change was observed at 1 and 17 months (**Fig. 6a**). However, quantification of ALDH1L1 and GFAP positive cells revealed no change in astrocyte numbers at,1 but a significant increase at 8.5 and17 months (**Extended Data Fig. 13b,c**), consistent with prior immunohistochemical studies of mouse hippocampus. Differences between the snRNAseq analysis and immunohistochemistry likely reflect technical considerations specific to astrocytes, discussed below in the Discussion, and noted previously in the literature^35–38^. From differential gene expression in astrocytes, we identified: 384 upregulated and 328 downregulated genes at 1 month; 194 upregulated and 132 downregulated genes at 8.5 months; and 885 upregulated and 577 downregulated genes at 17 months (with an adjusted p-value < 0.05 and log2FC > 0.1) **(Extended Data Table.7)**. Astrocyte sub-clustering analysis identified eight distinct astrocyte clusters, with cluster 5 being predominantly upregulated in the astrocytes of NHE6^-/y^ rats at 8.5 and 17 months (**Fig. 6b, c; Extended Data Fig. 13d,e)**. A comparison of the module scores for Disease DAA genes revealed that cluster 5 exhibited elevated expression of DAA genes at 8.5 and 17 months (**Fig. 6d**). GSEA pathway analysis from overall astrocytes using hallmark and KEGG datasets indicated an upregulation of inflammatory response (**Fig. 6e**). Additionally, CellChat signaling analysis also revealed upregulation of the Prosaposin (*PSAP*) pathways in cluster 5, indicating enhanced innate immune and inflammatory responses^39,40^ (**Fig. 6f**). These results indicate that NHE6^-/y^ rat astrocytes exhibit inflammatory responses and upregulation of notable AD associated cellular responses.

**Fig 6.**
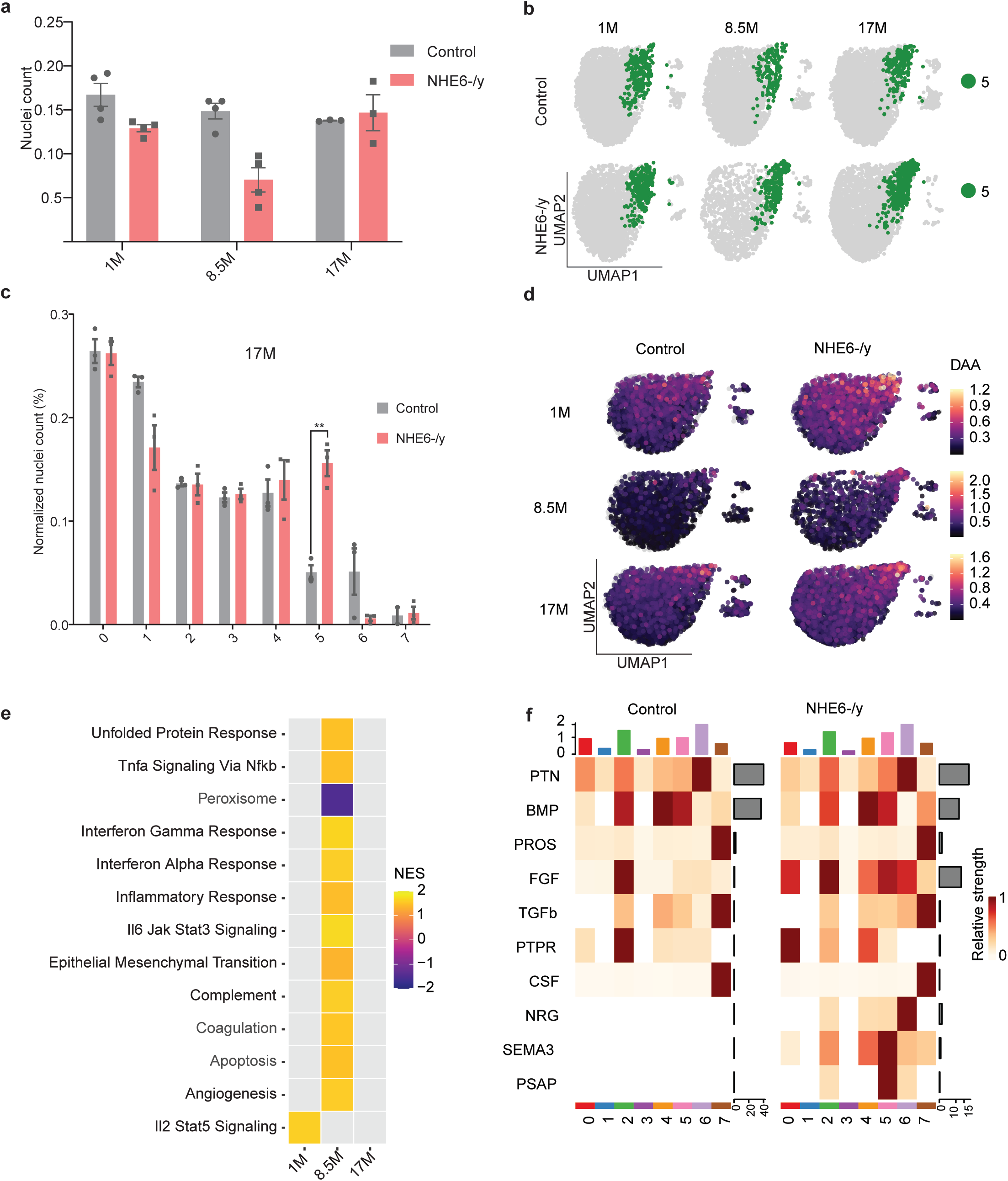
Astrocytes exhibit neuroinflammatory signatures in NHE6-/y rats strongly overlapping with common neurodegenerative disorders. **a,** Barplot shows overall number of astrocytes nuclei normalized with total number of nuclei from each animal at 1, 8.5 and 17 months. Data are presented as mean ± SEM. Two-tailed unpaired t-test with Welch’s correction was performed. p-value 0.0038 (8.5M). **b,** UMAP plots show astrocytes nuclei sub-clustering at 1, 8.5 and 17 months. **c,** Barchart indicates normalized frequency of astrocytes nuclei within each cluster at 17 months. Data are presented as mean ± SEM. Each point in the graphs represents an individual brain (Control = 3, NHE6-/y = 3). Two-tailed unpaired t-test with Welch’s correction was performed. **p-value, 0.004. **d,** UMAP plot shows DAA gene expression score across timepoints split by genotypes. Color scale bars denote the DAA gene module score. **e,** Heatmap show enriched pathways in astrocytes nuclei from hallmark and KEGG datasets across timepoints. Color show NES. **f,** Heatmap show Cell chat communication analysis at 8.5 months within each cluster.

### Oligodendrocytes show reduced myelination capacity in NHE6-/y rat hippocampus

Oligodendrocyte dysfunction has been linked to the pathogenesis of neurodegenerative diseases such as Alzheimer’s disease (AD), leading to the identification of disease-associated oligodendrocyte states (DAO)^37,41^. In our study, we also focused on oligodendrocytes (35,460 total nuclei) to explore their role in the pathogenesis of CS where oligodendrocytes have been understudied (**Extended Data Fig. 14a**). Normalized nuclei count of oligodendrocytes indicated reduced oligodendrocytes nuclei in NHE6^-/y^ rats at 8.5 and 17 months as compared with controls (**Fig. 7a**). To validate loss of oligodendrocytes, we stained control and NHE6^-/y^ rats at 8.5 and 17 months using an Olig2 antibody. We observed a significant reduction in oligodendrocyte numbers at 17 months in NHE6^-/y^ rats, while no significant change was observed at 8.5 months (**Extended Data Fig. 14b**). Further sub-clustering of oligodendrocytes identified five distinct clusters, which were classified into two major subtypes; newly formed and myelin-forming oligodendrocytes. Newly formed oligodendrocytes were defined by the expression of Neu4, Sox6, and Bmp4, whereas myelin-forming oligodendrocytes were annotated by the expression of Mal, Mog, and Opalin^42^ (**Fig. 7b**, **Extended Data Fig. 14c,d**). It is interesting to note that the number of newly formed oligodendrocytes were significantly increased in NHE6^-/y^ rats at 8.5 and 17 months; however, the number of myelin forming oligodendrocytes was reduced in NHE6-^/y^ rats at 8.5 and 17 months (**Fig. 7c**). We examined DEGs in myelin forming oligodendrocytes and identified: 709 upregulated and 493 downregulated genes at 1 month; 769 upregulated and 277 downregulated genes at 8.5 months; and 1025 upregulated and 434 downregulated genes at 17 months (with an adjusted p-value < 0.05 and log2FC > 0.1) (**Fig. 7d; Extended Data Table.8**). Pathway analysis using hallmark and KEGG datasets indicated an upregulation of innate immune pathways related to inflammatory responses and TNF-alpha signaling via NF-kB, suggesting activation of the innate immune pathways (**Fig. 7e**). Together these results indicate reduced myelinating capacity of oligodendrocytes in NHE6^-/y^ hippocampus as compared with controls, along with an increased inflammatory response and activation of innate immune pathway.

**Fig 7.**
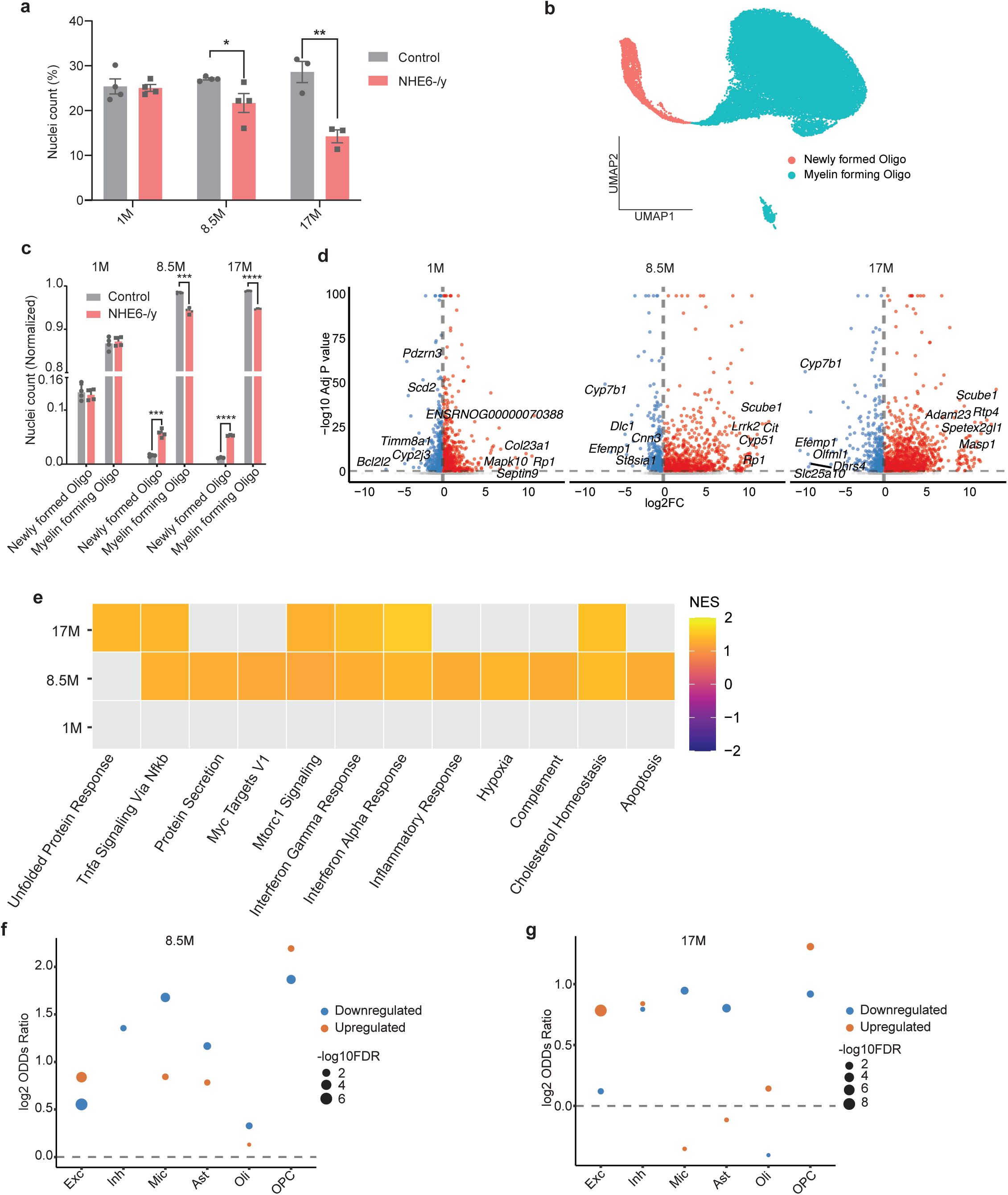
Oligodendrocytes show reduced myelination capacity in NHE6-/y rat hippocampus. **a,** Barplot depicts number of oligodendrocytes nuclei normalized with total number of nuclei from each animal. Data are presented as mean ± SEM. Each point in the graphs represents an individual brain. Two-tailed unpaired t-test with Welch’s correction was performed. *p-value, 0.038, ** p-value, 0.007. **b,** UMAP plot highlights two subtypes of oligodendrocytes; Newly formed Oligo and Myelin forming Oligo. **c,** Barplot shows reduced number of myelin forming oligodendrocytes at 8.5 and 17 months. Nuclei were normalized with the total number of oligodendrocytes from each animal. Data are presented as mean ± SEM. Each point in the graphs represents an individual brain. Two-tailed unpaired t-test with Welch’s correction was performed. ***p-value, 0.009 (Newly formed Oligo, 8.5M), ***p-value, 0.008 (Myelin forming Oligo, 8.5M), ****p-value < 0.0001. **d,** Volcano plots show DEGs from myelin forming oligodendrocytes across timepoints. Significant genes are labelled in color (Padj < 0.05, Log2FC > 0.1), while nonsignificant genes are shown in gray. Top 5 upregulated and downregulated genes are labelled. **e,** Heatmap show GSEA enriched pathways among hallmark and KEGG terms from myelin forming oligodendrocytes at all timepoints. **f,** Dot plot shows significant overlap of concordant DEGs shared between the 8.5 months old NHE6-/y rat and AD dataset across six major cell types. Dot color indicates direction of regulation (up- or downregulated), dot size indicates statistical significance, expressed as log10(FDR-adjusted p-value). Enrichment of concordant DEG overlap was examined for each cell type and direction of regulation (up- or downregulation) using Fisher’s exact test, with odds ratios and significance calculated relative to a background gene universe. P values were adjusted for multiple comparisons across all cell types and directions using the Benjamini-Hochberg method (FDR < 0.05). **g,** Dot plot shows significant overlap of concordant DEGs shared between the 17 months old NHE6-/y rat and AD dataset across six major cell types. Dot color indicates direction of regulation (up- or downregulated), dot size indicates statistical significance, expressed as log10(FDR-adjusted p-value). Enrichment of concordant DEG overlap was examined for each cell type and direction of regulation (up- or downregulation) using Fisher’s exact test, with odds ratios and significance calculated relative to a background gene universe. P values were adjusted for multiple comparisons across all cell types and directions using the Benjamini-Hochberg method (FDR < 0.05).

Additionally, we performed transcriptomic analysis of OPC (8,390 nuclei) across all timepoints. Sub-clustering analysis of OPC cells identified five distinct clusters (**Extended Data Fig. 15a**), with no significant change in the number of nuclei within each cluster across all timepoints (**Extended Data Fig. 15b-d**). Differential gene expression analysis in OPC identified; 131 upregulated and 122 downregulated genes at 1 month; 44 upregulated and 111 downregulated genes at 8.5 months; and 259 upregulated and 272 downregulated genes at 17 months (with an adjusted p-value < 0.05 and log2FC > 0.1) (**Extended Data Fig. 15e; Extended Data Table.9**). GSEA revealed early metabolic and inflammatory response in OPC, followed by robust activation of growth factor driven PI3K/Akt signaling (**Extended Data Fig. 15f)**.

### Strong overlap between dysregulated genes in aged CS rat and in human postmortem Alzheimer’s Disease hippocampus

Our studies revealed dysregulation of many important pathways in the CS rat hippocampus that are also altered in Alzheimer’s disease based on the literature. To this end, we therefore conducted a direct and systematic study of the degree of overlap in our snRNAseq dataset and robust, publicly-available snRNAseq studies from human postmortem hippocampus from patients with AD. We compared our data with snRNAseq data from postmortem hippocampus from 48 ROSMAP individuals selected based on modified NIA Reagan criteria and Braak staging, including 26 individuals with a positive pathological diagnosis of AD and 22 controls^43^. For analysis, we restricted the comparison to hippocampal snRNAseq data from ROSAMAP and used our 8.5 and 17 months timepoints from the CS rat snRNAseq data. At 8.5 months, excitatory neurons showed the most significant concordance in both downregulated and upregulated genes, indicating conserved neurodegenerative mechanisms. Microglia, astrocytes, oligodendrocytes and OPC also demonstrated significant overlap among downregulated genes, suggesting shared neuroinflammatory and glial stress responses (**Fig. 7f, Extended Data Table.10**). At 17 months, overlap with AD associated transcriptional changes was also shared across cell types. Excitatory neurons again showed significant concordance among upregulated genes, while OPCs showed concordance in both upregulated and downregulated genes. Microglia and astrocytes primarily showed overlap among downregulated genes (**Fig. 7g, Extended Data Table.11**). Together, these results demonstrate that the NHE6^-/y^ rat transcriptome shows broad concordance with the transcriptome changes of AD pathology in humans.

### Early activation of cGAS-STING pathway in NHE6^-/y^ rats

To complement the transcriptome studies, we performed unbiased proteomic liquid chromatography-tandem mass spectrometry (TMT-MS) analysis of hippocampus tissue from a total of 30 samples; control (n=7) and NHE6^-/y^ (n=8) rats at 1.5 and 8.5 months. We found that replicates within control and NHE6^-/y^ rat clusters separately by principal component analysis and replicate sample coefficients of variation are low, demonstrating proteomic differences between control and NHE6^-/y^ rats and excellent replicate reproducibility (**Extended Data Fig. 16a-d**). Interestingly, global protein abundance analysis revealed significant upregulation of DAM and DAA associated proteins in the NHE6^-/y^ rats at both timepoints (**Extended Data Table.12)**. Additionally, we also observed a reduction in the expression of myelination proteins myelin basic protein (*Mbp*) and myelin oligodendrocyte glycoprotein (*Mog*) at 8.5 months, again indicating decreased myelination capacity in NHE6^-/y^ rats (**Fig. 8a, Extended Data Table.12)**. We also validated reduced expression of *Mbp* in NHE6^-/y^ rats through western blot at 8.5 months (**Extended Data Fig. 16e**). MS analysis also showed increased expression of phagocytic genes, *Cd68* and *Gpnmb* at 8.5 months (**Fig.8a**). GO analysis indicated increased activity of innate immune response and microglial activation of phagocytic and AD associated amyloid beta or tau pathways (**Fig. 8b**). We also examined mitochondrial dynamics related proteins and identified 12 differentially expressed proteins associated with mitochondrial dysfunction at 8.5 months (**Extended Data Fig. 16f**). Moreover, to identify disease associations from MS analysis at 8.5 months, we performed Disease Gene Network (DisGeNET) enrichment analysis and found that 20 genes were associated with AD (**Extended Data Fig. 16g,h**). These findings were consistent with our snRNAseq analysis.

**Fig 8.**
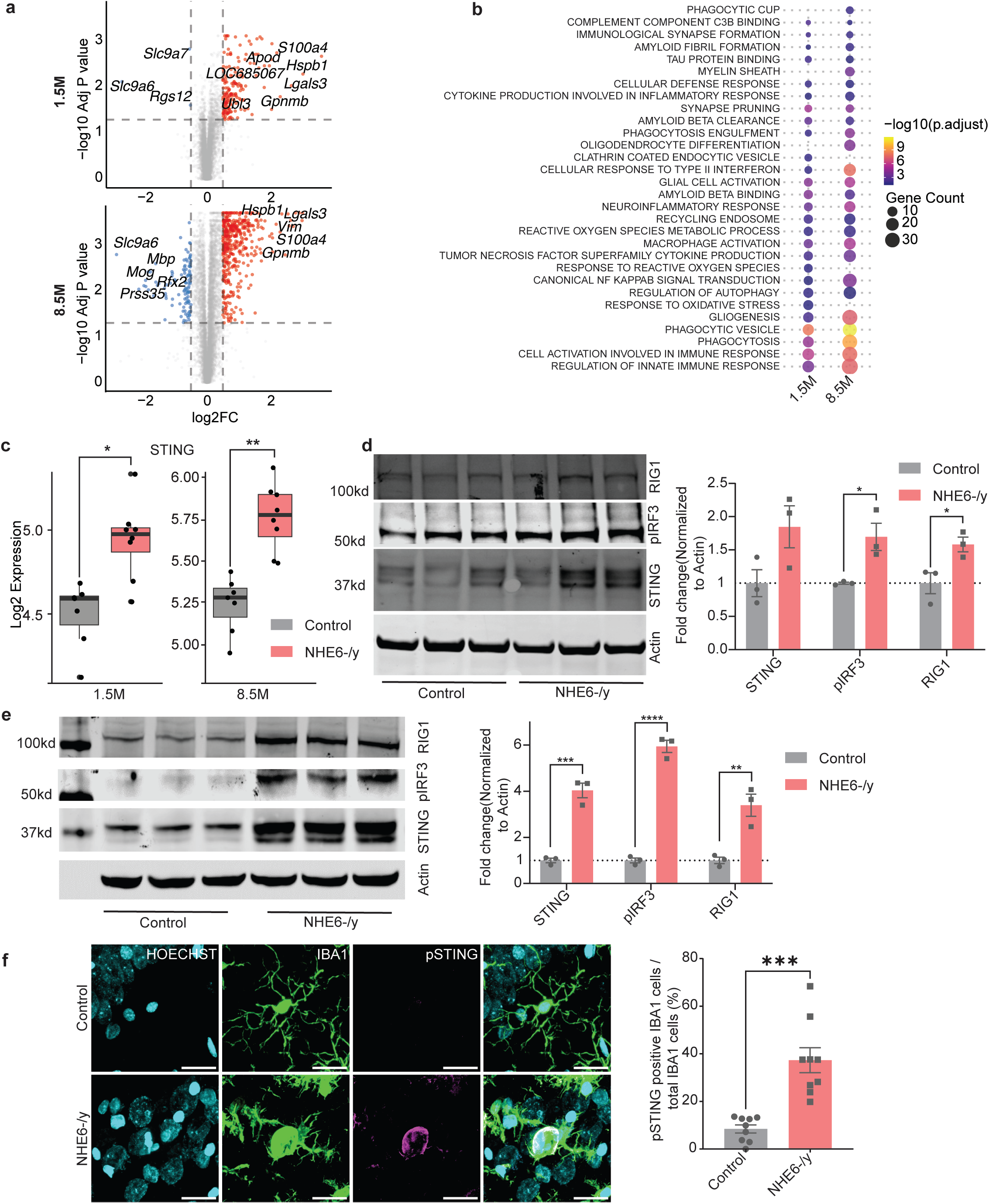
cGAS-STING pathway upregulates in NHE6-/y rats. **a,** Volcano plots show differentially abundant proteins identified by MS analysis from the hippocampus of NHE6-/y rats at 1.5 and 8.5 months. Significant proteins are labelled in color (log2FC > 0.5, q-value < 0.05), while nonsignificant proteins are shown in gray. Top 5 upregulated and downregulated proteins are labelled. Differential abundance was analyzed using Welch’s t-test with Benjamini-Hochberg FDR correction on log-transformed data. **b,** GO analysis shows significant enriched terms at 1.5 and 8.5 months from MS analysis. **c,** Box plots show abundance of STING at 1.5 and 8.5 months. Each point in the graphs represents an individual brain (Control = 7, NHE6-/y = 8). Data are presented as mean ± SEM. Differential abundance was analyzed using Welch’s t-test with Benjamini-Hochberg FDR correction on log-transformed data. *q-value, 0.02, **q-value, 0.005. **d,** Representative western blot and quantifications show elevation of STING, pIRF3 and RIG1 expression in the hippocampus of NHE6-/y rats at 1 month. Each point in the graphs represents an individual brain (Control = 3, NHE6-/y = 3). Data are presented as mean ± SEM. Protein levels were normalized to actin. Two-tailed unpaired student’s t-test with was performed. *Pvalue 0.02 (pIRF3), *pvalue, 0.03 (RIG1). **e,** Representative western blot and quantification show elevation of STING, pIRF3 and RIG1 expression in the hippocampus of NHE6-/y rats at 8.5 months. Each point in the graphs represents an individual brain (Control = 3, NHE6-/y = 3). Data are presented as mean ± SEM. Protein levels were normalized to actin. Two-tailed unpaired student’s t-test was performed. ***p-value, 0.0008 (STING), ****p-value<0.00001 (pIRF3), **p-value, 0.008 (RIG1). **f,** Immunofluorescence co-staining of IBA1 (green) and pSTING (magenta) shows enriched pSTING expression in NHE6-/y rats microglia at 8.5 months. Data are presented as mean ± SEM. Each point in the graphs represents an individual section (three sections per rat; Control = 3, NHE6-/y = 3). Two-tailed unpaired t-test with Welch’s correction were performed. ***p-value, 0.0004. Scale bar 20 µm.

Activation of the innate immune pathway via the cGAS-STING pathway contributes to neuronal death in conditions associated with lysosomal and mitochondrial dysfunction^44–46^. Age associated inflammation has also been shown to activate STING signaling, driving microglial activation and subsequent cognitive decline and neurodegeneration^30^. Given that CS impairs both lysosomal and mitochondrial function, and that dysfunction in these organellar systems is known to activate the cGAS-STING pathway^44^, we investigated protein changes in the cGAS-STING pathway in CS. Our MS analysis shows significant upregulation of cGAS-STING pathway associated proteins, including *STING1* (**Fig. 8c**), *STAT1*(**Extended Data Fig. 17a**), and *NFKB1*(**Extended Data Fig. 17b**), at both 1.5 and 8.5 months, indicating activation of the cGAS-STING pathway in NHE6^-/y^ hippocampus. To further validate these findings, we performed western blot analysis of hippocampus tissue from control and NHE6^-/y^ rats at P0 (postnatal day 0), 1 month, and 8.5 months. At P0, our results indicated that there was no change in the expression of STING pathway proteins between control and NHE6^-/y^ rats using the expression of *STING*, phosphorylated IRF3 (*pIRF3*), and retinoic acid-inducible gene I (*RIG1*), a well-defined cytosolic dsRNA sensor to induce type-I interferon (**Extended Data Fig. 17c**). Interestingly, the expression of STING pathway proteins was elevated at 1 month in NHE6^-/y^ rats indicating the activation of STING pathway before the activation of microglial reactive states (**Fig. 8d**). Moreover, the expression of cGAS-STING pathway proteins including *STING*, *pIRF3* and *RIG1* were also elevated in NHE6^-/y^ rats at 8.5 as compared with controls (**Fig. 8e**). We further confirmed significant elevation of phospho-STING (pSTING) in NHE6^-/y^ rats at both 1 and 8.5 months by immunostaining of hippocampus tissue (**Extended Data Fig. 17d**). To identify the role of specific cell types in STING activation, we immunostained hippocampal tissue using anti-pSTING combined with microglial (IBA1), neuronal (NeuN) and astrocytes (ALDH1L1) markers and identified that pSTING was predominately expressed and localized to microglia **(Fig. 8f)** in NHE6^-/y^ rats, as compared with neurons and astrocytes **(Extended Data Fig. 17e)**. Taken together, these results demonstrate that the cGAS-STING pathway is activated at an early stage in NHE6^-/y^ rats and may play a critical role in CS mediated neurodegeneration and neuroinflammation.

## Discussion

Endolysosomal defects in CS cause both neurodevelopmental and neurodegenerative abnormalities. In the current study, we define the transcriptional dynamics of CS pathogenesis at single-cell resolution across the lifespan in a robust rat model, and we thereby uncover underlying mechanisms associated with neurodegeneration. Importantly, we identify abnormalities in mitochondrial function early in both excitatory and inhibitory neuronal cell types. We subsequent demonstrate mitochondrial dysfunction in functional experiments, both in vitro and ex vivo, that support a mechanism of defective mitochondrial quality control and impaired mitochondrial bioenergetics in neurons. This linkage between a primary endolysosomal dysfunction in neurons and likely secondary defectives in mitochondrial function has broad significance in neurodegeneration. We further provide evidence for activation of innate immunity, including the cGAS-STING pathway which is known to be activated by lysosomal and mitochondrial dysfunction^45,47^. Further, our study demonstrates that multiple dynamic transcriptional profiles in this CS rat have striking parallels to human postmortem AD brains. We thereby suggest that our CS rat serves as a unique, single gene mutant that may inform neurodegenerative mechanisms in more complex disorders such as AD.

Mitochondrial dysregulation, specifically the loss of efficiency in the electron transport chain and ATP synthesis, has been previously implicated in various neurodevelopmental and neurodegenerative disorders^48–50^. Mitochondrial alterations, including swollen and ruptured mitochondria with small vacuolations, has only been reported in one previous study in CS, in patient-derived Epstein–Barr virus lymphoblastoid cell lines (EBV-LCLs)^51^. In snRNAseq analysis from excitatory and inhibitory neurons, we found transcriptional signatures reflecting impaired electron transfer in complex I at 1 and 8.5 months. We also found transcriptional signatures for dysregulated reactive oxygen species and oxidative phosphorylation pathways in both excitatory and inhibitory neurons (**Fig. 2a**). NHE6 deficiency disrupts mitochondrial homeostasis through a progressive, maturation-dependent sequence of dysfunction in hippocampal neurons. We identified reduced Δψm at DIV-15, in the absence of any detectable changes in mitochondrial mass, morphology and respiration (**Fig. 3a**). As neurons matured to DIV-21, mitochondrial dysfunction expanded beyond Δψm impairment and NHE6^-/y^ neurons exhibited elevated mitochondrial superoxide and reduced maximal and spare respiratory capacity, demonstrating declining respiratory reserve and diminished bioenergetic resilience (**Fig. 3b,c**). Because neurons depend on spare respiratory capacity to sustain synaptic transmission and calcium buffering during periods of increased activity^52^, this loss likely increases vulnerability to activity-dependent stress and may contribute to neuronal dysfunction in CS. Our complementary ex vivo mitochondria findings from 1 month hippocampus, including electron flow and coupling assays, show respiratory impairment spanning Complex I-IV together with preserved proton leak and substrate-independent reductions in respiratory control ratio, ruling out an isolated ETC entry-point defect and instead suggesting broader disruption of oxidative phosphorylation efficiency, potentially involving altered ETC organization or supercomplex stability (**Fig. 3d-f**). We also observed early activation of mitochondrial quality-control pathways. Increased Parkin recruitment at DIV-21, despite preserved morphology and bulk respiration, indicates that PINK1-Parkin signaling responds to early membrane depolarization before overt respiratory failure develops (**Fig. 3g**). Collectively, the current findings, combined with our prior studies demonstrating lysosomal dysfunction resulting from loss of NHE6, potentially supports a progressive model in which NHE6 loss causes impaired mitophagy, leading to build up of damaged, failing mitochondria. When treated with Antimycin A which induces mitophagy, NHE6^-/y^ neurons demonstrate an increase in mitochondrial volume relative to control which suggests impairments in mitophagy (**Fig. 3h**). Overall, the mitochondrial functions studies validate the mitochondrial stress signatures identified by snRNAseq and positions mitochondrial dysfunction as a central downstream consequence of NHE6 deficiency and a potential driver of neuronal vulnerability in CS.

Intriguingly, we show that NHE6 is predominantly expressed in neuronal cells, with notably higher expression in inhibitory and excitatory neurons. Conversely, NHE6 expression is very low in astrocytes and microglia (**Extended Data Fig. 4d, 5a**). These results suggest that loss of NHE6 may exert non-cell-autonomous effects in astrocytes and microglia. Few studies have explored microglial states in rats at the single-cell transcriptome level, and they generally lack comprehensive analysis across the lifespan and in neurodegenerative models^53–55^. Although, the activation of microglia and astrocytes in CS has been established previously^13,15^, no study has focused on the molecular identification of distinct states of these glial cells. Using snRNAseq we define the “CS-enriched” microglia transcriptomes across the lifespan. This CS-enriched microglia transcriptome also has overlap with microglial signatures in the aged normal control; however, the CS transcriptome emerges much earlier in the lifespan in CS rats (by 8.5 months) (**Fig. 4b**). Notably, the CS-enriched microglia states also express elevated transcriptional expression of DAM and NG genes along with upregulation of phagocytosis, inflammatory, innate immune, cellular response to amyloid beta and synaptic pruning pathways (**Fig. 4d,e**). The phagocytic expression of microglia was also confirmed by augmented CD68 expression at 8.5 and 17 months, accompanied by enlarged soma size and reduced branching morphology (**Fig. 5c-h**). Our MS analysis also supported microglial phagocytic activity and innate immune activation across different timepoints.

It is very important to note that we did not observe an increase in the number of astrocyte nuclei in NHE6^-/y^ rats at 1, 8.5 and 17 months of age. Instead, a significant decrease in astrocyte nuclei was observed at 8.5 months (**Fig. 6a**). This was unexpected given our previous reports of astrogliosis in NHE6^-/y^ rats at 3 months^13^. The decrease in nuclei number is not observed in our immunohistochemistry studies of ALDH1L1 and GFAP positive cell counts **(Extended data figure 13b,c)**. We believe this discrepancy reflects technical limitations of snRNAseq methods regarding astrocyte studies rather than a true reduction in astrocyte number. Astrocytic nuclei are small and well-known to be harder to recover than larger neuronal nuclei during drople-based isolation^35,36^, and are consistently underrepresented relative to their true brain abundance across snRNAseq studies^37,38,56^. This limitation may be augmented during active astrogliosis, as reactive astrocytes show reduced intron retention^57^, further diminishing the nuclear signal needed to detect reactive astrocytes and reactive astrocytes are particularly elevated at the 8.5 months. As the 8.5 months timepoint shows this potential technical artifact, we acknowledge that this may affect downstream data analyses; however, results from these astrocyte transcriptome analyses are consistent across timepoints (where the technical issue is not observed) and results were generally otherwise validated with other western, proteomic and microscopy studies. The snRNAseq analysis revealed a “CS-enriched” astrocyte cluster (cluster 5) in NHE6^-/y^ rats at both 8.5 and 17 months, showing elevated expression of DAA genes within this cluster (**Fig. 6d**). Additionally, MS analysis indicated upregulation of the vimentin (*Vim*) gene, a marker of astrocyte reactivity (**Fig. 8a**). We also observe in astrocytes transcriptional upregulation of *PSAP,* a conserved lysosomal protein and a neurotrophic factor, which is associated with lysosomal storage and neurodegenerative disorders. Its upregulation has been observed in the hippocampus of Alzheimer’s disease (AD) patients^58–60^. Similarly, prior work has suggested that the overall number of myelinating oligodendrocytes was decreased with age in AD patients and mouse AD models^61–63^. In line with these results, our data show that the number of myelinating oligodendrocytes was reduced in the NHE6^-/y^ rats at 8.5 and 17 months (**Fig. 7c**). MS analysis at 8.5 months also shows reduced expression of myelination proteins (Mbp, Mog) (**Fig. 8a**). Thus, reduced number of myelinating oligodendrocyte and myelination genes suggests that oligodendrocytes defects in NHE6^-/y^ rats may play a role in CS progression and share common patterns with other neurodegenerative diseases.

Our concordant overlap analysis with the transcriptome of human AD postmortem hippocampus shows significant overlap in downregulated genes across most cell types including excitatory neurons, microglia, astrocytes and OPC at both the 8.5 and 17 months timepoints, whereas significant overlap in upregulated genes was observed in excitatory neurons at both timepoints. Excitatory neurons showed the most robust and consistent overlap signal overall, with significant concordance in both upregulated and downregulated genes at 8.5 months. Downregulated genes in microglia and astrocytes also showed significant overlap at both ages, consistent with a shared neuroinflammatory and glial response across disease progression. Notably, although the number of overlapping genes from OPC was relatively small, it consistently exhibited the largest effect sizes (odds ratios) among significant results at both 8.5 and 17 months, suggesting a highly conserved AD associated gene expression (**Fig. 7f-g**). Thus, the transcriptome of NHE6^-/y^ rat broadly overlaps with the transcriptome of AD pathology in human.

One key finding of this study is impaired mitochondrial function in neurons, which has been associated with lysosomal storage disorders that activate the cGAS-STING pathway, making it a promising therapeutic target for the treatment of lysosomal dysfunction diseases^44^. Our proteomic and western blot analysis revealed significant upregulation of cGAS-STING associated proteins as early as 1 month and persisting through 8.5 months. Notably, this activation preceded the onset of glial reactivity, suggesting that cGAS-STING activation is an early process in NHE6^-/y^ brain. Our cell type specific immunostaining demonstrated a strong and specific upregulation of pSTING in microglia, with no significant changes observed in neurons or astrocytes (**Fig. 8f**). This microglial specificity suggests that STING is activated cell non-autonomously within microglia in NHE6^-/y^ rats, likely as a consequence of the mitochondrial and endolysosomal dysfunction in neurons. The microglia specific pattern of STING activation is consistent with a previous report showing that neuronal mitochondrial dysfunction can drive cell non-autonomous activation of the cGAS-STING pathway in microglia, contributing to neurodegeneration^64^.

Our study provides important biological insights in CS that may serve to inform more complex neurodegenerative diseases. The dataset will also serve as a valuable resource for the community of researchers studying endolysosome and neurodegenerative disorders. While extensive datasets are available from mouse models, traditional neurodegenerative pathology often fail to fully develop in mice, requiring transgenic expression of mutant proteins. Our rat dataset serves an important niche as this model demonstrates broadly significant features of neurodegeneration endogenously (resulting from one genetic mutation and not transgenic overexpression); thereby, these data offer a robust resource for studying and comparing neurodegenerative processes between human and animal models. In conclusion, using snRNAseq profiles from control and NHE6^-/y^ rats across the lifespan, our study establishes a model for CS disease progression involving impaired endolysosomal and mitochondrial function in neurons at early time points, which triggers innate immunity, including the activation of the cGAS-STING signaling pathway. The upregulation of this innate immune pathway subsequently leads to the activation of glial states, likely contributing to neurodegeneration and cognitive decline in CS. Intriguingly, disease specific cell states were similar to disease associated states observed in other neurodegenerative diseases such as AD, thereby providing further support for an endolysosomal role in these disorders.

## Acknowledgements

This research was supported by funding for HR and EMM (R01NS121618, R01AG087455, R01NS113141).

## Author Contribution

H.R. and E.M. designed the study. H.R. performed snRNAseq experiments, processed and analyzed the snRNAseq data. H.R. performed all IHC, WB experiments and analyzed the data. H.R. performed dissection, processed and analyzed the MS data. S.K.S. designed, performed, analyzed and wrote mitochondrial function experiments. J.D. analyzed the NeuN data. G.K. assisted in rat colony. Y.L. performed dissection of 1.5 months tissue for MS. H.R. and E.M. wrote and edited the manuscript. E.M. supervised the entire study.

## Declaration of Interest

The authors declare no competing interests.

## RESOURCE AVAILABILITY

### Lead contact

Further information and requests for resources and reagents should be directed to and will be fulfilled by the lead contact, Eric Morrow

### Materials availability

Animals generated in this study are available upon request.

### Data availability

Fastq files for raw snRNA-seq and Seurat objects were deposited at the Gene Expression Omnibus. The proteomics data have been deposited in PRIDE via ProteomeXchange. These datasets will be made publicly available upon completion of the publication. Human hippocampal AD datasets were obtained from AD Knowledge Portal on Synapse; syn52408592 under controlled use conditions set by human privacy regulations.

### Code availability

- This manuscript does not report new code.
- All data reported and any additional information required for reanalysis of the data in this paper will be shared by lead contact upon request.

## METHOD DETAILS

### Experimental design

NHE6^-/y^ rats were generated on a Sprague-Dawley background using CRISPR-Cas9 genomeediting technology, by inducing frameshifts and a premature stop codon in exon 7, as described in previous report^13^. For the lifespan analysis of control and NHE6^-/y^ rats for snRNAseq analysis, three timepoints were chosen: 1 month (4 animals per condition), 8.5 months (4 animals per condition), and 17 months (3 animals per condition). At these times, single nuclei were isolated from the hippocampus of control and NHE6^-/y^ male rats. To reduce batch effects, samples from each time point were processed and analyzed simultaneously. Rats were maintained and utilized in compliance with the Brown University IACUC-approved protocol (IACUC 29-09-0012) and national and institutional guidelines were followed. Our snRNAseq datasets are the first datasets generated from the hippocampus of control and NHE6^-/y^ rats at multiple aging timepoints and can serve as a valuable resource for endolysosomal and neurodegenerative diseases.

### Nuclei isolation

Working on ice throughout, nuclei were isolated from hippocampus tissue from control and NHE6^-/y^ rats by following manufacturer’s instructions from Nuclei Isolation Kit (Millipore sigma NUC201). Briefly, 5ml lysis buffer was prepared for each animal and hippocampal tissues were carefully minced with the tight pestle in the Dounce homogenizer for at least 20 times. The nuclei suspension was filtered through a 40-micron cell strainer to remove residual tissues. For sucrose purification step, nuclei suspensions were centrifuged at 30,000g for 45 min at 4 C. Nuclei pellets were dissolved in the nuclei storage buffer and the number of nuclei was counted by using a hemocytometer. Finally, 6,000 nuclei (10,000 nuclei for 17months) from each rat were used as input to 10x Genomics Chromium Controller using the Chromium Next GEM Single Cell 3′ Reagent kits v3.1(dual index). cDNA libraries from nuclei were generated and quantified by following the manufacturer’s instructions (10 X Genomics). The final barcoded libraries were generated through Dual Index Plate TT Set-A kit by following manufacturer instructions (10X genomics) and were sequenced using the Illumina Hiseq (8 months old) and Novaseq (1 and 17 months old) sequencing platforms with 150-bp paired-end reads. The average sequencing depth was 51,404 reads per cell at 1 month, 61,860 reads per cell at 8.5 months, and 57,417 reads per cell at 17 months.

### Pre-processing of snRNAseq data

Sequence alignment to the Rattus norvegicus reference genome (mRatBN7.2.109) was performed by using the CellRanger-7.0.0 software provided by 10X genomics. Since nuclear RNA is expected to include roughly equal proportions of exonic and intronic reads, default parameters were used with the option --include intron flags. The resulting raw feature barcode matrix files were read into either Python or R studio for further processing and analysis.

### Ambient mRNA removal

Ambient RNA in snRNAseq data can introduce batch effects or systematic biases during downstream analysis. The CellBender Python package addresses this issue by using a neural network algorithm to differentiate between actual and background RNA expression distributions, effectively eliminating the background noise from genuine expression profiles^65^. For each sample, CellBender was employed with specific parameters (--fpr 0.01 and --epochs 150) to enhance cell clustering and increase the specificity of cell markers. The corrected outputs were subsequently analyzed using Seurat package (v5.0.2)^66^. Although the DecontX and SoupX packages were also tested for removing ambient RNA (data not included), CellBender proved to be the most effective method in our study for removal of the ambient RNA.

### Doublet detection and removal

DoubletFinder uses the principal component matrix to integrate artificial doublets by averaging transcriptional profiles from randomly selected cells and uses the proximity of real expression of a cell in gene expression space to identify real doublets^67^. DoubletFinder function is based on these parameters, (i) the number of artificial doublets (pN), (ii) the neighborhood size to compute the number of artificial nearest neighbors (pK) and (iii) the expected number of doublets(nEXP) as recommended by the 10X protocols. It determines the optimal pK value for each sample by using the Ground-truth doublets function and identifies real doublets. Each sample was processed individually with DoubletFinder, and an nEXP value (3% for 6,000 nuclei) for 1and 8.5-months old samples and nEXP value (5% for 10,000 nuclei) for 17 months old samples were used. Only nuclei confirmed as “singlets” were used for downstream analysis.

### Quality controls and normalization

To maintain high data quality, nuclei expressing fewer than 300 genes or those with more than 3500 genes were excluded. Genes expressing fewer than 5 nuclei were also excluded from downstream analysis. Mitochondrial contamination was filtered by removing nuclei with more than 10 percent of mitochondrial content. After QC processing, data was normalized (normalization.method = “LogNormalize”) by the gene expression of each cell with the total expression and multiplied by a default scaling factor (scale.factor = 10,000) through the Seurat R package. The normalized data were scaled using the default parameters of ScaleData function from Seurat. The scaled expression matrix for Principal component analysis (PCA) in Seurat was used to compute the top 30 significant principal components.

### Integration, clustering and visualization

To reduce batch correction; Integration of the data was performed through IntegrateLayers function in Seurat by using ‘method = HarmonyIntegration’. We further employed elbow method (PCElbowPlot) to select the top 30 significant principal components (PCs) as an input for downstream analysis. For cell type specific sub-clustering and analysis, top 15 principal components were selected as an input for downstream analysis. Transcriptionally similar clusters from similar single nuclei were generated by using K-nearest neighbor (K-NN) graph followed by Louvain community detection algorithm from the previously selected PCs through Graph based clustering algorithm from Seurat^68^. Clustering was optimized by varying PCs (10-30) and resolution parameters (0.2, 0.6, 1.0 and 1.2). All these varying parameters were analyzed for cell types and the final parameter (PCs25, resolution 0.6) was selected on the basis of the presence of the largest number of cell types. For visualization, uniform manifold approximation and projection (UMAP) dimensionality reduction method was employed by using the RunUMAP function in Seurat by using the top selected PCs as the input.

### Cell type identification

Unsupervised clustering analysis revealed 8 different categories of cell identities. Cell identities were determined by the expression of canonical marker genes. For Inhibitory neurons (*Gad1*, *Gad2*), excitatory neurons (*Camk2a, Slc17a7*), Microglia (*C1qa, Arhgap15*), astrocytes (*Aqp4, Gja1*), Oligodendrocyte (*Mbp, Cnp*), OPC (*C1ql1, Pdgfra*) and VLMC (*Slc6a13*) marker genes were used^69,70^. Cell identities were further confirmed through the Allen Cell Type Database (MapMyCells) nomenclature by using hierarchical mapping^71^.

### Subclustering and subtypes identification

To identify subtypes, analysis stated in the “Normalization, clustering and visualization” section were performed to subcluster individual cell types. For excitatory and inhibitory neurons, subcluster were mapped to Allen Cell Type Database (MapMyCells) database and later manually annotated based on the subtype nomenclature^71^. Any cluster having ambiguous identification were removed from downstream analysis. For microglia, astrocyte and oligodendrocytes nuclei were subsets into the individual datasets and subclustered as stated in the “Normalization, clustering and visualization”. In the microglial dataset, the Seurat AddModuleScore function was used to identify NG and DAM score clusters through the gene list **(Extended Data Table.13)**^30^ and visualized by the FeaturePlot_scCustom function from the R package scCustomize(v2.1.2)^72^. Similarly, DAA module score from astrocytes were determined through AddModuleScore function. To infer trajectory analysis of the microglia, we used Monocole3 package in R^73^.

### Differential expression and pathway analysis

Differential expression analysis was performed through MAST (with random effect for sample origin and sequencing depth) to identify DEGs and genes were considered significant if the Log2FC was > 0.1 or <-0.1 and the adjusted P value is <0.05^74^. To minimize the batch effect, DEGs were determined from the integrated dataset at each time point.

### Gene set enrichment analysis

The differential expression signatures were tested for Gene set enrichment analysis (GSEA) for gene sets (hallmark and KEGG) and pathways through fgsea (v1.24.0)^75^. Gene sets and pathways were taken from mSigDB resource. Conversion between rat and human annotation along with entrezgene_id/ ensembl_gene_id performed through merge function from msigdbr (v 7.5.1) package. For each cell type, genes were ranked by decreasing order of Log2FC and analysis was performed through fgsea function with score_type = “std”, num_perm = 1000 parameters. Gene sets pathways were considered to be significant if the adjusted p value was <0.1. Gene Ontology analysis performed through enricher function and/or DisGeNET analysis package^76^.

### Overlap of rat and human DEGs

To examine the extent of concordant overlap (defined as genes showing significant differential expression in the same direction in both datasets) between our snRNAseq analysis from NHe6^-/y^ rats and AD, we analyzed a published snRNAseq dataset of 48 ROSAMAP individuals selected according to modified NIA-Reagan criteria and Braak staging. The cohort included 26 individuals with a pathological diagnosis of AD and 22 control individuals^43^. Hippocampal snRNAseq data were extracted from the AD dataset, and analysis were restricted to the six major cell types included in our study. Differential expression analysis was performed through MAST and genes were considered significant if the Log2FC was > 0.1 or <-0.1 and the adjusted P value is <0.05. Rat gene symbols were converted to their human orthologues using msigdbr. For each cell type and direction of regulation (upregulation and downregulation), overlap enrichment of concordant DEGs was assessed using Fisher’s exact test. For each cell type, the odds ratio and significance of the overlap relative to a background gene universe were calculated. P values were adjusted for multiple testing across all cell types and direction of regulation using the Benjamini-Hochberg procedure with an FDR adjusted significance threshold of 0.05.

### Primary hippocampal neuronal culture

Primary hippocampal neuronal culture was prepared as previously described^8^. Briefly, Hippocampi were isolated from P-0/1 rat pups and dissociated with papain dissociation kit (Worthington #LK003150) in bicarbonate buffered Earle’s Balanced Salt Solution (EBSS) at 37°C for 10 minutes. The tissue was then mechanically triturated 15 times using a 1 mL pipette. Cultures were maintained in Neurobasal medium (Invitrogen, #21103049) supplemented with 2% B27 (Invitrogen, #17504044) and 1% GlutaMAX (Invitrogen, #35050061) and maintained until the day of the experiment.

### Mitochondrial Membrane Potential Assessment by TMRM Fluorescence Imaging

Mitochondrial membrane potential (Δψm) was assessed using protocol adopted from previously published protocols^77,78^. Δψm was estimated in primary hippocampal neurons derived from postnatal day 0 (P0) control and NHE6^-/y^ rats. Neurons were plated on glass-bottom 35 mm dishes and maintained under standard culture conditions until day in vitro 15 (DIV-15). Imaging was performed in freshly prepared, filter-sterilized Tyrode’s buffer (145 mM NaCl, 5 mM KCl, 10 mM glucose, 1.5 mM CaCl₂, 1 mM MgCl₂, 10 mM HEPES), warmed to 37°C prior to use. Tetramethylrhodamine methyl ester (TMRM; Image-iT™, I34361, Thermo Fisher Scientific) was used as a potentiometric fluorescent indicator of MMP. A working solution of 20 nM TMRM, a concentration within the non-quench mode range, was freshly prepared in Tyrode’s buffer from a 100 µM DMSO stock. Cells were washed once with warm Tyrode’s buffer, then incubated with 500 µl of 20 nM TMRM for 15 minutes at 37°C before transfer to a pre-equilibrated microscope environmental chamber (37°C, 5% CO₂) for thermal stabilization. Carbonyl cyanide 4-(trifluoromethoxy)phenylhydrazone (FCCP), a protonophore that collapses MMP, was used as a negative control to validate assay sensitivity. FCCP was prepared from a 100 mM DMSO stock to a final working concentration of 2 µM in the imaging dish. Live-cell imaging was performed on a Nikon Eclipse Ti2 AX R confocal microscope equipped with NSPARC. TMRM was excited at 561 nm, and timelapse images were acquired at low laser power at 10-second intervals over 10 minutes. Baseline fluorescence was established prior to FCCP addition, and mean TMRM fluorescence intensity within mitochondrial regions of interest (ROIs) was measured across the time series, with changes relative to baseline serving as the primary endpoint.

### Mitochondria stress test by Seahorse XFe96 Extracellular Flux Assay

Mitochondrial respiratory function was evaluated using the Agilent Seahorse XFe96 Extracellular Flux Analyzer and the XF Cell Mito Stress Test Kit (Agilent 103015-100), following the manufacturer’s protocol with minor modifications^79^. The day prior to the assay, the sensor cartridge was hydrated with 200 µl per well of Agilent Seahorse XF Calibrant Solution (pH 7.4) and incubated overnight at 37°C in a CO_2_ free humidified incubator. Assay medium was prepared fresh on the day of the experiment by supplementing Agilent Seahorse XF Base Medium with 10 mM glucose (XF 1.0 M Glucose Solution, p/n 103577-100), 1 mM pyruvate (XF 100 mM Pyruvate Solution, p/n 103578-100), and 2 mM glutamine (XF 200 mM Glutamine Solution, p/n 103579-100) to a final volume of 50 ml. The assay medium was warmed to 37°C in a water bath prior to use; no pH adjustment was required. Compounds from the XF Cell Mito Stress Test Kit were reconstituted in assay medium at 10× working concentrations immediately before loading: oligomycin (15 µM), carbonyl cyanide 4-(trifluoromethoxy) phenylhydrazone (FCCP; 10 µM), and rotenone/antimycin A (Rot/AA; 5 µM). Compounds were loaded into the designated ports of the hydrated sensor cartridge according to the sequential injection order specified by the Mito Stress Test protocol. On the day of the assay, culture medium was aspirated from the Seahorse XFe96 Cell Culture Microplate, and cells were washed once with 200 µl of warmed assay medium before the addition of 180 µl of fresh assay medium per well. The cell plate was then transferred to a 37°C CO₂-free incubator for 60 minutes to allow equilibration prior to the run. The XFe96 Analyzer was initialized using the Wave software, and the loaded sensor cartridge (without lid) was inserted for calibration, which required approximately 30-60 minutes. Following successful calibration, the cell plate (without lid) was loaded and the assay initiated. The instrument sequentially measured the oxygen consumption rate (OCR) at baseline and following injection of oligomycin, FCCP, and Rot/AA, enabling calculation of basal respiration, ATP-linked respiration, maximal respiratory capacity, spare respiratory capacity, and non-mitochondrial oxygen consumption. Upon assay completion, the sensor cartridge was discarded in biohazard waste. The cell plate was retained for post-assay normalization of OCR values to nuclei count determined by fluorescence imaging.

### Mitochondrial superoxide assessment by MitoSOX fluorescence imaging

Mitochondrial superoxide was estimated in control and NHE6-/y rat primary hippocampal neurons at DIV-21 using MitoSOX Red mitochondrial superoxide indicator (Thermo Fisher Scientific, M36008) and MitoTracker Green FM (Thermo Fisher Scientific, M7514), adapted from previously described methods^80^. On the day of imaging, neurons were washed with pre-warmed phenol red-free Neurobasal medium and incubated with 0.5 µM MitoSOX Red and 100 nM MitoTracker Green for 20 min at 37°C protected from light. Cells were then washed twice with pre-warmed phenol red-free medium and maintained in fresh medium during image acquisition. Live-cell imaging was performed using a Nikon Eclipse Ti2 AX R confocal microscope equipped with NSPARC with a 60×/1.4 NA oil-immersion objective. MitoTracker Green was excited at 488 nm and fluorescence was collected at 500–550 nm, whereas MitoSOX was excited at 561 nm and fluorescence was collected at 570–620 nm. MitoSOX fluorescence was quantified using Fiji/ImageJ following background subtraction. For each neuron, MitoSOX integrated fluorescence was normalized to MitoTracker Green fluorescence to account for differences in mitochondrial content.

### Isolation of Mitochondria

Hippocampal mitochondria isolation protocol was adopted from published protocols^81,82^. Mitochondria were isolated acutely dissected brain tissue on the day of the assay. Following euthanasia, the brain was rapidly extracted and kept on ice. The hippocampus was dissected and transferred to a pre-chilled 1.5 ml Eppendorf tube containing 500 µl of ice-cold mitochondrial isolation buffer (MSHE: 210 mM D-mannitol, 70 mM sucrose, 5 mM HEPES, 1 mM EGTA, pH 7.2 adjusted with KOH) freshly supplemented with 0.5% (w/v) fatty acid-free BSA (MSHE+BSA). Tissue was washed three times with 500 µl MSHE+BSA, then resuspended in 1.5 ml MSHE+BSA and mechanically dissociated with scissors before transfer to a pre-chilled Dounce tissue grinder. Homogenization was performed on ice with 8-12 strokes, taking care to avoid foaming. The homogenate was transferred to a pre-chilled 2 ml Eppendorf tube and centrifuged at 800 × g for 10 minutes at 4°C to pellet nuclei and cellular debris. The resulting supernatant was transferred to a fresh tube and centrifuged at 8,000 × g for 10 minutes at 4°C to pellet mitochondria. The mitochondrial pellet was washed in 500 µl MSHE+BSA and centrifuged at 4,000 × g for 5 minutes at 4°C. The supernatant was discarded and the pellet resuspended in 50 µl MSHE buffer without BSA. An aliquot of 5 µl was diluted into 45 µl of RIPA buffer for protein quantification by bicinchoninic acid (BCA) assay. The remaining mitochondrial suspension was kept on ice until use.

### Assessment of Mitochondrial ETC Function by Seahorse XFe96

Electron Flow Assay was performed using a protocol adopted from Agilent white sheet and a published protocol^83^. 3× mitochondrial assay solution (MAS) stock was prepared as follows: 660 mM D-mannitol, 210 mM sucrose, 30 mM KH₂PO₄, 15 mM MgCl₂, 6 mM HEPES, and 3 mM EGTA, pH adjusted to 7.2 with KOH. A working 1× MAS solution was prepared by diluting 33.3 ml of 3× MAS to 100 ml with ultrapure water, supplemented with 0.2% (w/v) fatty acid-free BSA. For the assay substrate solution (1× MAS+), 1× MAS was supplemented with 10 mM pyruvate (from 100 mM stock), 2 mM malate (from 500 mM stock), and 4 µM FCCP (from a 10 mM substock prepared in 95% ethanol), and warmed to 37°C prior to use. All substrate and inhibitor stocks were prepared in advance, pH-adjusted to 7.2 where applicable, and stored at 4°C until the day of the assay. The day prior to the assay, the Seahorse XFe96 sensor cartridge was hydrated by adding 200 µl of Agilent XF Calibrant Solution (pH 7.4) per well and incubated overnight at 37°C in a CO_2_-free humidified incubator. Compounds targeting individual ETC complexes were prepared in 1× MAS at 10× working concentrations and loaded into the sensor cartridge ports as follows: Port A, rotenone (Complex I inhibitor; 20 µl of 20 µM, final concentration 2 µM); Port B, succinate (Complex II substrate; 22 µl of 100 mM, final concentration 10 mM); Port C, antimycin A (Complex III inhibitor; 24 µl of 40 µM, final concentration 4 µM); Port D, ascorbate and TMRM (N,N,N′,N′-tetramethyl-p-phenylenediamine; Complex IV electron donors; 26 µl of 100 mM ascorbate and 1 mM TMPD, final concentrations 10 mM and 100 µM, respectively). Port volumes were calculated to achieve the indicated final concentrations in a total well volume of approximately 272 µl following sequential injections. Isolated mitochondria were diluted in 1× MAS+ to achieve seeding densities of 2.5 and 5 µg total protein per well. A volume of 25 µl of mitochondrial suspension was dispensed per well into the Seahorse XFe96 Cell Culture Microplate, omitting background control wells. The plate was centrifuged at 2,000 × g for 20 minutes at 4°C using a swinging-bucket rotor to adhere mitochondria to the plate surface. Following centrifugation, 155 µl of pre-warmed (37°C) 1× MAS+ was carefully added to each well, and the plate was incubated at 37°C for 5 minutes prior to the assay. The assay was initiated on the Seahorse XFe96 Analyzer using the Wave software. Following sensor cartridge calibration, the cell plate was loaded and oxygen consumption rate (OCR) measurements were recorded at baseline and after sequential injection of rotenone (Port A), succinate (Port B), antimycin A (Port C), and ascorbate/TMPD (Port D). This sequential inhibition and substrate supplementation protocol enabled functional assessment of Complex I-, II-, and IV-linked respiration in isolated mitochondria. Following completion of the assay, OCR values were normalized to mitochondrial protein content as determined by BCA assay.

### Assessment of Mitochondrial Coupling by Seahorse XFe96 Coupling Assay

Mitochondrial coupling assay was performed using a protocol adopted from Agilent white sheet and a published protocol^83^. The day prior to the assay, the Seahorse XFe96 sensor cartridge was hydrated with 200 µl of Agilent XF Calibrant Solution (pH 7.4) per well and incubated overnight at 37°C in a CO₂-free humidified incubator. A 3× MAS stock buffer (660 mM D-mannitol, 210 mM sucrose, 30 mM KH₂PO₄, 15 mM MgCl₂, 6 mM HEPES, 3 mM EGTA, pH 7.2) was prepared in advance and stored at -20°C. On the day of the assay, 1× MAS was prepared by diluting the 3× stock with ultrapure water and supplementing with 0.2% (w/v) fatty acid-free BSA. Two substrate solutions were prepared in 1× MAS: (i) for Complex II-driven respiration, succinate (10 mM final) and rotenone (2 µM final), prepared from a 2 mM ethanolic sub-stock; and (ii) for Complex I-driven respiration, glutamate (10 mM final) and malate (10 mM final). All solutions were pH-adjusted to 7.2 and stored at 4°C until use. ADP was prepared fresh on the day of the assay. Compounds were prepared in 1× MAS at 10× working concentrations and loaded into the sensor cartridge ports to achieve the following final concentrations in a total well volume of approximately 272 µl: Port A, ADP (4 mM final; to stimulate State 3 respiration); Port B, oligomycin (2.5 µg/ml final; ATP synthase inhibitor); Port C, FCCP (4 µM final; uncoupler); Port D, antimycin A (4 µM final; Complex III inhibitor). Isolated mitochondria were diluted in 1× MAS+ substrate solution to a seeding density of 10 µg protein per well and dispensed at 25 µl per well into the Seahorse XFe96 microplate, omitting background control wells. The plate was centrifuged at 2,000 × g for 20 minutes at 4°C using a swinging-bucket rotor to adhere mitochondria to the plate surface. Uniform adherence was briefly confirmed by light microscopy at 20× magnification. Pre-warmed (37°C) 1× MAS+ substrate solution (155 µl) was then added per well, and the plate was incubated at 37°C for 5 minutes before loading onto the XFe96 Analyzer. Following cartridge calibration, OCR was recorded at baseline and after sequential injection of ADP, oligomycin, FCCP, and antimycin A, enabling calculation of State 3 (ADP-stimulated), State 4o (oligomycin-inhibited), and maximal uncoupled respiration, as well as the respiratory control ratio (RCR). OCR values were normalized to mitochondrial protein content as determined by BCA assay.

### Parkin translocation by Immunocytochemistry

Parkin translocation assay was adopted from published protocols^84^. Primary hippocampal neurons from control and NHE6^-/y^ P0 pups were plated on glass-bottom dishes and maintained until DIV-21. Cells were fixed in 4% paraformaldehyde in PBS for 15 minutes at room temperature, permeabilized with 0.1% Triton X-100 in PBS for 10 minutes, and blocked in 5% normal donkey serum in PBS for 1 hour at room temperature. Neurons were incubated overnight at 4°C with primary antibodies against Parkin (14060-1-AP, 1:250), the mitochondrial marker ATP5B (ab14730, 1:1000) and neuronal dendrite marker MAP2b (ab5392, 1:1000), diluted in blocking solution. Following three washes in PBS, cells were incubated with species-appropriate fluorophore-conjugated secondary antibodies for 1 hour at room temperature in the dark. Nuclei were counterstained with Hoechst. Coverslips were washed and mounted with an aqueous anti-fade mounting medium. Imaging was performed on a Nikon Ti2 laser scanning confocal microscope using a [63×/1.4 NA oil-immersion] objective. Identical acquisition settings were applied across all experimental groups. Colocalization between Parkin and ATP5B signals was quantified using Manders’ overlap coefficient, calculated using the [JACoP plugin in Fiji/ImageJ]. A threshold was applied consistently across all images prior to analysis. A minimum of [30] neurons per group were analyzed across at least three independent biological replicates. Data are expressed as mean ± SEM, and group comparisons were performed using an unpaired Student’s t-test, with statistical significance defined as p < 0.05.

### Cell-cell communication

Cell-to-cell communication was analyzed using the CellChat package (v2.1.2) ^85^. In brief, following the official workflow, we employed the functions “identifyOverExpressedGenes” and “identifyOverExpressedInteractions” with standard parameters. Since a database for the rat genome is unavailable, we utilized mouse “Secreted Signaling” pathways for the analysis. A heatmap “netAnalysis_signalingRole_heatmap” function was used to visualize and compare the expression of the identified signaling pathways.

### Liquid chromatography-tandem mass spectrometry proteomic analysis

Rats were euthanized at 1.5 (control (n=7) and NHE6^-/y^ (n=8)) and 8.5 (control (n=7) and NHE6^-/y^ (n=8)) months, 200 μm thick coronal sections were dissected on pre-chilled brain matrices and hippocampal tissue were isolated and snap-frozen in liquid nitrogen. Liquid chromatography coupled to tandem mass spectrometry was conducted and the analyses were performed as described previously^86^. Briefly, tissues were homogenized in 8 M urea lysis buffer and protein concentration was determined by bicinchoninic acid (BCA) assay (Pierce). Tandem mass tag (TMT) labeling and high pH fractionation were performed and samples were dried using a SpeedVac. MS was conducted with an Orbitrap Eclipse (Thermo Scientific) in positive ion mode using data-dependent acquisition with 3 second top speed cycles. Proteome Discoverer Suite (v.2.5, Thermo Scientific) software was used to analyze raw datasets. MS/MS spectra were searched against the UniProtKB human proteome database (29918 total sequences) and peptide were assembled into proteins after spectral assignment following strict parsimony principles. Downstream analysis was performed in Python (v3.12.5) using matplotlib (3.9.1), SciPy (1.14.0), NumPy (2.0.1), scikit learn (1.5.1), and pandas (2.2.2). Differential abundance was assessed using Welch’s t-test with Benjamini-Hochberg FDR correction on log-transformed data. PCA was used to assess replicate clustering and reproducibility. Volcano plots and GseGO and enricher analysis were performed in R.

### Immunohistochemistry

Immunohistochemistry was performed on rat brain sections. Rats were perfused with PBS and fixed with 4% Paraformaldehyde. Brain sections (40μm) were collected using cryostat and stored at -20℃ until processed for antibody staining. Sections were rehydrated in PBS and blocked with primary blocking buffer at room temperature for 1 hour. Sections were incubated overnight with Primary antibody including mouse anti-GAD1 (CL488-67648, 1:200), rabbit anti-NHE6 (1:500, Covance, housemade,1:500), mouse anti-CaMKII-alpha (MA1048, 1:300), chicken anti-IBA1 (SYSY#234009, 1:500), chicken anti-GFAP (Aveslab-GFAP, 1:500), mouse anti-NeuN (AB77315, 1;500), mouse anti-CD68 (Bio-Rad #MCA341, 1:500), mouse-anti ALDH1L1 (SKU: 75-164-020, 1:300), rabbit anti-Olig2 (ab109186, 1:500) and rabbit anti-pSTING^-S3^^65^ (D1C4T, cell signaling #62912S, 1:200) were diluted in the blocking buffer at 4℃ and then washed three times with PBS for 5 minutes incubation at room temperature. Alexa-flour conjugated secondary antibodies diluted in the blocking buffer were added to the sections for 1 hour at room temperature and then washed three times with the PBS for 5 minutes at room temperature. Sections were imaged using Olympus and/or Nikon Eclipse Ti2 AX R confocal microscope equipped with NSPARC and analyzed through FIJI.

### Image analysis

To access microglial morphology at 1, 8.5 and 17 months; hippocampus were stained for IBA1 antibody. For each condition, 3-5 independent fields from each section and three sections from each animal (3 animals per condition) were imaged. Briefly, The Freehand selection tool in FIJI was used to measure size of the soma. A minimum of 8 randomly selected microglia soma were measured for and analyzed. The fluorescence intensity of CD68 was quantified through FIJI.

To access neuronal loss, hippocampus were stained for NeuN antibody. For each condition, 3-5 independent fields from each section and three sections from each animal (3 animals per condition) were imaged. The number of NeuN positive cells per each image was manually counted through Fiji from CA1 and DG area. Similarly, the number of IBA1, ALDH1L1, GFAP and Olig2 positive cells per image were manually counted through Fiji.

To access pSTING elevation at 1 and 8.5 months; hippocampus were stained for pSTING antibody. For each condition, 3-5 independent fields from each section and three sections from each animal (3 animals per condition) were imaged. The % area coverage of pSTING was quantified through FIJI. To access pSTING co-localization with cell type markers, hippocampus were stained with pSTING, IBA1, NeuN and ALDH1L1 antibody. The number of IBA1-, NeuNand ALDH1L1positive cells with pSTING expression were manually counted through Fiji.

### Western blot

For western blot, Hippocampi from Control and NHE6-/y rat brain were lysed in RIPA lysis buffer containing (50 mM Tris pH 7.4, 1 mM EDTA, 150 mM NaCl, 10% of TritonX-100, 0.1% of sodium dodecyl sulfate, 0.1% of sodium deoxycholate, protease inhibitor cocktail (Sigma #5892970001), phosphatase inhibitor (Sigma #4906837001) and phenylmethylsulfonyl fluoride (Sigma #93482)). Tissue lysates were lysed for 30 min on ice and were centrifuged at 13,200 rpm for 10 min at 4°C.Protein concentration was determined using a BCA assay with the Pierce BCA Kit (Thermo Scientific # 23225). The protein lysates (30ug) were mixed with NuPage 10x sample reducing agent (Invitrogen #NP0009) and NuPage 4× LDS sample buffer (Invitrogen #NP0007) and then boiled at 70 °C for 10minutes. Subsequently, the samples were loaded onto 4–12% SDS-PAGE gels (Novex #NP0321Box) and transferred to a nitrocellulose membrane (Novex #LC2000). The primary antibodies including rabbit anti-STING (D1V5L, cell signaling #50494, 1:500), rabbit antiphospho-IRF3^-Ser396^ (4D4G, cell signaling #4947, 1:2000), rabbit anti-Rig1 (D14G6, cell signaling #3743, 1:1000) and rabbit anti-MBP (MA5-35074, 1:1000) were incubated in blocking buffer (92765001, LI-COR Biosciences) at 4°C for overnight on a shaker. After three washes, the membranes were incubated with IRDye 680W and 800W goat anti-rabbit and anti-mouse secondary antibodies, diluted in the blocking buffer, for 1 hour at room temperature. analyzed using the Li-CoR Odyssey Imaging System. The membranes were analyzed using the Li-CoR Odyssey Imaging System.

### Statistical analysis

Statistical tests for each experiment are provided in the figure legends.

## SUPPLEMENTAL INFORMATION

## Extended Data Figure Legends

**Extended Data Fig. 1:**
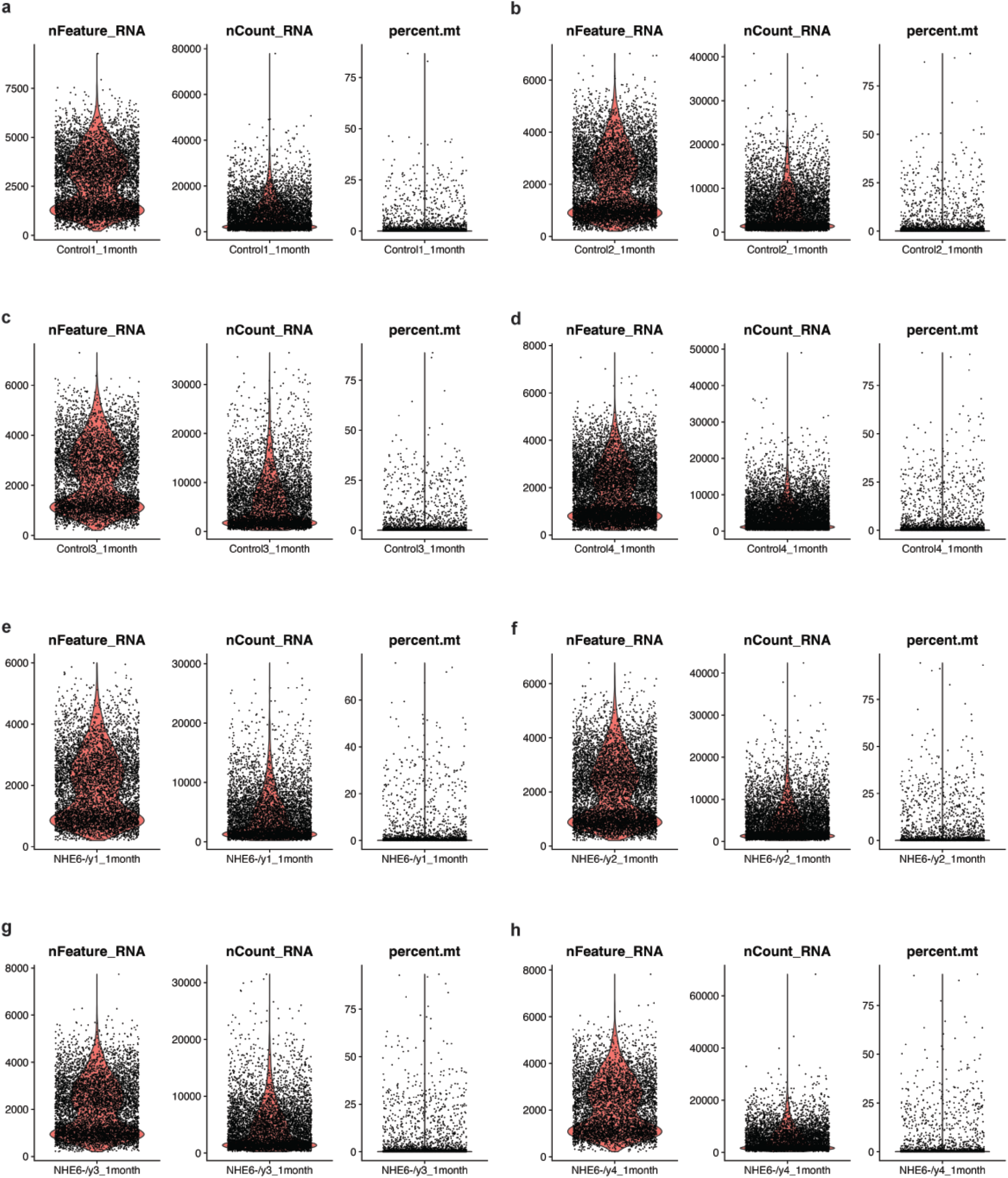
Quality matric calculated by Seurat. **a-h,** snRNAseq profiles of control (n=4) and NHe6-/y rats (n=4) at 1 month. The number of unique genes detected per cell (nFeature_RNA), total number of reads within each cell (nCount_RNA), and percentage of mitochondrial reads (percent.mt) are shown.

**Extended Data Fig. 2:**
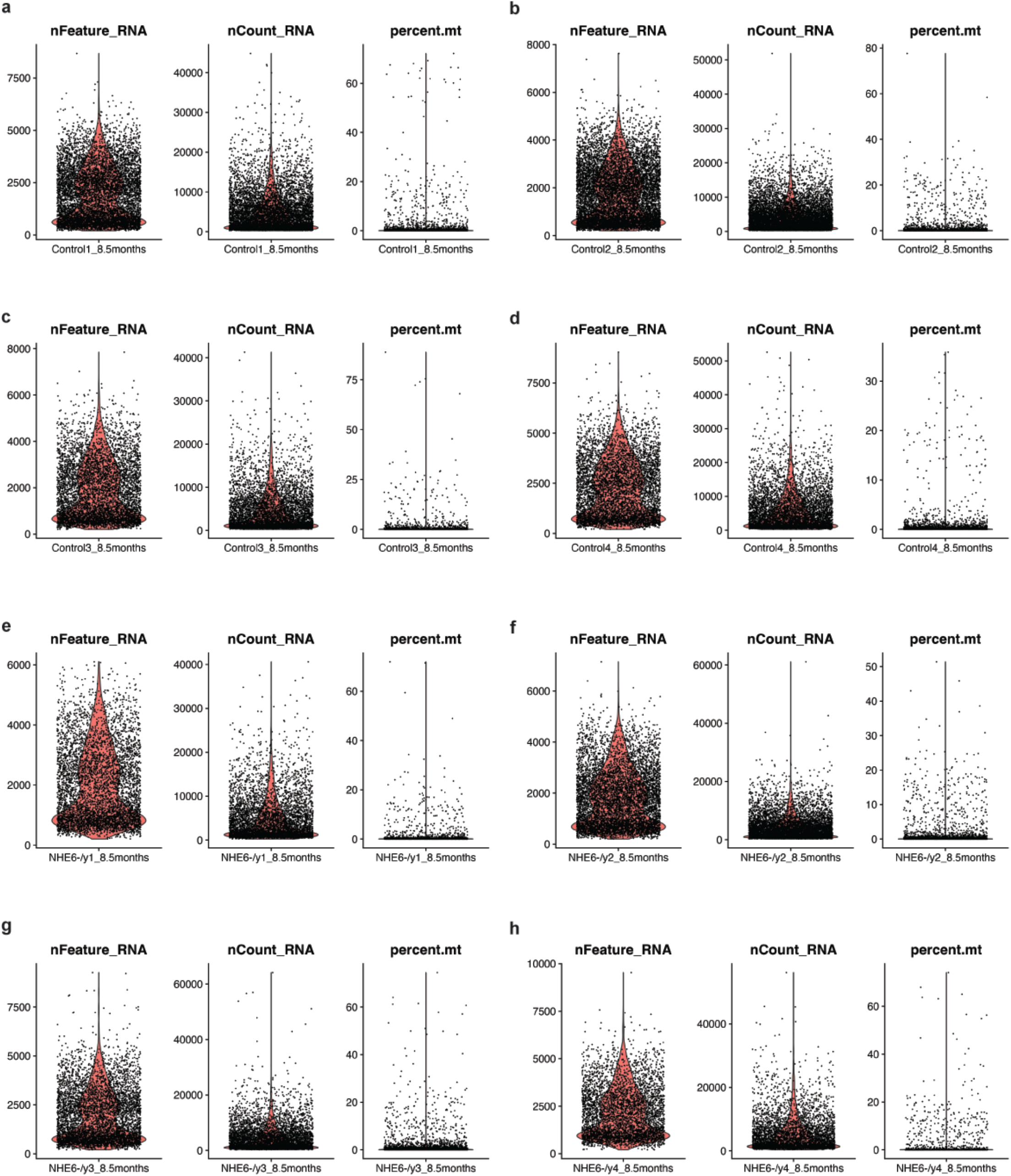
Quality matric calculated by Seurat. **a-h,** snRNAseq profiles of control (n=4) and NHe6-/y rats (n=4) at 8.5 months. The number of unique genes detected per cell (nFeature_RNA), total number of reads within each cell (nCount_RNA), and percentage of mitochondrial reads (percent.mt) are shown.

**Extended Data Fig. 3:**
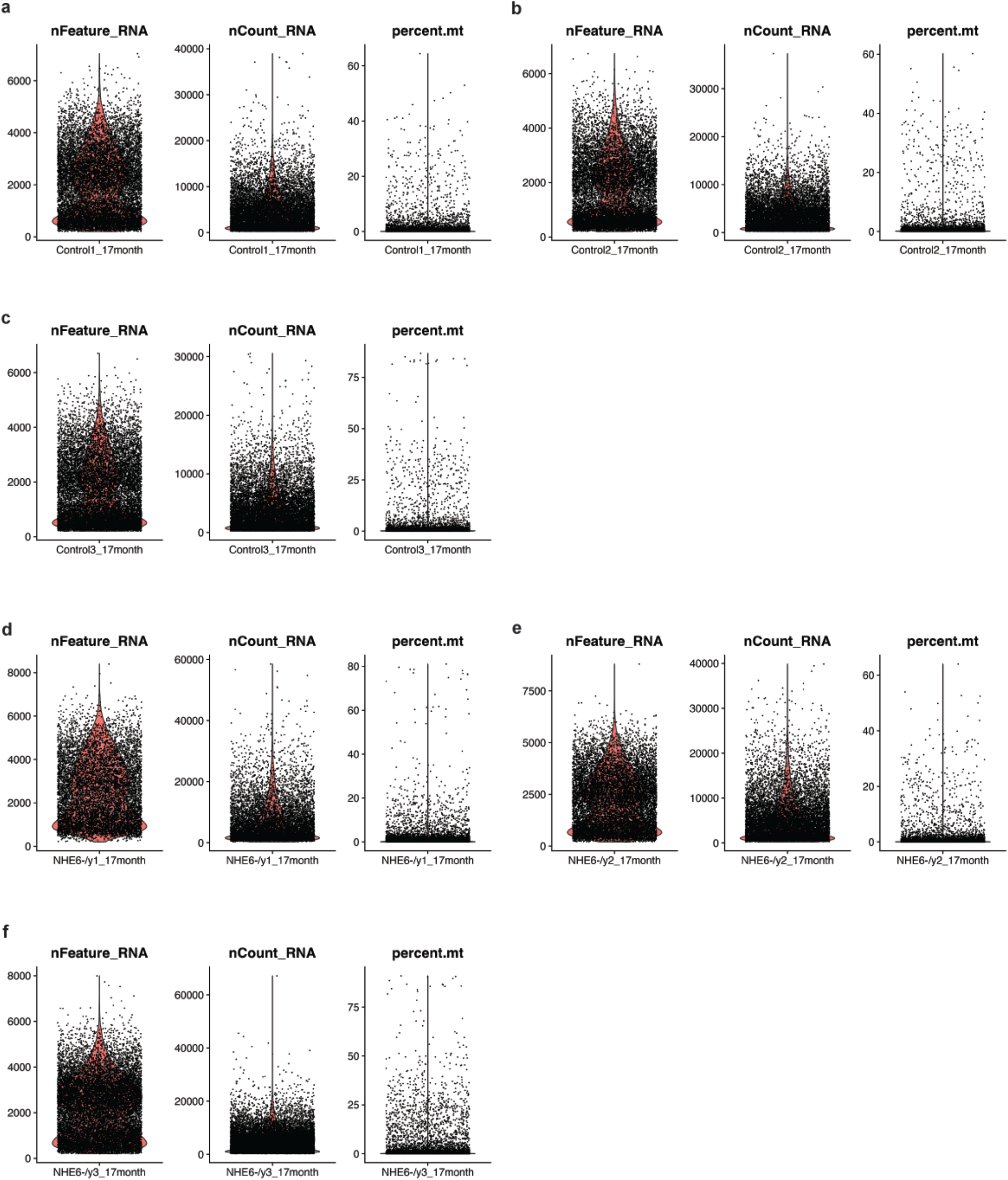
Quality matric calculated by Seurat. **a-f,** snRNAseq profiles of control (n=3) and NHe6-/y rats (n=3) at 17 months. The number of unique genes detected per cell (nFeature_RNA), total number of reads within each cell (nCount_RNA), and percentage of mitochondrial reads (percent.mt) are shown.

**Extended Data Fig. 4:**
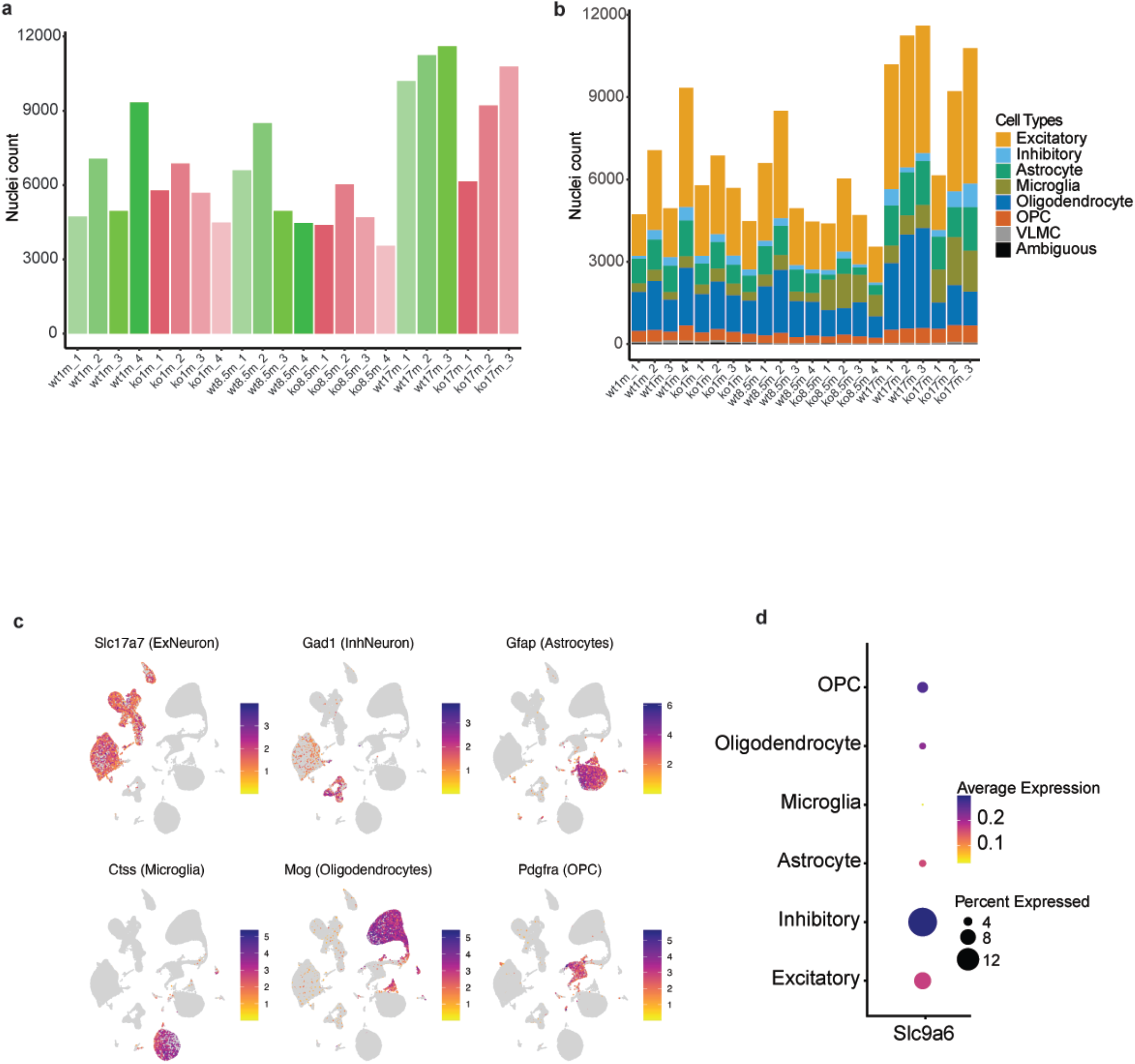
Quality control metrics for single nuclei data. **a,** Total number of nuclei from each sample (all timepoints) in the snRNAseq**. b,** Number of nuclei of each sample in the cell types. **c,** Feature plot showing expression of marker genes used to label main cells: Slc17a7 (excitatory neurons), Gad1 (inhibitory neurons), Gfap1 (astrocytes), Ctss (microglia), Mog (oligodendrocytes) and Pdgfra (OPCs). **d,** Dotplot shows NHE6 expression in individual cell types at 1 month.

**Extended Data Fig. 5:**
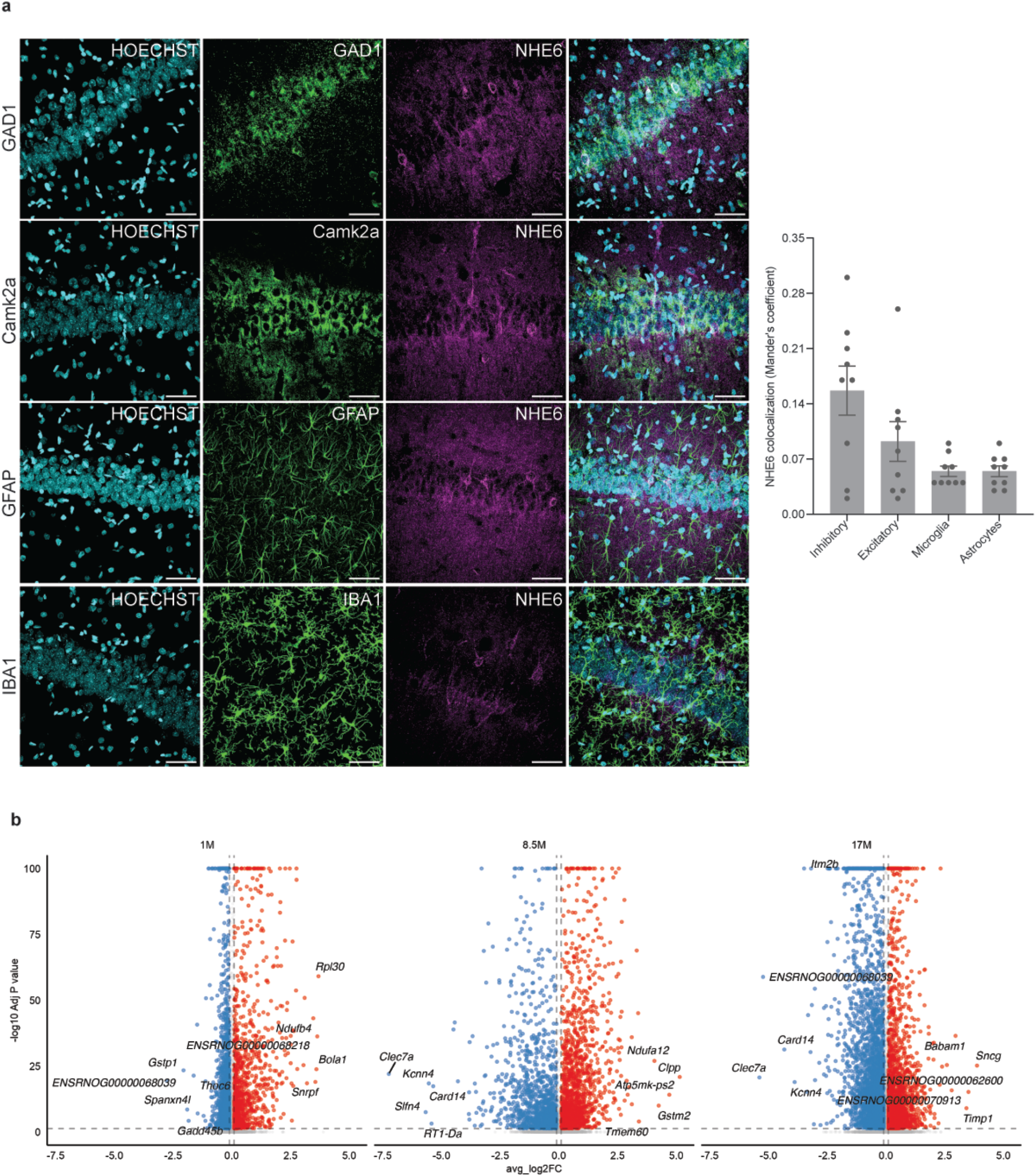
NHE6 expression in cell types. **a,** NHE6 co-localization with marker genes at 1month within major cell types: Gad1 (inhibitory neurons), Camk2a (excitatory neurons), Gfap1 (astrocytes) and IBA1 (microglia). Data are presented as mean ± SEM. Each point in the graphs represents an individual brain section (three sections per rat; Control = 3, NHE6-/y = 3). **b,** Volcano plot shows DEGs between control and NHE6-/y rats nuclei at all time points. Significant genes are labelled in color (Padj < 0.05, Log2FC > 0.1), while nonsignificant genes are shown in gray. The top five upregulated and top five downregulated genes are labeled for each timepoint. Differential expression analysis was performed using MAST, with sample of origin and sequencing depth included as random effects and P values adjusted using the Bonferroni method.

**Extended Data Fig. 6:**
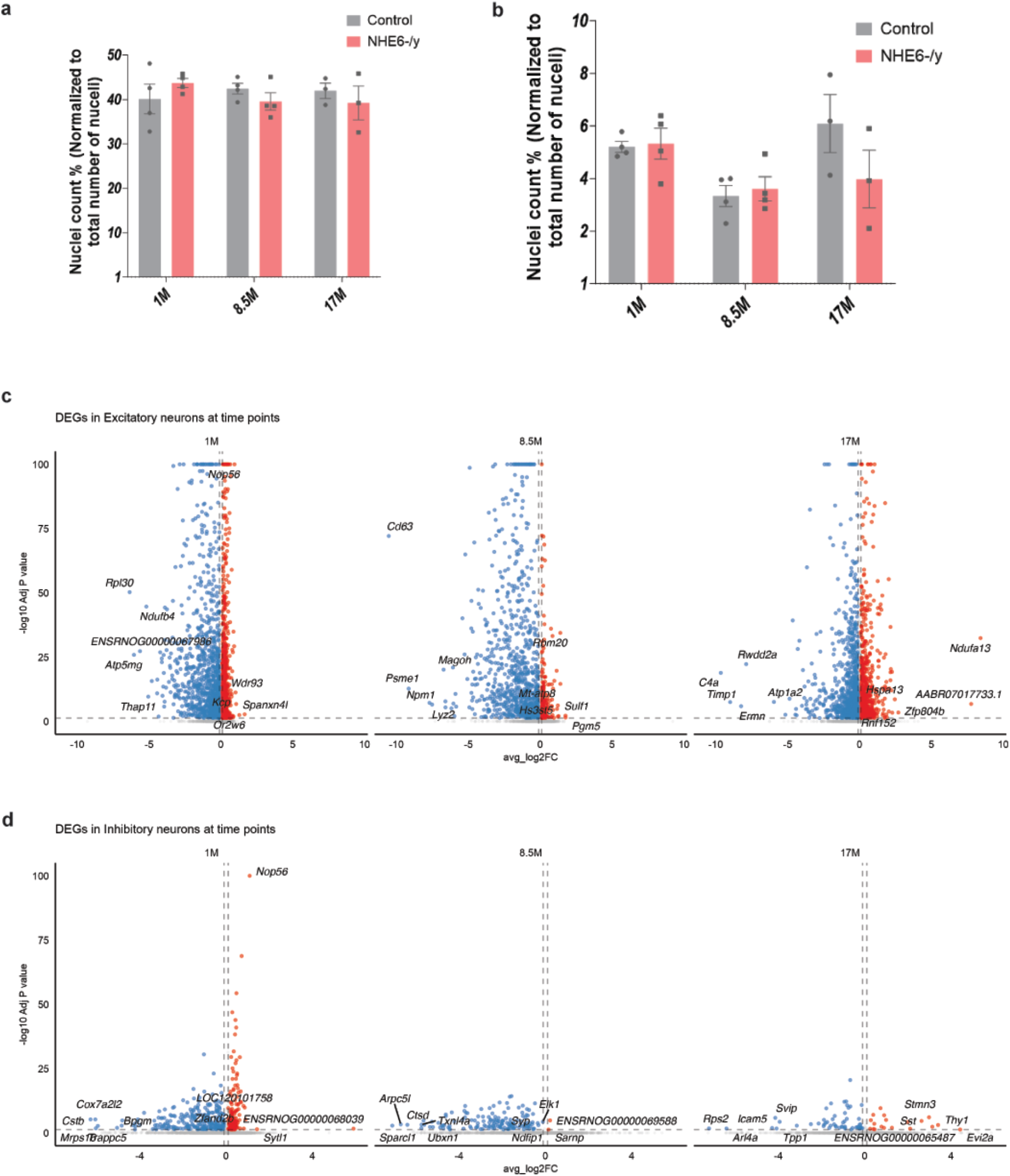
snRNAseq analysis from neurons. **a,** Bar plot indicates no significant change in the number of excitatory nuclei at all timepoints. Data are presented as mean ± SEM. Each point in the graphs represents an individual brain. **b,** Bar plot indicates no significant change in the number of inhibitory nuclei at all timepoints. Data are presented as mean ± SEM. Each point in the graphs represents an individual brain. **c,** Volcano plot shows DEGs between control and NHE6-/y excitatory neurons at all time points. Significant genes are labelled in color (Padj < 0.05, Log2FC > 0.1), while nonsignificant genes are shown in gray. Differential expression analysis was performed using MAST, with sample of origin and sequencing depth included as random effects and P values adjusted using the Bonferroni method. **d,** Volcano plot shows DEGs between control and NHE6-/y inhibitory neurons at all time points. Significant genes are labelled in color (Padj < 0.05, Log2FC > 0.1), while nonsignificant genes are shown in gray. Differential expression analysis was performed using MAST, with sample of origin and sequencing depth included as random effects and P values adjusted using the Bonferroni method.

**Extended Data Fig. 7:**
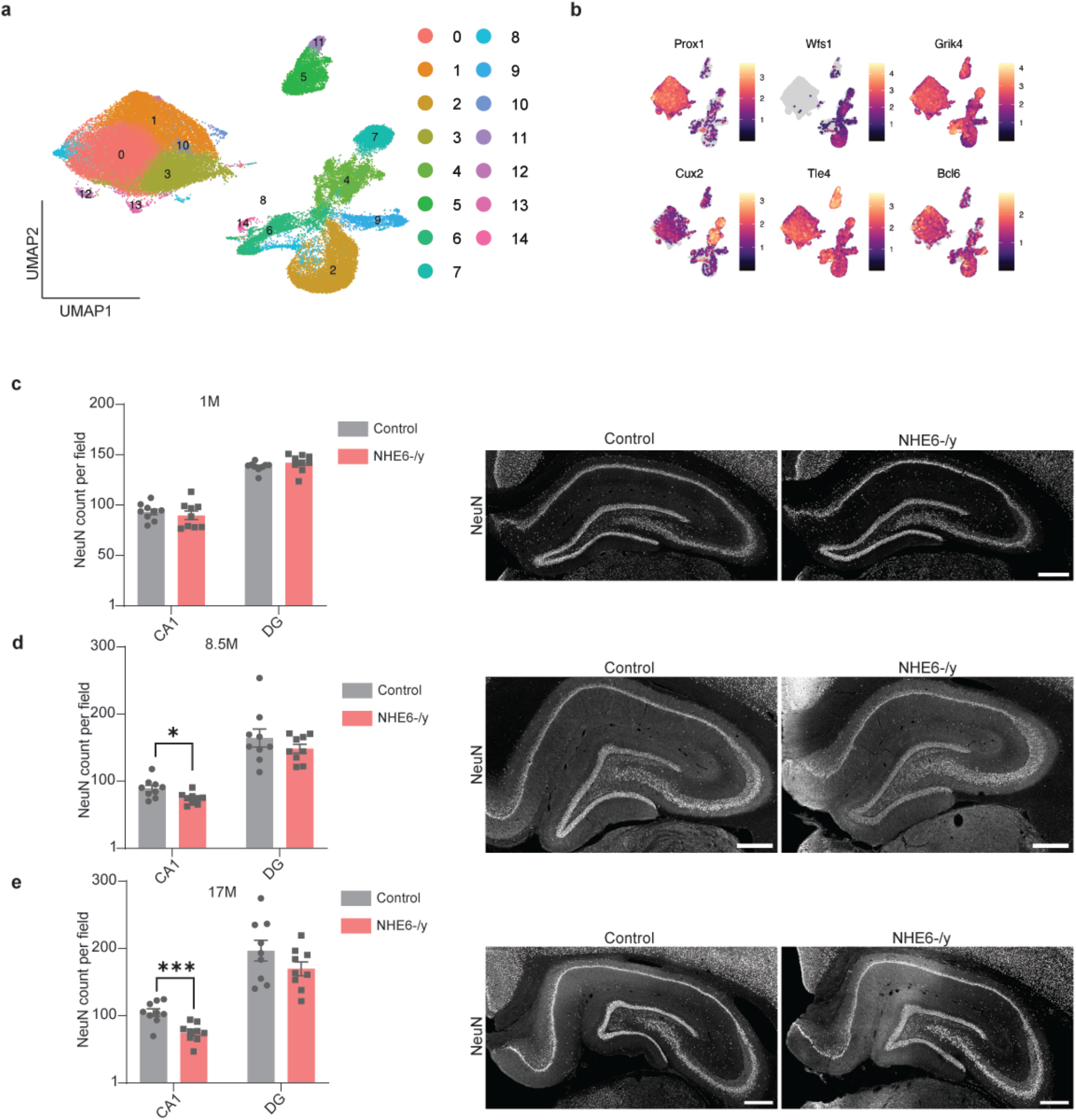
snRNAseq analysis from excitatory neurons. **a,** UMAP plot shows integrated excitatory nuclei clusters at all timepoints. **b,** Featureplot indicates expression of canonical marker genes of excitatory nuclei subtypes. Prox1, DG; Wfs1, CA1; Grik3, CA3; Cux2, L2/3 IT PPP; Tle4, L6b/CT ENT; Bcl6, SUB-ProS. Color scale shows level of gene expression. **c,** Representative images and quantification of number of NeuN positive cells in the CA1 and DG regions at 1 month. Each point in the graphs represents an individual brain section (three sections per rat; Control = 3, NHE6-/y = 3). Data are presented as mean ± SEM. **d,** Representative images and quantification of number of NeuN positive cells in the CA1 and DG regions at 8.5 months. Each point in the graphs represents an individual brain section (three sections per rat; Control = 3, NHE6-/y = 3). Data are presented as mean ± SEM. Two-tailed unpaired t-test with Welch’s correction was performed. *p-value, 0.017. **e,** Representative images and quantification of number of NeuN positive cells in the CA1 and DG regions at 17 months. Each point in the graphs represents an individual brain section (three sections per rat; Control = 3, NHE6-/y = 3). Data are presented as mean ± SEM. Two-tailed unpaired t-test with Welch’s correction was performed. ***p-value, 0.0009. Scale bar 500 µm.

**Extended Data Fig. 8:**
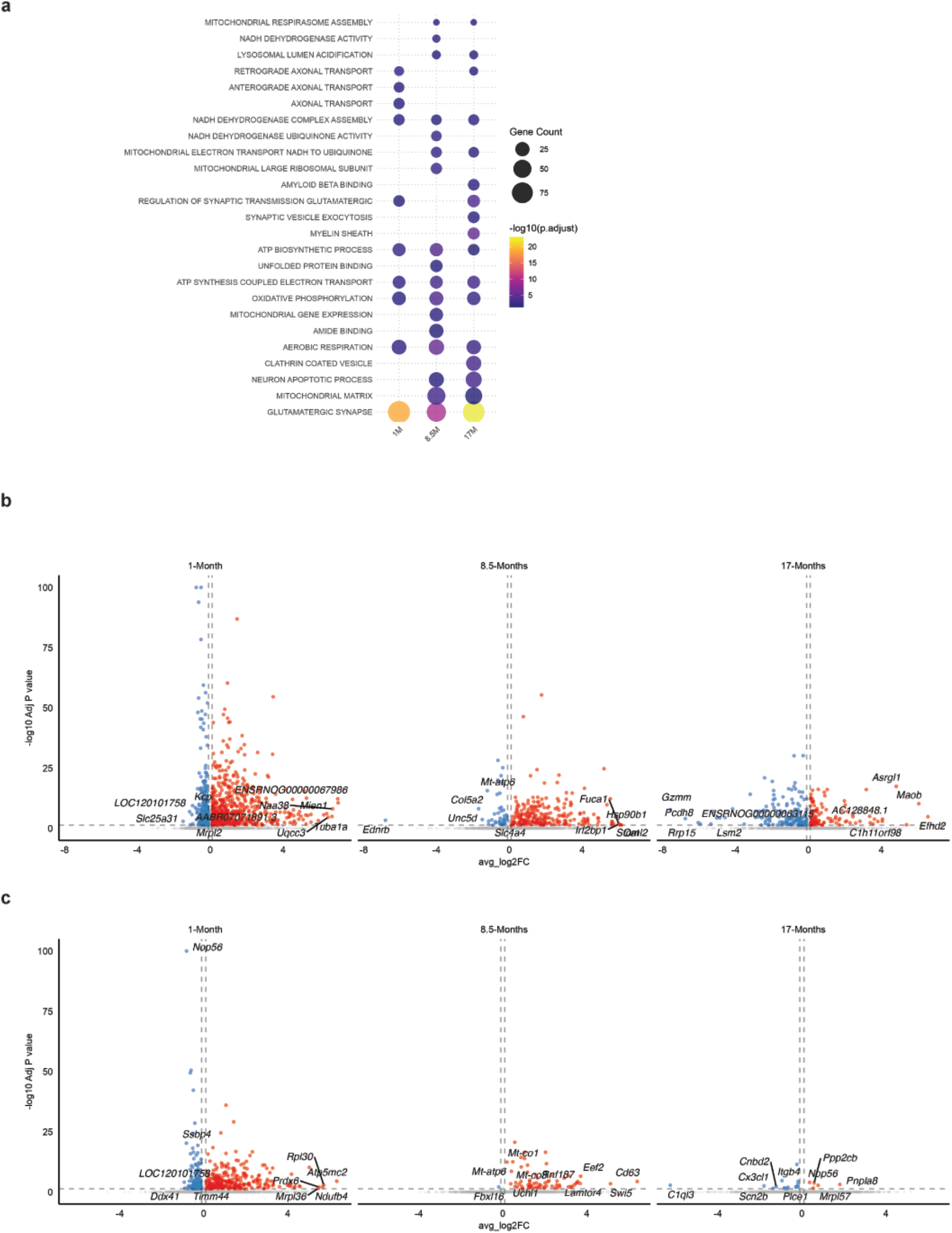
snRNAseq analysis from subtypes of excitatory neurons. **a,** GO analysis from DEGs from DG excitatory neurons. **b,c,** Volcano plots show DEGs from control and NHE6-/y CA1(b) and CA3 (c) nuclei at all time points. Significant genes are labelled in color (Padj < 0.05, Log2FC > 0.1), while nonsignificant genes are shown in gray. Differential expression analysis was performed using MAST, with sample of origin and sequencing depth included as random effects and P values adjusted using the Bonferroni method.

**Extended Data Fig. 9:**
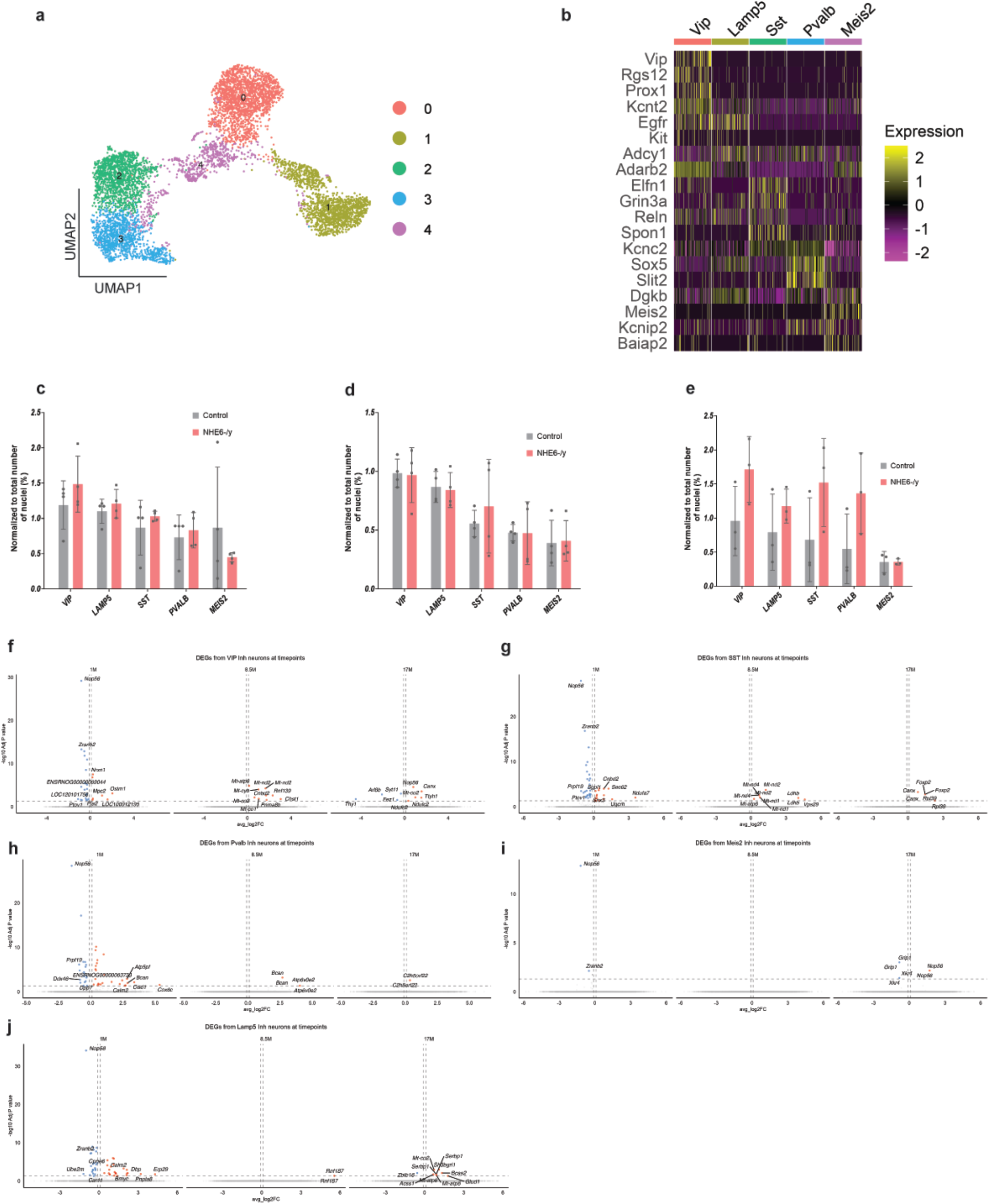
snRNAseq analysis from inhibitory neurons. **a,** UMAP plot shows integrated inhibitory nuclei clusters from all timepoints. **b,** Heatmap shows expression of marker genes of inhibitory nuclei subtypes. Color scale shows level of gene expression. **c-e,** Bar plot indicates no significant change in the number of subtypes of inhibitory nuclei at 1 (c), 8.5 (d) and 17 (e) months. Data are presented as mean ± SEM. Each point in the graphs represents an individual brain. **f-i,** Volcano plots showing DEGs between control and NHE6-/y inhibitory nuclei subclasses VIP (f), SST (g), Pvalb (h), Meis2 (i) and Lamp5 (j) at all time points. Significant genes are labelled in color (Padj < 0.05, Log2FC > 0.1), while nonsignificant genes are shown in gray. Differential expression analysis was performed using MAST, with sample of origin and sequencing depth included as random effects and P values adjusted using the Bonferroni method.

**Extended Data Fig. 10:**
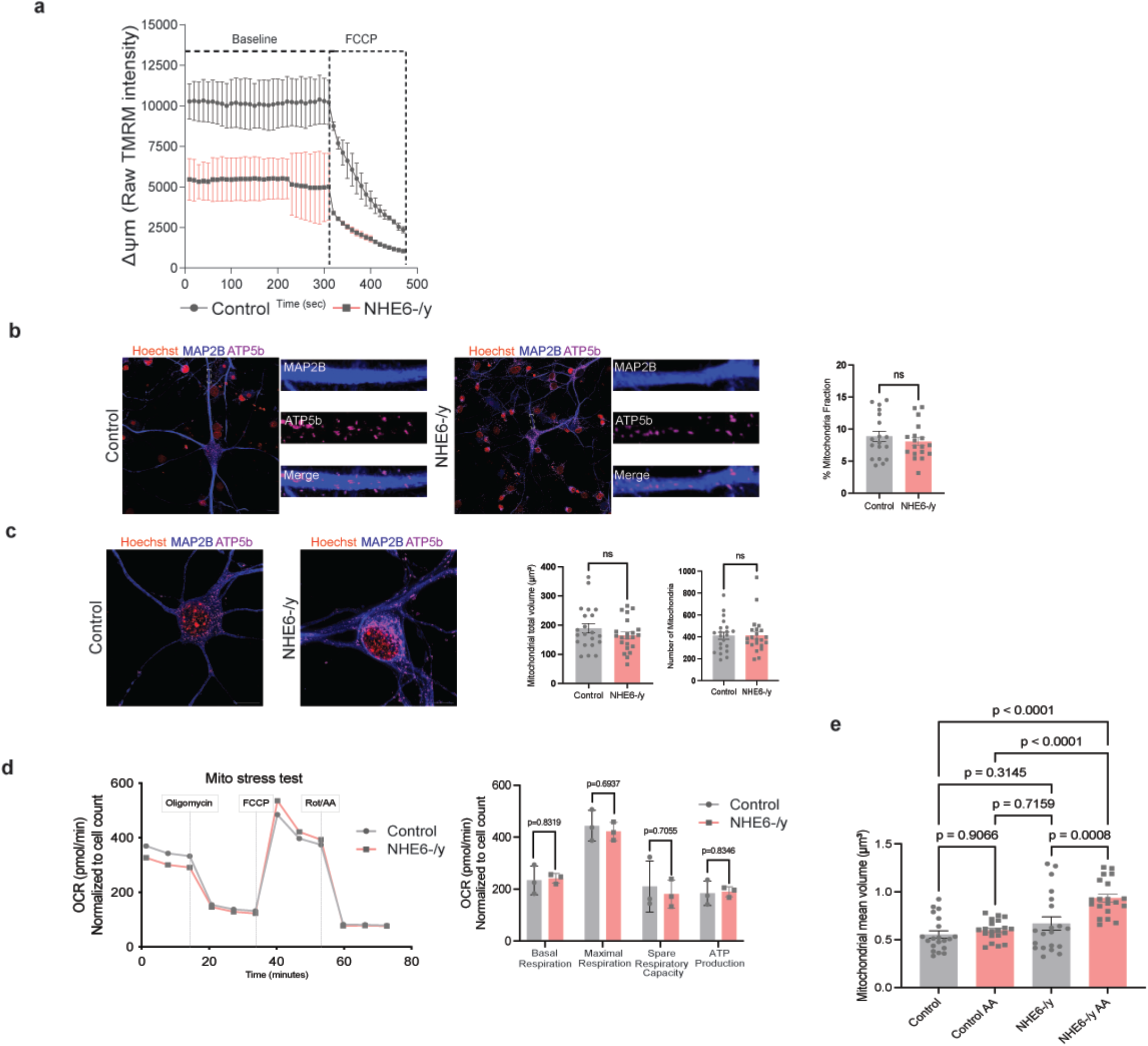
Mitochondrial dysfunction in NHE6-/y neurons. **a,** Time-lapse measurements of raw TMRM fluorescence intensity (Ψm) in control and NHE6-/y neurons. FCCP treatment caused progressive depolarization in both conditions. Data are shown as Mean ± SEM, n = 131 (Control) and 132 (NHE6-/y) neurons from 3 animals per genotype. Two-tailed unpaired t-test with Welch’s correction were performed. **b,** Representative images of DIV-15 Control and NHE6-/y hippocampal neurons immunostained for the dendritic marker MAP2B (blue) and the mitochondrial marker ATP5b (magenta). Insets show magnified views of individual dendrites for each channel and the merged image. Quantification of mitochondrial fraction (% mitochondria fraction) within MAP2B-positive dendrites is shown in the bar graph. Data are presented as mean ± SEM. Each data point represents an individual neuron. Two-tailed unpaired t-test with Welch’s correction was performed. ns, not significant. Scale bar, 10 μm. **c,** Representative images of DIV-15 Control and NHE6-/y hippocampal neurons immunostained for MAP2B (blue), ATP5b (magenta), and Hoechst (Red). Quantification of mitochondrial total volume (μm³) and mitochondrial number per neuron are shown in the bar graphs. Data are presented as mean ± SEM. Each data point represents an individual neuron. Two-tailed unpaired t-test with Welch’s correction was performed. ns, not significant. Scale bar, 10 μm. **d,** Representative Seahorse XF in vitro Mito Stress Test traces show oxygen consumption rate (OCR) in DIV-15 control and NHE6-/y hippocampal neurons. OCR was measured under basal conditions and following sequential injection of oligomycin, FCCP, and rotenone/antimycin A (Rot/AA). Quantification of basal respiration, maximal respiration, spare respiratory capacity and ATP production revealed significantly OCR in NHE6-/y neurons compared to controls. OCR values were normalized to cell count. Data are presented as mean ± SEM. Each data point represents an individual brain (Control = 3, NHE6-/y = 3). Two-tailed unpaired t-test with Welch’s correction was performed. p values are indicated in the graph. **e,** Quantification of mitochondrial mean volume (μm³) in DIV-21 Control and NHE6-/y hippocampal neurons treated with antimycin A (AA) or vehicle. NHE6-/y neurons treated with antimycin A show a significant increase in mitochondrial mean volume compared to all other groups. Data are presented as mean ± SEM. Each data point represents an individual neuron. One-way ANOVA with Tukey’s post hoc test was performed. p values are indicated in the graph. Scalebar 10 μm.

**Extended Data Fig. 11:**
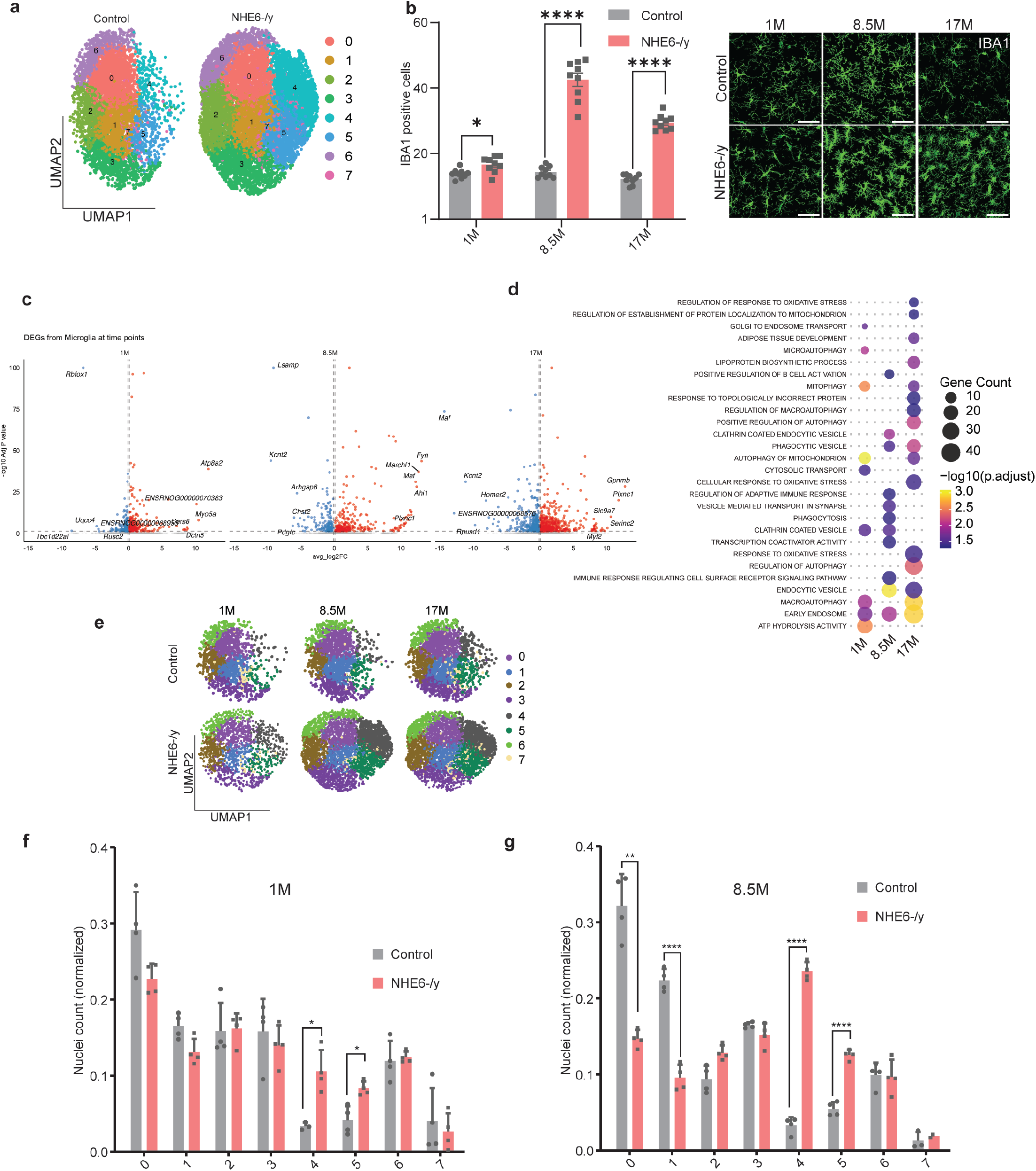
snRNAseq dynamics in microglia. **a,** UMAP plot shows integrated microglial nuclei at all timepoints. **b,** Representative images and quantification of number of IBA1 positive cells at 1, 8.5 and 17 months. Each point in the graphs represents an individual brain section (three sections per rat; Control = 3, NHE6-/y = 3). Data are presented as mean ± SEM. Two-tailed unpaired t-test with Welch’s correction was performed. *p-value, 0.016, ****p-value, p-value<0.000001, ****p-value<0.000001. **c,** Volcano plots show DEGs between control and NHE6-/y rat microglial nuclei at all time points. Significant genes are labelled in color (Padj < 0.05, Log2FC > 0.1), while nonsignificant genes are shown in gray. The top five upregulated and top five downregulated genes are labeled for each timepoint. Differential expression analysis was performed using MAST, with sample of origin and sequencing depth included as random effects and P values adjusted using the Bonferroni method. **d,** Dotplot indicates GO enriched terms between control and NHE6-/y rats at all timepoints. **e,** UMAP plot shows microglial nuclei separated by timepoint and condition. **f,g,** Barplot shows frequency of microglial nuclei in subclusters at 1 and 8.5 months in NHE6-/y rats. Each point in the graphs represents an individual brain. Data are presented as mean ± SEM. Two-tailed unpaired t-test with Welch’s correction was performed. *p-value, 0.012(cluster4-1M), *p-value, 0.013(cluster5-1M), **p-value, 0.002 (cluster0-8.5M), ****p-value, 0.00003 (cluster1-8.5M), ****p-value<0.000001(cluster4-8.5M), ****p-value<0.00002(cluster5-8.5M).

**Extended Data Fig. 12:**
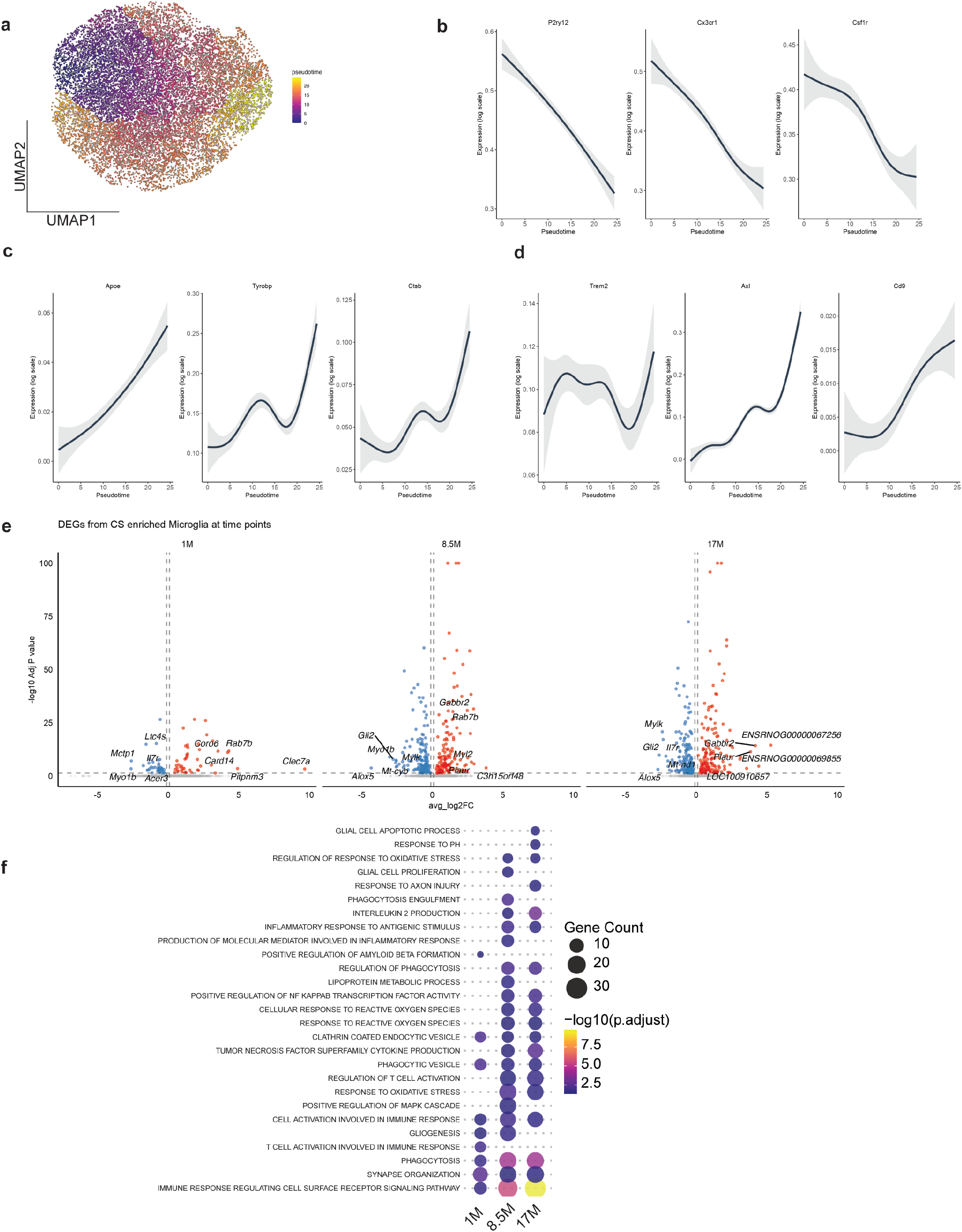
snRNAseq analysis shows increased phagocytic pathways in NHE6-/y rats. **a,** UMAP plot exhibit pseudotemporal ordering of microglia nuclei at all timepoints performed by Monocle3. **b-d,** Gene expression pattern of representative homeostatic (P2ry12, Cx3cr1, Csf1r (b)), DAM1 (Apoe, Tyrobp, Ctsb (c)), and DAM2 (Trem2, Axl, Cd9 (d)) genes along inferred pseudotime, modeled using generalized additive models (GAMs). **e,** Volcano plots indicate DEGs between CS enriched and other microglial nuclei at all time points. Significant genes are labelled in color (Padj < 0.05, Log2FC > 0.1), while nonsignificant genes are shown in gray. The top five upregulated and top five downregulated genes are labeled for each timepoint. Differential expression analysis was performed using MAST, with sample of origin and sequencing depth included as random effects and P values adjusted using the Bonferroni method. **f,** Dotplot indicates significant GO enriched terms between CS enriched and other microglial nuclei.

**Extended Data Fig. 13:**
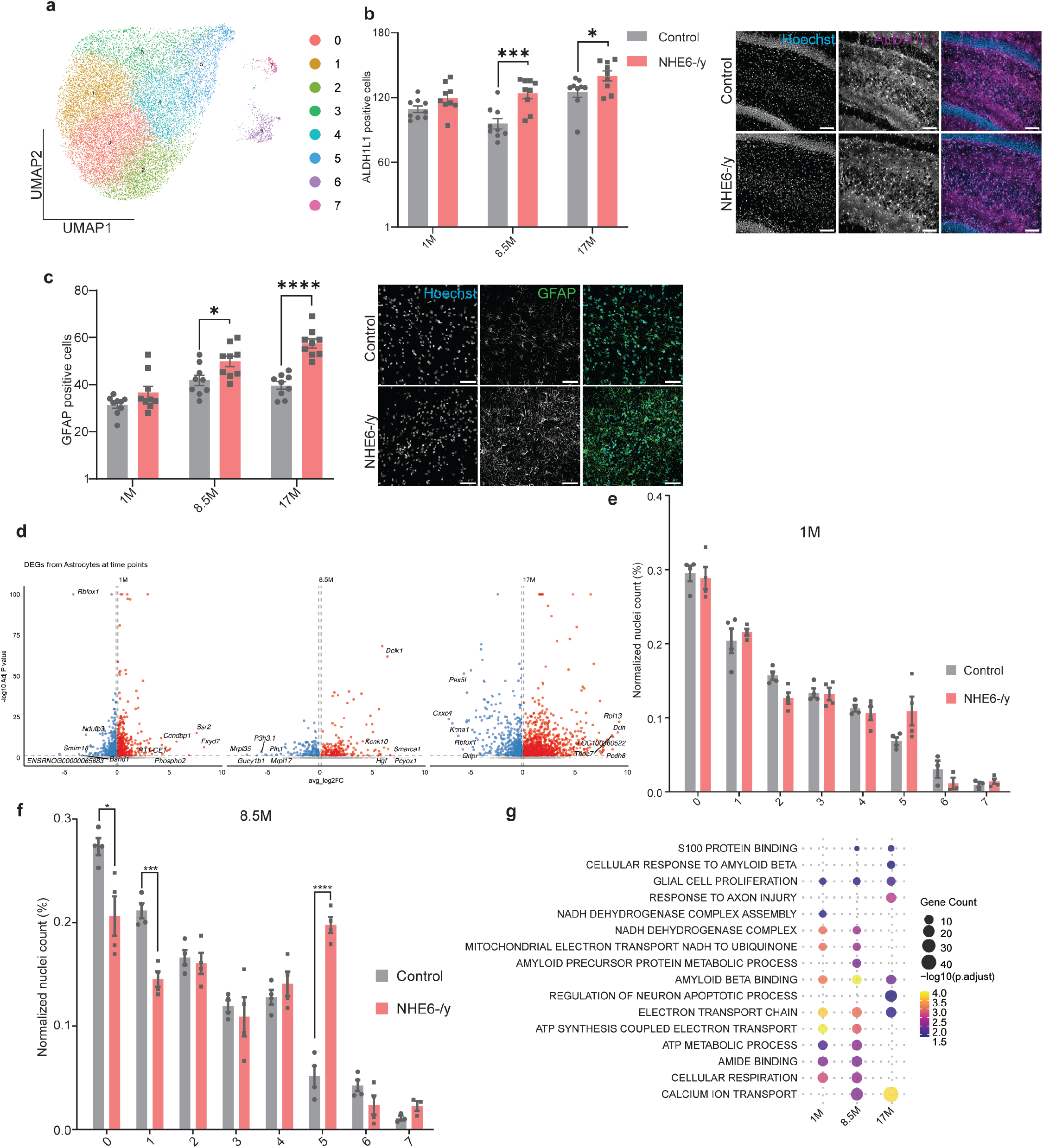
Astrocytes show activated cellular response. **a,** UMAP plot shows integrated astrocytes nuclei at all timepoints. **b,** Representative images and quantification of number of ALDH1L1 positive cells at 1, 8.5 and 17 months. Each point in the graphs represents an individual brain section (three sections per rat; Control = 3, NHE6-/y = 3). Data are presented as mean ± SEM. Two-tailed unpaired t-test with Welch’s correction was performed. *p-value, 0.04, ***p-value, 0.0008. **c,** Representative images and quantification of number of GFAP positive cells at 1, 8.5 and 17 months. Each point in the graphs represents an individual brain section (three sections per rat; Control = 3, NHE6-/y = 3). Data are presented as mean ± SEM. Two-tailed unpaired t-test with Welch’s correction was performed. *p-value, 0.04, *** p-value, 0.0008. **d,** Volcano plots indicate DEGs between control and NHE6-/y astrocytes at all time points. Significant genes are labelled in color (Padj < 0.05, Log2FC > 0.1), while nonsignificant genes are shown in gray. The top five upregulated and top five downregulated genes are labeled for each timepoint. Differential expression analysis was performed using MAST, with sample of origin and sequencing depth included as random effects and P values adjusted using the Bonferroni method. **e,f,** Barplot shows normalized number of astrocytes nuclei at 1 and 8.5 months. Each point in the graphs represents an individual brain (Control = 4, NHE6-/y = 4). Data are presented as mean ± SEM. Two-tailed unpaired t-test with Welch’s correction was performed. *p-value, 0.03, ***p-value, 0.0007, ****p-value, 0.00004.**g,** GO enriched pathways between cluster 5 and other clusters at each timepoints.

**Extended data Fig. 14:**
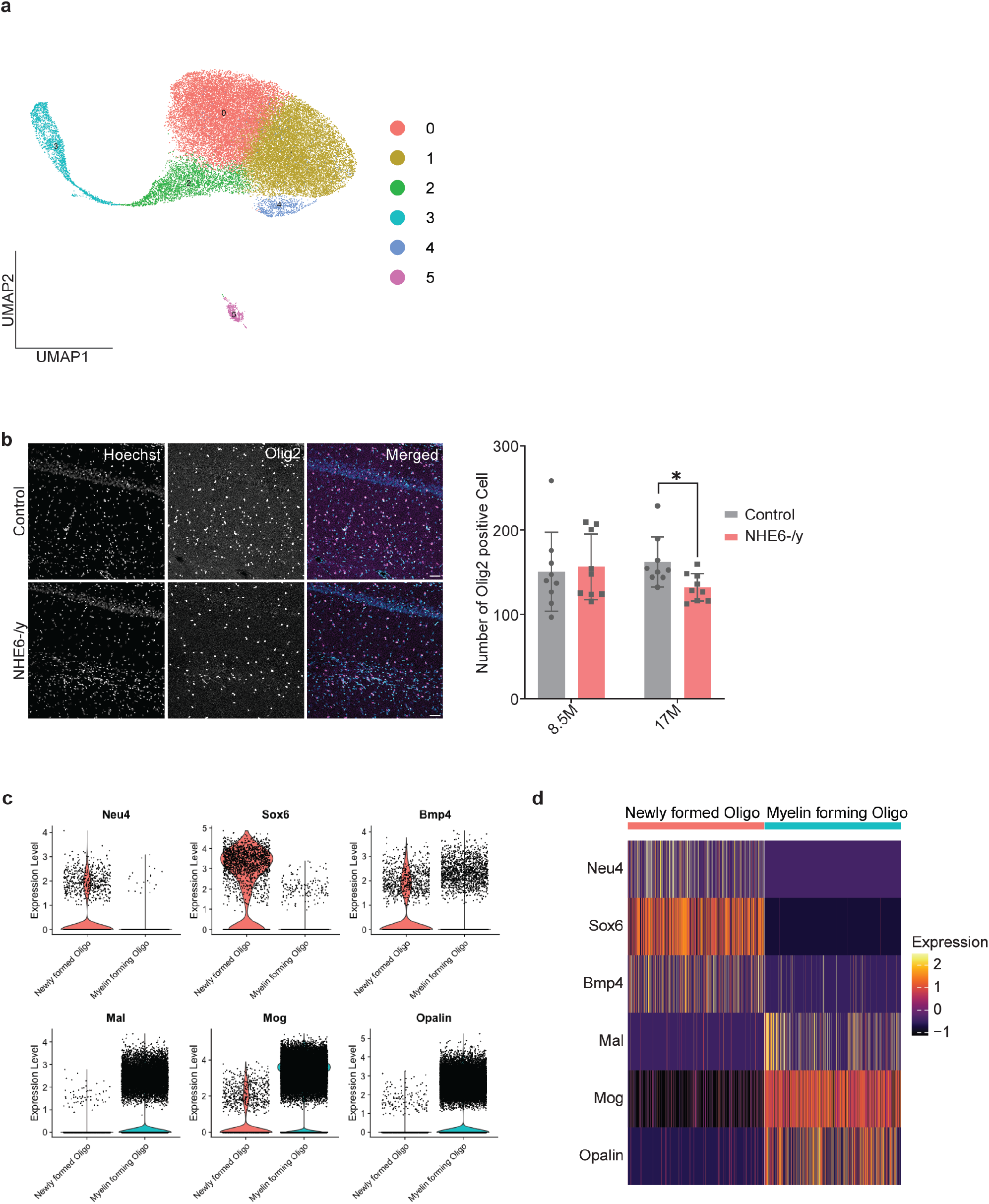
subclassification of Oligodendrocytes. **a,** UMAP plot shows integrated oligodendrocytes nuclei at all timepoints. **b,** Representative images and quantification of number of Olig2 positive cells at 8.5 and 17 months. Each point in the graphs represents an individual brain section (three sections per rat; Control = 3, NHE6-/y = 3). Data are presented as mean ± SEM. Two-tailed unpaired t-test with Welch’s correction was performed. *p-value, 0.02. Scalebar 50 µm. **c,** Violin plots indicate expression of oligodendrocyte marker genes in thesubtypes; Neu4, Sox6 and Bmp4 are marker genes of newly formed oligodendrocytes are enriched in the newly formed oligodendrocytes. Mal, Mog and Opalin are the marker genes of myelin forming oligodendrocytes are enriched in the myelin forming oligodendrocytes. **d,** Heatmap depicts the expression of cell type markers of newly formed oligodendrocytes and myelin forming oligodendrocytes, a maximum of 1000 nuclei per subtypes are displayed.

**Extended data Fig. 15:**
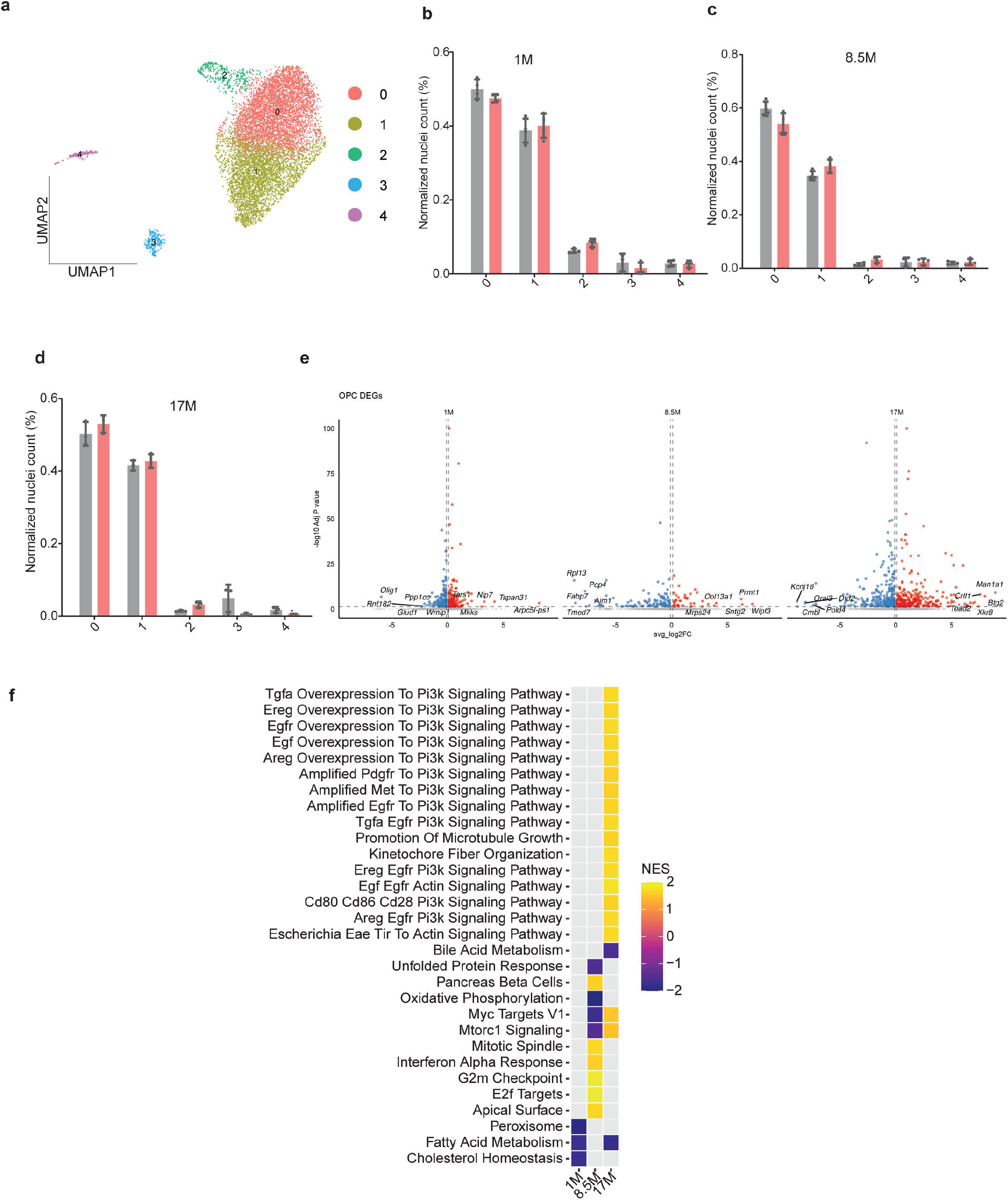
snRNAseq analysis of OPC. **a,** UMAP plot shows integrated OPC nuclei at all timepoints. **b-d,** Barplot shows normalized number of astrocyte nuclei at 1 (b), 8.5 (c) and 17 months (d). Each point in the graphs represents an individual brain. Data are presented as mean ± SEM. **e,** Volcano plots indicate DEGs between control and NHE6-/y astrocytes at all time points. Significant genes are labelled in color (Padj < 0.05, Log2FC > 0.1), while nonsignificant genes are shown in gray. The top five upregulated and top five downregulated genes are labeled for each timepoint. Differential expression analysis was performed using MAST, with sample of origin and sequencing depth included as random effects and P values adjusted using the Bonferroni method. **f,** Heatmap shows GSEA enriched pathways among hallmark and KEGG terms from OPC at all timepoints.

**Extended data Fig. 16:**
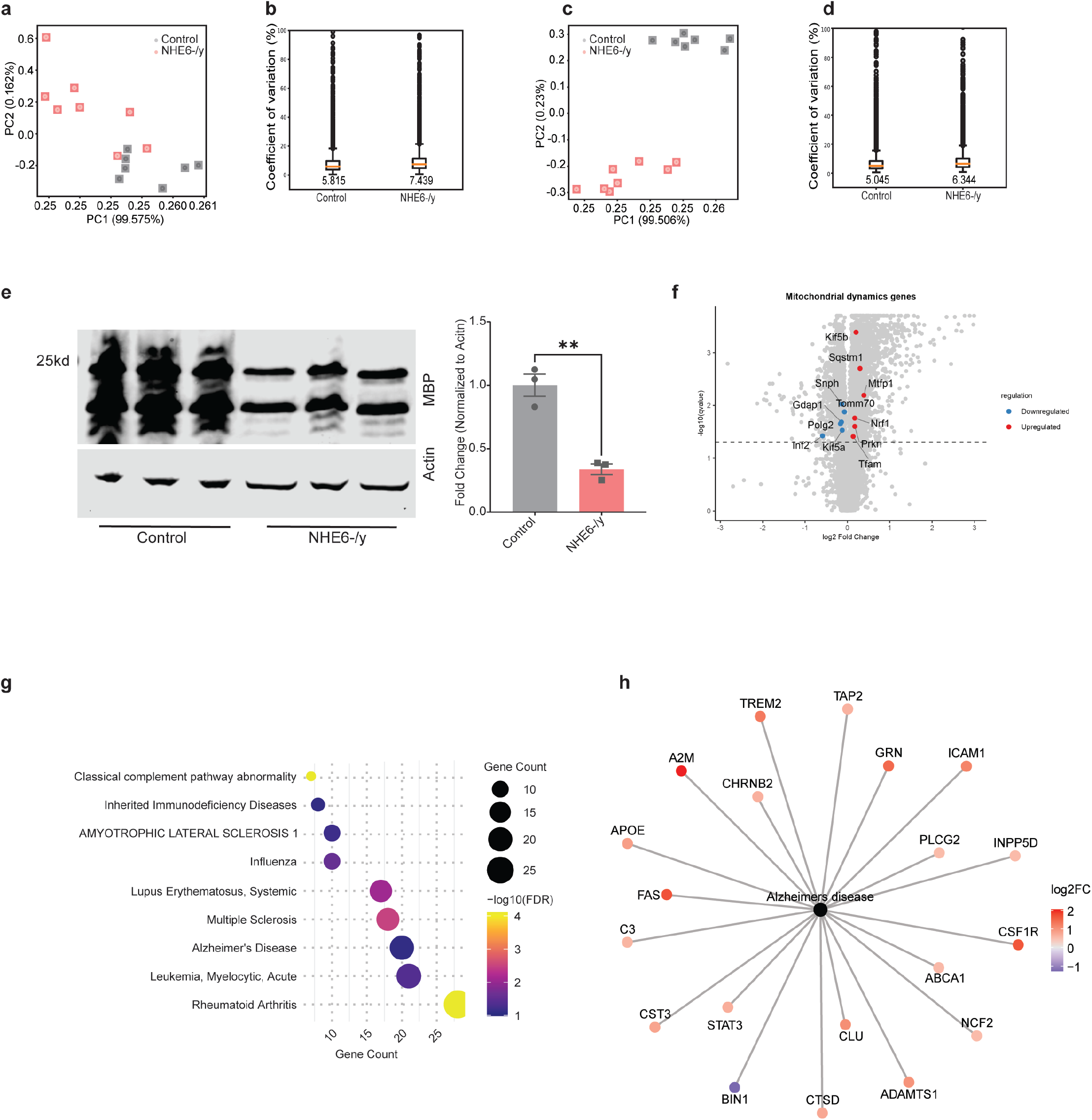
MS analysis from NHE6-/y rats at 15 and 8.5 months. **a,** Principal component analysis (PCA) plot at 1.5 months of Hippocampus MS analysis. Each point in the graphs represents an individual brain (Control = 7, NHE6-/y = 8). **b,** Coefficient of variance (CV) analysis plot at 1.5 months of Hippocampus MS analysis. Each point in the graphs represents an individual brain (Control = 7, NHE6-/y = 8). **c,** PCA plot at 8.5 months of Hippocampus MS analysis. Each point in the graphs represents an individual brain (Control = 7, NHE6-/y = 8). **d,** CV analysis plot at 8.5 months of Hippocampus MS analysis from hippocampus. Each point in the graphs represents an individual brain (Control = 7, NHE6-/y = 8). **e,** Representative western blot and quantification show reduced MBP expression in the hippocampus of NHE6-/y rats at 8.5 months of age. Each point in the graphs represents an individual brain (Control = 3, NHE6-/y = 3). Data are presented as mean ± SEM. Protein levels were normalized to actin. Two-tailed unpaired t-test with Welch’s correction was performed. **p-value, 0.0074. **f,** Volcano plot shows significantly enriched mitochondrial dynamics protein at 8.5 months. Differential abundance was analyzed using Welch’s t-test with Benjamini-Hochberg FDR correction on log-transformed data (q-value < 0.05). **g,** Dot plot shows DisGeNET enrichment analysis performed at 8.5 months. Diseases with FDR (Benjamini-Hochberg adjusted p-value) < 0.1 were considered significantly enriched. Dot size shows the number of input genes associated with each disease (Gene Count), Color represents statistical significance, expressed as -log10(FDR-adjusted p-value). **h,** Network plot shows 20 shared proteins between the CS rat and DisGeNET curated AD associated proteins at the 8.5 months timepoint. Color represents Log2FC in the CS rats dataset.

**Extended data Fig. 17:**
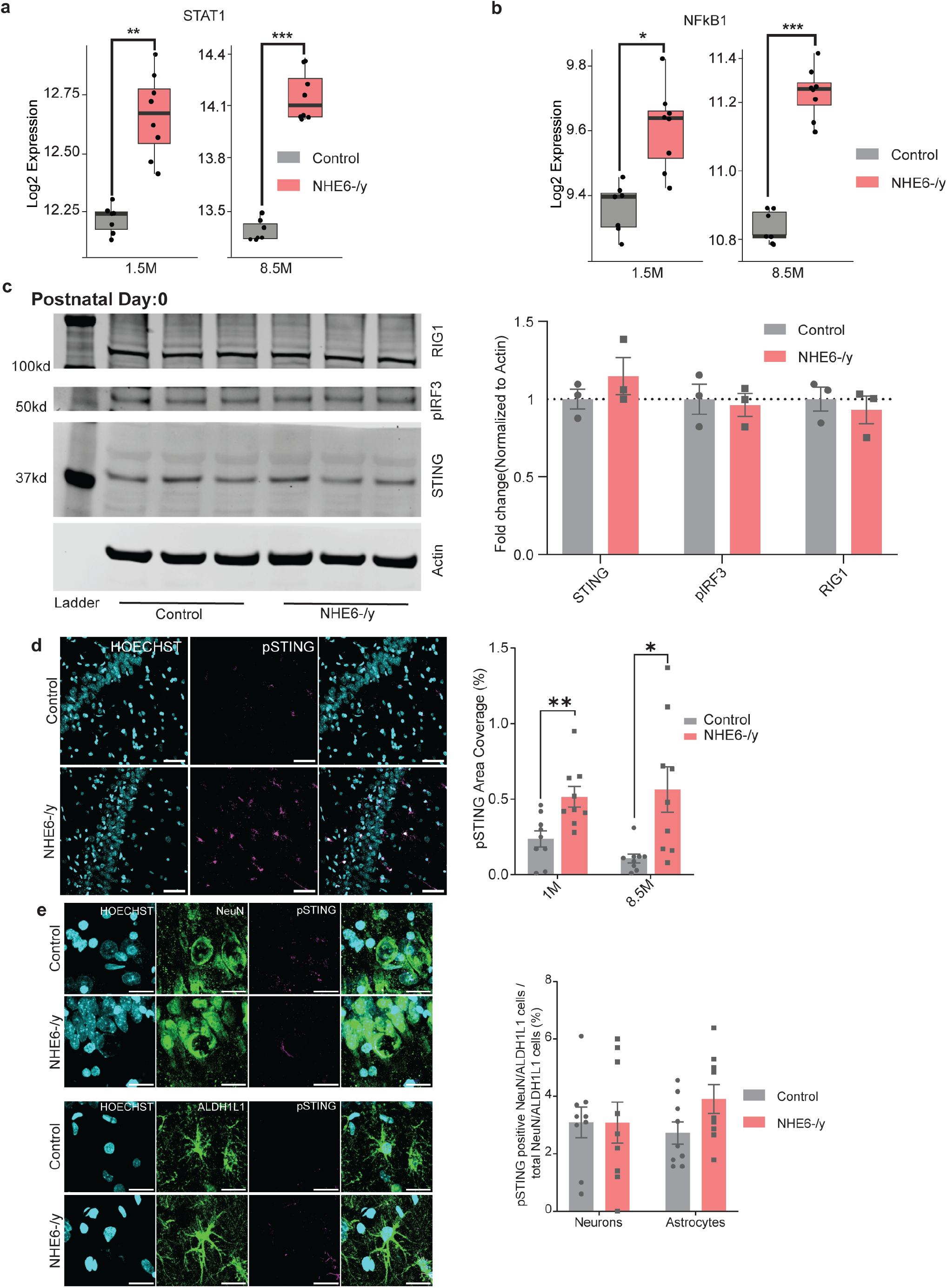
cGAS-STING pathway upregulates in NHE6-/y rats. **a,** Box plots show abundance of STAT1 at 1.5 and 8.5 months. Each point in the graphs represents an individual brain (Control = 7, NHE6-/y = 8). Data are presented as mean ± SEM. Differential abundance was analyzed using Welch’s t-test with Benjamini-Hochberg FDR correction on log-transformed data (q-value < 0.05). **q-value, 0.006, ***q-value, 0.0003. **b,** Box plots show abundance of NFKB1 at 1.5 and 8.5 months. Each point in the graphs represents an individual brain (Control = 7, NHE6-/y = 8). Data are presented as mean ± SEM. Differential abundance was analyzed using Welch’s t-test with Benjamini Hochberg FDR correction on log-transformed data (q-value < 0.05). *q-value, 0.025, ***q-value, 0.0002. **c,** Representative western blot and quantification show no change in STING, pIRF3 and RIG1 expression in the hippocampus of NHE6-/y rats at postnatal day-0. Each point in the graphs represents an individual brain (Control = 3, NHE6-/y = 3). Data are presented as mean ± SEM. Protein levels were normalized to actin. **d,** pSTING (magenta) staining shows elevated expression of pSTING in NHE6-/y rats hippocampus at 1 and 8.5 months. Each point in the graphs represents an individual brain section (three sections per rat; Control = 3, NHE6-/y = 3). Data are presented as mean ± SEM. Two-tailed unpaired t-test with Welch’s correction was performed. **p-value, 0.005, *p-value, 0.016. Scale bar 50 µm. **e,** pSTING colocalization with marker genes NeuN (neurons) and ALDH1L1 (astrocytes) at 8.5months. Each point in the graphs represents an individual brain section (three sections per rat; Control = 3, NHE6-/y = 3). Data are presented as mean ± SEM. Scale bar 20 µm.

**Extended Data Fig.18:**
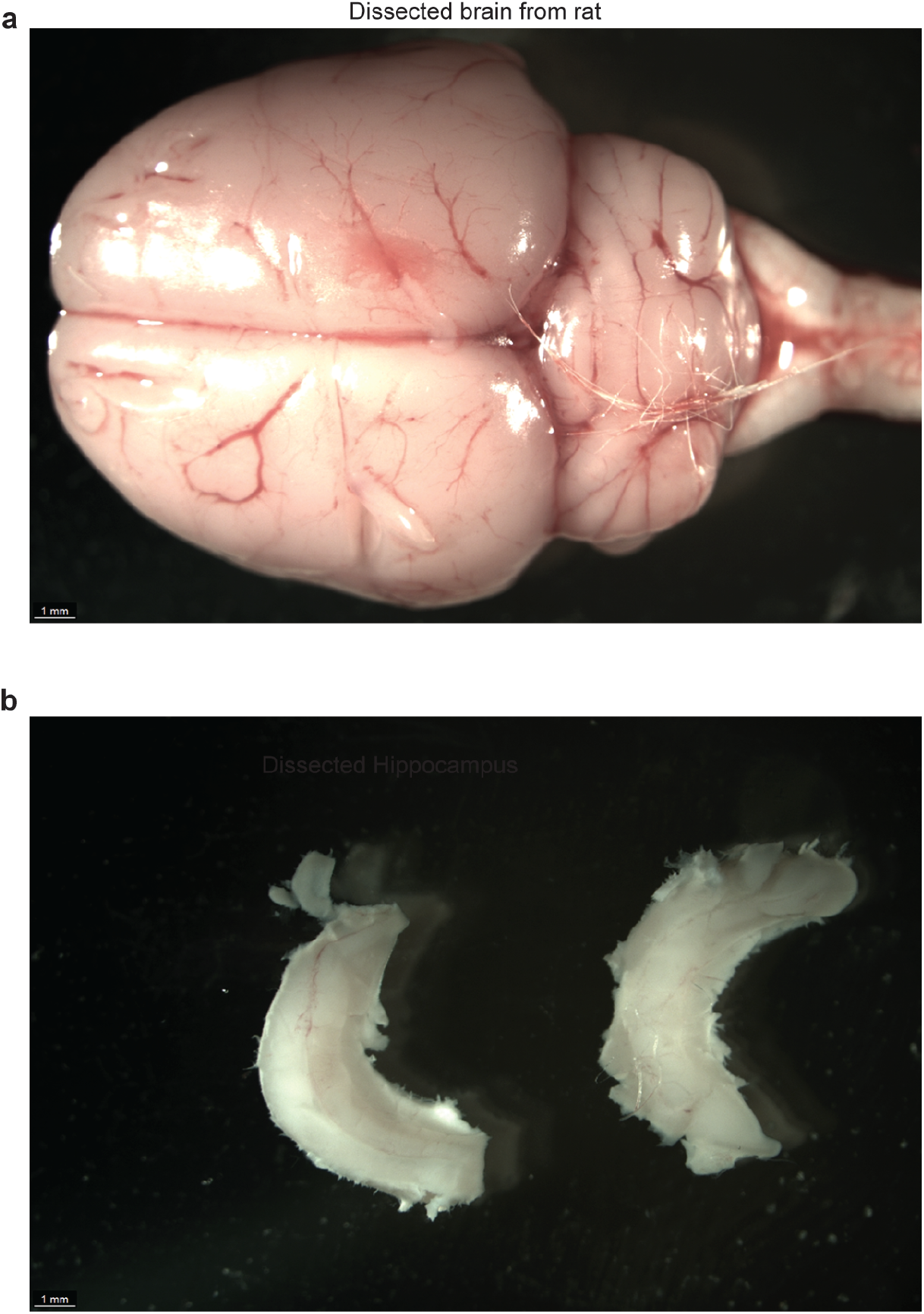
Dissection of hippocampus. **a,** Representative image of dissected rat brain. **b,** Representative image of dissected hippocampus.

## Notes

### Competing Interest Statement

The authors have declared no competing interest.

